# Genomic Touchstone: Benchmarking Genomic Language Models in the Context of the Central Dogma

**DOI:** 10.1101/2025.06.25.661622

**Authors:** Yihui Wang, Zhiyuan Cai, Qian Zeng, Yihang Gao, Jiarui Ouyang, Yingxue Xu, Shu Yang, Sunan He, Yuxiang Nie, Yu Cai, Fengtao Zhou, Cheng Jin, Xi Wang, Zhi Xie, Danqing Zhu, Ting Xie, Kwang-Ting Cheng, Can Yang, Xi Fu, Jiguang Wang, Kang Zhang, Jianhua Yao, Raul Rabadan, Hao Chen

## Abstract

The emergence of genomic language models (gLMs) has revolutionized the analysis of genomic sequences, enabling robust capture of biologically meaningful patterns from DNA sequences for an improved understanding of human genome-wide regulatory programs, variant pathogenicity and therapeutic discovery. Given that DNA serves as the foundational blueprint within the central dogma, the ultimate evaluation of a gLM is its ability to generalize across this entire biological cascade. However, existing evaluations lack this holistic and consistent framework, leaving researchers uncertain about which model best translates sequence understanding into downstream biological prediction. Here we present **Genomic Touchstone**, a comprehensive benchmark designed to evaluate gLMs across 36 diverse tasks and 88 datasets structured along the central dogma’s modalities of DNA, RNA, and protein, encompassing 5.34 billion base pairs of genomic sequences. We evaluate 34 representative models encompassing transformers and convolutional neural networks, as well as emerging efficient architectures such as Hyena and Mamba. Our analysis yield four key insights. First, gLMs achieve comparable or superior performance on RNA and protein tasks compared to models pretrained on these molecules. Second, transformer-based models continue to lead in overall performance, yet efficient sequence models show promising task-specific capabilities and deserve further exploration. Third, the scaling behavior of gLMs remains incompletely understood. While longer input sequences and more diverse pretraining data consistently improve performance, increases in model size do not always translate into better results. Fourth, pretraining strategies, including the choice of training objectives and the composition of pretraining corpora, exert substantial influence on downstream generalization across different genomic contexts. Genomic Touchstone establishes a unified evaluation framework tailored to human genomics. By spanning multiple molecular modalities and biological tasks, it offers a valuable foundation for guiding future gLM design and understanding model generalizability in complex biological contexts.

## Introduction

Large language models (LLMs) have revolutionized natural language processing (NLP) by leveraging large-scale pretraining on diverse textual data to capture complex syntactic and semantic patterns^1,2^. Inspired by the similarity between genome’s biological code and human language^3^, as well as the abundance of omics data driven by advances in high-throughput sequencing technologies, recent studies have extended LLM frameworks to biological sequences, which, like language, encode hierarchical and context-dependent information. Notably, the success of AlphaFold^4^ in predicting protein structures and models such as ESM^5^ and DNABERT^6^ has demonstrated that language models can uncover meaningful patterns in biological sequences, thereby opening new avenues for computational biology and genomics^3^.

In the evolution of genomic language models (gLMs), researchers have explored a range of design choices to better capture the statistical and functional properties of DNA sequences. Early models focused on adapting successful architectures from NLP, such as BERT-style transformers^6^, to represent genomic sequences. As the field progressed, later models began to address domain-specific challenges including efficient long-range dependency modeling, scalability to long input sequences, and cross-species generalization^7–11^. This led to the emergence of a diverse ecosystem of models that differ in several key dimensions: model architectures, which vary in how they capture sequence dependencies and scale with input length^7,10,11^; pretraining strategies, with different training data scopes and learning objectives^8,12^; and evaluation protocols, which impact how effectively models are compared in terms of downstream performance^8,13,14^. Despite their diverse architectures and pretraining, the vast majority of gLMs are fundamentally pretrained on DNA sequences. This foundation in DNA sequences perfectly aligns with its fundamental role as the starting point of the central dogma^15^—the process governing the flow of genetic information to RNA and protein. As the primary information carrier, DNA encodes the foundational instructions for all cellular functions. This creates a natural predictive hierarchy and thus, the ultimate evaluation for a model trained on the language of the genome is not just its ability to understand patterns within DNA, but its power to generalize and predict functional consequences across the entire biological cascade. However, current practices fall far short of this ultimate evaluation of providing a holistic, biologically-grounded assessment, and often even lack the basic standardization required for consistent comparison. Many existing benchmarks focus on a limited set of tasks, primarily classification problems, which are often constructed in ways that deviate from real-world biological distributions or lack relevance to broader genomic contexts^13^. For example, datasets may be artificially balanced or based on short, isolated sequence segments, which restricts their ability to assess whether models can capture biologically important characteristics such as regulatory signals or long-range dependencies^16,17^. Moreover, current evaluation protocols often lack sufficient discriminative power to differentiate between models with varying architectures or pretraining datasets^8,18^, making it difficult to identify strengths and weaknesses or to inform practical model selection. A further critical limitation is that most benchmarks concentrate solely on DNA-based tasks, thereby failing to apply the ultimate evaluation of a model’s utility: its ability to generalize across the RNA and protein layers as prescribed by the central dogma^6,8,13^. Collectively, these deficiencies have led to an absence of consensus on optimal model selection, often leaving biological researchers uncertain which tool is best suited to address their specific scientific questions. Therefore, a comprehensive and biologically representative framework is urgently needed to serve as a Touchstone, rigorously evaluating whether the diverse and rapidly evolving gLMs are truly capable of addressing real-world biological challenges.

Here, we introduce **Genomic Touchstone** (Fig. 1), a comprehensive benchmarking framework structured around the central dogma to evaluate genomic language models in a consistent, discriminative, and biologically meaningful manner. This benchmarking framework is specifically designed to thoroughly assess the ability of gLMs to model complex genomic data and address real-world biological challenges. Specifically, given the central role of human data in biomedical research, where accurate modeling of gene regulation, variant function, and molecular phenotypes is critical for understanding disease mechanisms, improving diagnostics, and enabling therapeutic development, **Genomic Touchstone** includes **34** widely used human-centric gLMs, with diverse architectures (e.g., CNN, Transformer, Bigbird, Hyena, Mamba), pretraining paradigms, and model sizes ranging from 3.3 million to 2.5 billion parameters. We perform extensive experiments on **88** real-world, human-centric biological datasets, which in total comprise **5.34** billion base pairs of genomic sequences. These datasets are used to integrate **36** tasks spanning multiple length scales and diverse task types, each carefully curated to cover different human genomic regions and biological processes aimed at fostering an improved understanding of human genome-wide regulatory programs, variant pathogenicity and therapeutic discovery. Unlike previous benchmarks that focused exclusively on DNA sequences, Genomic Touchstone is the first to include tasks involving RNA and protein sequences. This expanded scope not only offers a more realistic evaluation of model performance but also rigorously tests their ability to capture the entire spectrum of the central dogma of biology^15^, thereby reflecting the complete flow of genetic information. Moreover, for practitioners, Genomic Touchstone would serve as crucial references to inform decisions regarding model selection for particular genomic applications (Fig. 2). Ultimately, the establishment of Genomic Touchstone is essential not only for effectively applying these models but also for guiding future research toward developing approaches that provide genuine scientific and practical utility.

**Figure 1.**
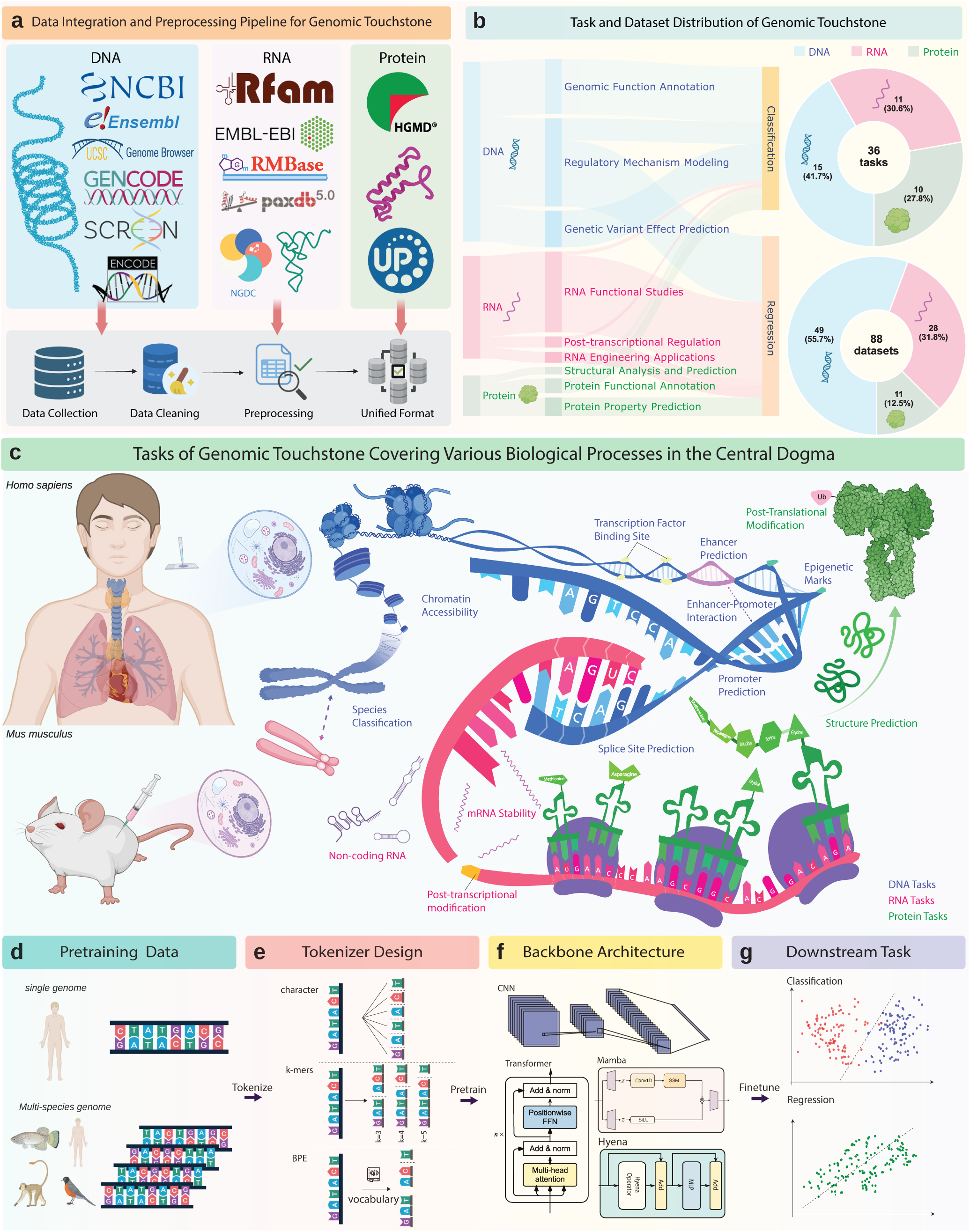
**Overview of Genomic Touchstone**. **a**, The data integration pipeline collects raw resources from omics repositories, performs data cleaning, and converts DNA, RNA and protein sequences into a unified format that preserves consistent meta-data for downstream use. **b**, Composition of the Genomic Touchstone benchmark, which comprises 36 tasks and 88 datasets. The left panel groups the tasks into nine main categories, while the right panel visualizes their distribution across the DNA, RNA, and protein modalities. **c**, Graphical representation of central-dogma coverage considered for benchmark tasks. **d-g**, The pipeline of our Genomic Touchstone benchmarking framework, including the key component categories at each stage of the framework: **d**, Composition of pretraining data, including human and multi-species genomic sequences; **e**, Tokenization strategies adopted by gLMs. **f**, Model architectures utilized across all evaluated gLMs. **g**, Fundamental groups of downstream tasks.

**Figure 2.**
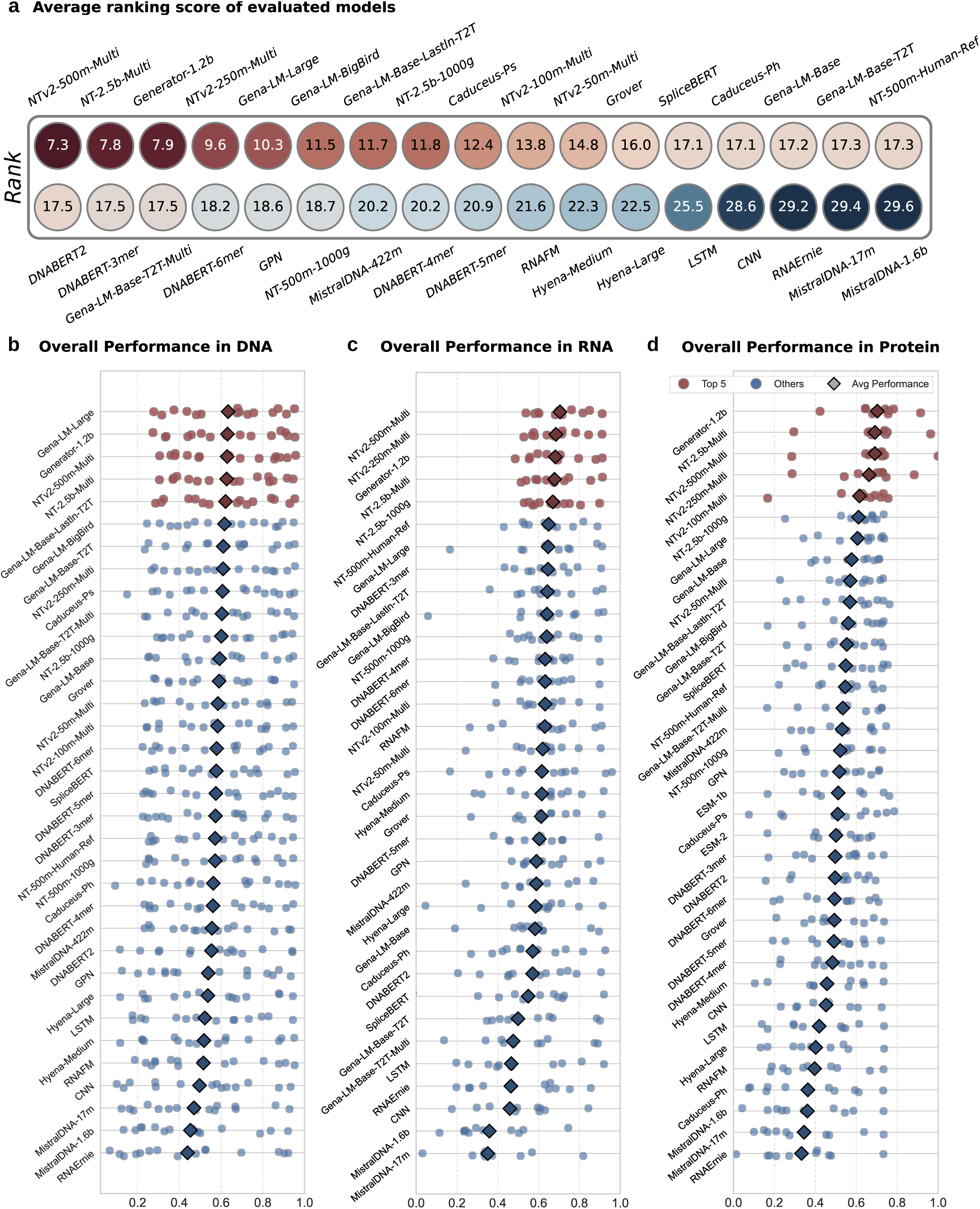
Benchmarking Results of gLMs. **a**, Average ranking of models based on their overall performance. Each circle denotes a model, colored by rank (darker indicating superior performance), with the numeric rank displayed at the center. **b-d**, Distribution of task-specific performance (dots) and average performance (black diamonds) for each model on DNA (**b**), RNA (**c**), and protein (**d**) benchmarks. Top five performing models per modality are highlighted in red, others in blue. transfer of information underpins every biological process and illustrates the intrinsic link between a genome’s sequence and the proteome it encodes (Fig. 1c). Given that the genome, particularly the coding DNA sequences (CDS), encodes all proteins, gLMs pretrained on genomic data have the potential to predict not only DNA sequence properties but also protein characteristics. To assess how effectively gLMs can handle protein-related tasks, we curated a series of tasks addressing various facets of protein biology. For this purpose, we translated protein-level datasets into a DNA-based format by extracting the corresponding CDS for each protein, following the approach of previous work^23^. The precise protocol for sequence retrieval is detailed in the Methods section. Specifically, our task suite includes three distinct task categories (Fig. 1b). First, **Structural Analysis and Prediction** evaluates the ability of gLMs to infer secondary and tertiary protein structures directly from sequence data. In addition, **Protein Functional Annotation** assesses how effectively the models can identify and assign biological functions to proteins based on their sequence features. Finally, **Protein Property Prediction** focuses on estimating key physicochemical attributes such as stability, solubility, and binding affinity, thereby demonstrating the models’ capacity to predict the impact of sequence variations on protein function.

## Results

### The Overview of Genomic Touchstone

Genomic Touchstone is an expansive benchmarking framework designed to deliver a thorough and fair assessment of gLMs in tackling real-world human-centric biological challenges. Motivated by the central dogma, which describes the seamless flow of genetic information from DNA to RNA to protein, Genomic Touchstone is the first benchmark to incorporate tasks spanning DNA, RNA, and protein sequence modalities (Fig. 1) to rigorously evaluate the performance and versatility of current gLMs. At the heart of these challenges lies the complexity of the genome itself^19^. DNA is the fundamental molecule of life, carrying the genetic instructions essential for cellular function, development, and evolution. It contains diverse elements such as protein-coding genes, non-coding RNAs, and various regulatory motifs (Fig. 1c). Genomic function annotation is dedicated to identifying and categorizing these components to reveal their roles in cellular processes. High-performing gLMs can leverage knowledge gained from large genomic datasets to detect intricate sequence patterns^6,8,10^. This capability yields a detailed map of functional elements across the genome and is critical both for advancing theoretical genomics and for practical applications such as disease diagnosis. Moreover, gene expression is controlled by complex regulatory networks that include promoters, enhancers, and other elements. Accurately modeling these mechanisms demands that gLMs capture long-range dependencies and context-specific interactions, which is essential for understanding how genes are activated or silenced in different cellular environments^20^. In addition, variations in the DNA sequence, even at a single nucleotide level, can have profound implications for gene function and lead to diseases, making the prediction of the effects of these genetic variants crucial for understanding pathogenesis and developing targeted therapies^21^. gLMs that excel in this domain should be able to analyze the subtle differences introduced by mutations, providing insights into their potential impact on gene regulation and phenotypes, and thus offer a predictive capability vital for translating computational genomic insights into actionable clinical strategies. Therefore, our evaluation of the gLMs in the level of DNA focuses on three categories: **Genomic Function Annotation**, **Regulatory Mechanism Modeling**, and **Genetic Variant Effect Prediction** (Fig. 1b). Collectively, these three categories encompass 15 tasks that challenge the models to accurately decipher and interpret the diverse aspects of genomic information.

Unlike DNA, which mainly serves as a repository for genetic information, RNA actively converts these instructions into functional proteins, modulates gene expression and regulates cellular responses to both internal and external stimuli (Fig. 1c). As an essential intermediary between DNA and proteins, RNA plays a critical role in numerous biological processes, including protein synthesis, enzyme activity, and gene regulation. Thus, unraveling the diverse functions of RNA is vital for understanding cellular complexity and the underlying mechanisms of diseases. From a linguistic perspective, the vocabularies of RNA and DNA differ by a single letter: *uracil (U)* in RNA replaces *thymine (T)* in DNA. However, the structural similarity between uracil and thymine, both pyrimidines that pair with adenine, suggests they are functionally analogous. Based on this similarity, we hypothesize that RNA and DNA may be represented within a unified semantic space. This suggests that gLMs pretrained on DNA could generalize effectively to RNA-related tasks. Inspired by BEACON^22^, we collected three major categories of RNA tasks: **RNA Functional Studies**, **Post-transcriptional Regulation** and **RNA Engineering Applications** (Fig. 1b). **RNA Functional Studies** analyse how RNA modulates gene expression and contributes to disease, providing mechanistic insight for therapeutic intervention. In parallel, **Post-transcriptional Regulation** focuses on RNA processing, stability, and modification events that occur after transcription, all of which shape RNA lifespan and regulatory output. Finally, **RNA Engineering Applications** explore RNA’s potential in synthetic biology to enhance its utility in biotechnology and medicine and address complex biological challenges.

Proteins and DNA are fundamentally interconnected through the central dogma of molecular biology^15^, which describes the flow of genetic information from DNA to RNA and ultimately to protein. In this process, DNA serves as a stable repository of genetic instructions, which are first transcribed into messenger RNA and subsequently translated into proteins. Proteins, in turn, carry out critical cellular functions ranging from catalysis and structural support to signaling and regulation. This seamless

Building on our carefully curated datasets, we selected a diverse set of gLMs, along with two baseline models (CNN and LSTM), to evaluate their performance on human-relevant tasks. To ensure a comprehensive and biologically meaningful assessment, we focused on gLMs that were pretrained on human genomic data, including models trained exclusively on the human reference genome as well as those using multi-species corpora that contain substantial human sequence content, as detailed in the Methods section. The selected models span a wide range of architectures, from traditional CNN and LSTM to transformer-based models and emerging structures such as BigBird, Hyena, and Mamba, each paired with different tokenization strategies (e.g., char, k-mer and BPE). Model sizes range from compact architectures with millions of parameters to large-scale models with up to 2.5 billion parameters, enabling us to examine how capacity influences biological sequence modeling. This collection of human-pretrained models allows us to systematically evaluate how architectural choices, tokenization schemes, and pretraining data impact downstream performance in human functional genomics. By finetuning all models on a shared benchmark suite—including tasks spanning DNA, RNA, and protein levels—we isolated the effects of specific modeling factors such as architecture design, pretraining strategy, and parameter scales. The inclusion of RNA and protein language models further enabled us to explore how well these models generalize across molecular modalities. Notably, no single model consistently outperformed others across all tasks (Fig. 2a); even the top-performing model achieved an average rank of 7.3, underscoring the trade-offs and specialization inherent to current genomic architectures. This variability highlights the importance of multi-dimensional evaluation and suggests that different tasks may benefit from different inductive biases. Finally, we extended our analysis to assess whether the scaling laws observed in LLMs apply to the genomic domain, providing insights into how model capacity and data composition jointly affect generalizability in both health-related and broader biological settings.

### Genomic Language Models Exhibit Superior Performance on DNA Tasks

#### Genomic Function Annotation

**Promoter Annotation** is a fundamental task aimed at classifying promoter regions based on key functional elements. This task comprises two distinct levels of analysis: a broader full promoter annotation and a more focused core promoter annotation. The former involves identifying elements like the TATA box, Initiator, CCAAT box, and GC box within the full promoter structure. The latter, core promoter annotation, is a more challenging endeavor as it concentrates on predicting the minimal, essential sequences for transcription initiation within a narrow window (-34 bp to +35 bp) immediately surrounding the transcription start site (TSS)^24^. As shown in Fig. 3a, in the promoter annotation task, NTv2-500m-Multi achieves the highest F1 score (0.863), followed closely by NTv2-250m-Multi (0.846) and Generator-1.2b (0.841). A large group of models, including DNABERT variants and Gena-LM series, also perform competitively within the 0.800–0.820 range. However, as presented in Table 3, models such as MistralDNA-1.6b (0.329), MistralDNA-422m (0.520), and LSTM (0.577) exhibit substantial performance gaps, suggesting that promoter annotation is a task where model design and training data have a pronounced impact on outcomes. **Enhancer Types Classification** is another fundamental task focused on distinguishing between tissue-specific and tissue-invariant enhancers, as well as non-enhancer regions. Enhancers are regulatory DNA elements that increase the transcriptional activity of target genes, often functioning over long genomic distances^25^. Tissue-specific enhancers contribute to cell-type–specific gene expression, while tissue-invariant enhancers are active across multiple tissues, reflecting core regulatory programs. As shown in Fig. 3a and Table 4, in the enhancer type classification task, all models achieve relatively similar performance, with DNABERT2 attaining the highest F1 score of 0.552, and MistralDNA-1.6b obtaining the lowest score of 0.461, resulting in a narrow performance gap of 9.1%. Top-performing models such as NT-2.5b-Multi (0.543) and Gena-LM-Base-T2T (0.539) achieve comparable F1 scores, while traditional models like LSTM (0.482) and RNAErnie (0.479) also perform competitively, trailing only slightly behind. This suggests that the task is relatively straightforward, limiting the performance advantage of more complex gLM architectures. Given its 1.6 billion parameters, the model’s subpar performance is surprising. One possible explanation is its pretraining solely on the human genome (hg38) using only 200 bp sequences, which limits its ability to capture long-range dependencies critical for regulatory element modeling.

**Figure 3.**
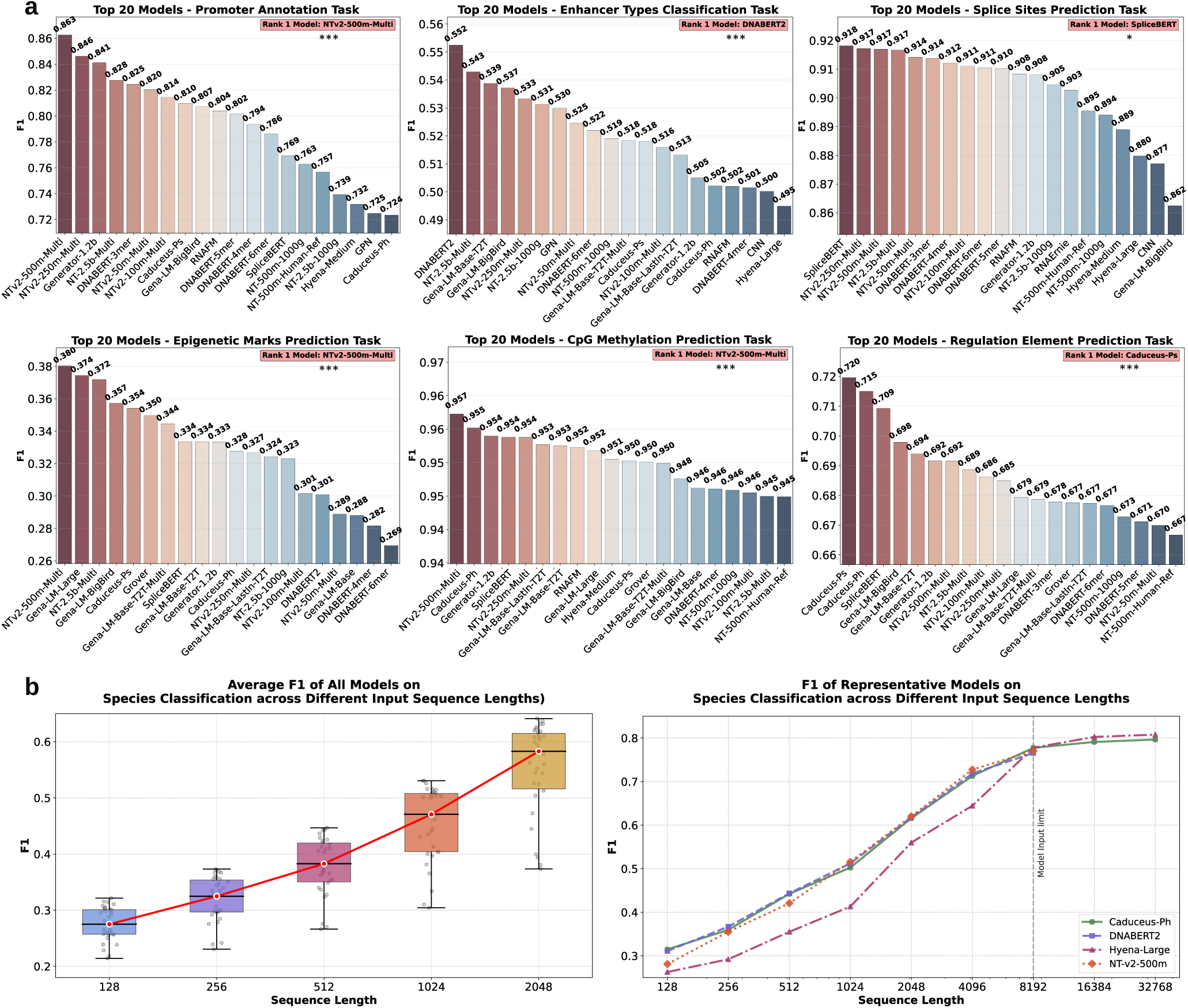
Model Performance on Genomic Function Annotation Tasks. **a**, Top 20 performing models on six representative Genomic Function Annotation benchmarks, measured by F1 score: promoter annotation, enhancer type classification, splice site prediction, epigenetic mark prediction, CpG methylation prediction, and regulatory element prediction (***: *P <* 0.001, *: *P <* 0.05). Performance varies by task, with models such as NTv2-500m-Multi, DNABERT, SpliceBERT, and Caduceus-Ps each achieving top rank in different contexts, indicating that distinct architectural biases and training regimes specialize models toward particular genomic functions. **b**, Species classification performance across varying input sequence lengths. Left: Distribution of the average F1 scores for multiple models at increasing sequence lengths, with the red trend line indicating mean F1 gain. Right: Increasing the input sequence length substantially boosts the F1 score for species classification, with the effect being most pronounced for models with long-context capability. The advantage of Caduceus-Ph and Hyena-Large becomes exclusive at sequences longer than 8,192 base pairs, as other models could not support these extended input lengths. This demonstrates that their architectural design is superior for leveraging long-range context to identify species-specific genomic patterns.

**Table 1.**
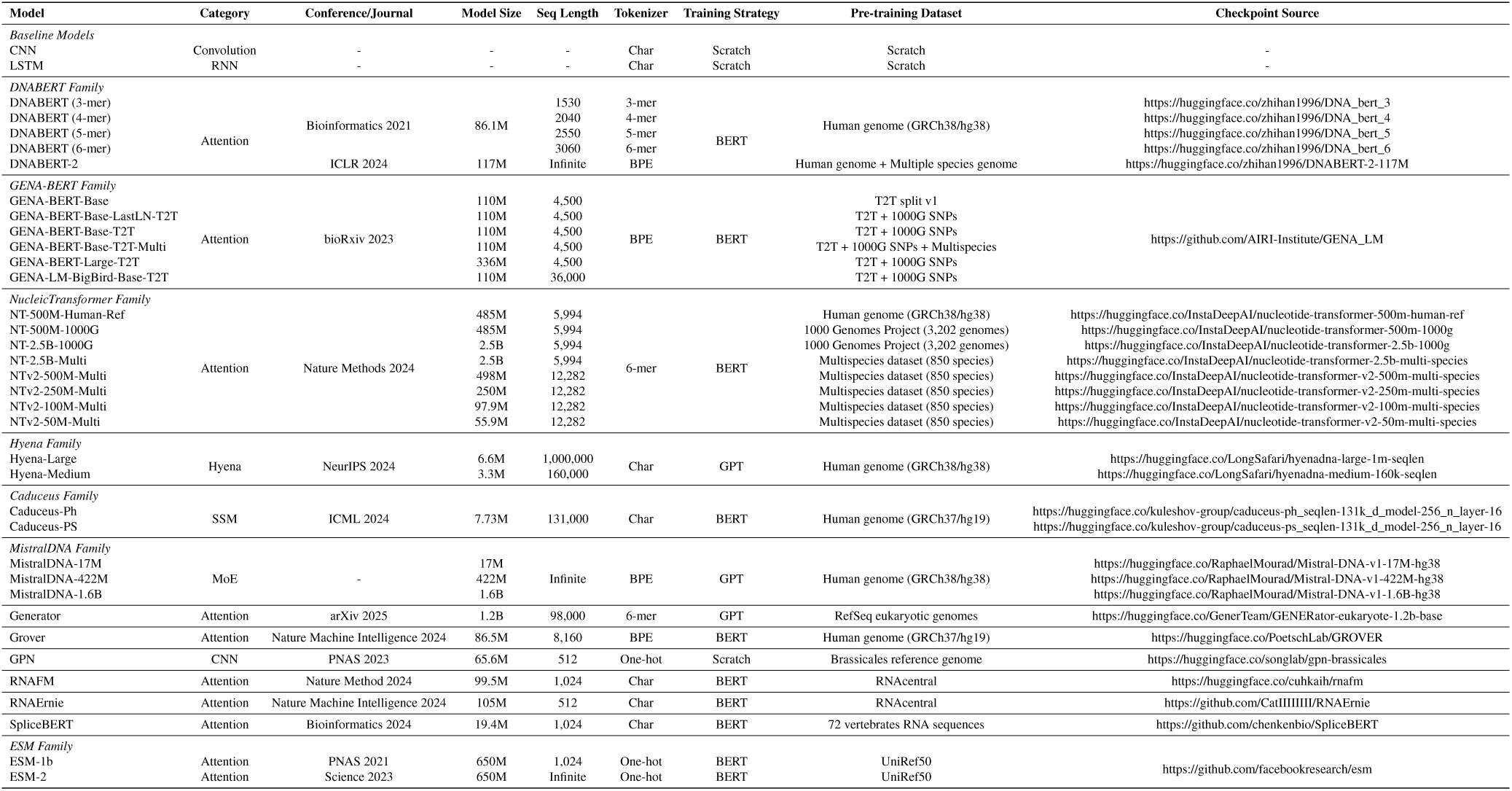
Model List.

**Table 2.** Task and Dataset Information.

| Task | Dataset | Type | Training size | Testing size | Validation size | Median length | Max Length |
| --- | --- | --- | --- | --- | --- | --- | --- |
| DNA |  |  |  |  |  |  |  |
| Promoter Annotation | Core_Promoter_Annotation | Sequence Classification (4 classes) | 34,736 | 4,342 | 4,342 | 70 | 70 |
|  | Promoter_Annotation | Sequence Multilabel Classification (6 classes) | 35,508 | 4,444 | 4,440 | 500 | 500 |
| Enhancers Types Classification | Enhancers_Types | Sequence Classification (3 classes) | 27,000 | 3,000 | 3,000 | 400 | 400 |
| Splice Sites Prediction | Splice_Sites_All | Sequence Multilabel Classification (3 classes) | 80,000 | 10,000 | 10,000 | 512 | 512 |
|  | Splice_Sites_Donor | Sequence Classification (2 classes) | 80,000 | 10,000 | 10,000 | 512 | 512 |
|  | Splice_Sites_Acceptor | Sequence Classification (2 classes) | 80,000 | 10,000 | 10,000 | 512 | 512 |
| Epigenetic Marks Prediction | Histone_annotation | Sequence Multilabel Classification (10 classes) | 96,642 | 12,080 | 12,081 | 1,024 | 1,024 |
| CpG Methylation Prediction | CpG_methylation | Sequence Multilabel Classification (7 classes) | 74,309 | 10,622 | 10,971 | 512 | 512 |
| Regulation Element Prediction | Regulation | Sequence Classification (4 classes) | 125,336 | 15,668 | 15,667 | 1,024 | 1,024 |
|  | Regulation_multilabel | Sequence Multilabel Classification (4 classes) | 277,660 | 34,708 | 34,707 | 1,024 | 1,024 |
| Species Classification | Species_Classification | Sequence Classification (6 classes) | 40,000 | 5,000 | 5,000 | 128/256/512/1,024/2,048 | 128/256/512/1,024/2,048 |
| Regulatory Activity Prediction | GM12878 | Sequence Regression | 98,996 | 29,112 | 9,866 | 2,114 | 2,114 |
|  | H1ESC | Sequence Regression | 133,947 | 31,474 | 13,312 | 2,114 | 2,114 |
|  | HEPG2 | Sequence Regression | 129,733 | 40,206 | 14,644 | 2,114 | 2,114 |
|  | IMR90 | Sequence Regression | 136,706 | 39,588 | 14,012 | 2,114 | 2,114 |
|  | K562 | Sequence Regression | 141,691 | 36,747 | 15,883 | 2,114 | 2,114 |
| Enhancer Activity Prediction | DeepSTAR_HCT116 | Sequence Regression | 17,226 | 2,154 | 2,153 | 249 | 249 |
|  | HCT116_Brd2 | Sequence Regression | 4,999 | 624 | 626 | 1,199 | 1,199 |
|  | HCT116_Brd4 | Sequence Regression | 4,999 | 624 | 626 | 1,199 | 1,199 |
|  | HCT116_CDK7 | Sequence Regression | 4,999 | 624 | 626 | 1,199 | 1,199 |
|  | HCT116_CDK9 | Sequence Regression | 4,999 | 624 | 626 | 1,199 | 1,199 |
|  | HCT116_Med14 | Sequence Regression | 4,999 | 624 | 626 | 1,199 | 1,199 |
|  | HCT116_Parental | Sequence Regression | 4,999 | 624 | 626 | 1,199 | 1,199 |
|  | HCT116_p300CBP | Sequence Regression | 4,999 | 624 | 626 | 1,199 | 1,199 |
| Enhancer Promoter Interaction Prediction | GM12878 | Sequence Classification (2 classes) | 10,000 | 2,000 | 2,000 | 3,000 | 3,000 |
|  | H1ESC | Sequence Classification (2 classes) | 10,000 | 2,000 | 2,000 | 3,000 | 3,000 |
|  | HEPG2 | Sequence Classification (2 classes) | 10,000 | 2,000 | 2,000 | 3,000 | 3,000 |
|  | IMR90 | Sequence Classification (2 classes) | 10,000 | 2,000 | 2,000 | 3,000 | 3,000 |
|  | K562 | Sequence Classification (2 classes) | 10,000 | 2,000 | 2,000 | 3,000 | 3,000 |
|  | NHEK | Sequence Classification (2 classes) | 10,000 | 2,000 | 2,000 | 3,000 | 3,000 |
| Bulk RNA Expression Prediction | Bulk_RNA_Expression_Prediction | Sequence Regression | 20,544 | 2,283 | 990 | 128/256/512/1,024/2,048 | 128/256/512/1,024/2,048 |
| Genetic Disease Classification | Breast_Disease_Type_Classification | Sequence Classification (4 classes) | 14,779 | 1,848 | 1,847 | 512 | 512 |
|  | Cardiovascular_Disorders_Classification | Sequence Classification (8 classes) | 36,480 | 4,561 | 4,560 | 512 | 512 |
|  | Kidney_Disease_Type_Classification | Sequence Classification (7 classes) | 11,671 | 1,459 | 1,459 | 512 | 512 |
| Pathogenicity Classification | Pathogenicity_Classification | Sequence Classification (4 classes) | 47,121 | 5,890 | 5,890 | 512 | 512 |
| Splice Variant Effect Prediction | Exonic | Sequence Classification (2 classes) | 7,232 | 904 | 904 | 1,024 | 1,024 |
|  | Intronic | Sequence Classification (2 classes) | 11,581 | 1,448 | 1,448 | 1,024 | 1,024 |
| BRCA1/2 SNV Functional Impact Classification | BRCA1_coding | Sequence Classification (2 classes) | 2,214 | 277 | 277 | 1,024 | 1,024 |
|  | BRCA1_noncoding | Sequence Classification (2 classes) | 900 | 112 | 113 | 1,024 | 1,024 |
|  | BRCA2_coding | Sequence Classification (2 classes) | 4,939 | 617 | 618 | 1,024 | 1,024 |
|  | BRCA2_noncoding | Sequence Classification (2 classes) | 530 | 66 | 67 | 1,024 | 1,024 |
| RNA |  |  |  |  |  |  |  |
| Mean Ribosome Loading Prediction | Mean_Ribosome_Loading | Sequence Regression | 76,319 | 7,600 | 7,600 | 61 | 100 |
|  | Mean_Ribosome_Loading_Human | Sequence Regression | 6,080 | 760 | 760 | 62.5 | 100 |
| mRNA Stability Prediction | mRNA_Stability | Sequence Regression | 52,283 | 6,536 | 6,536 | 1,239 | 3,066 |
| Abundance Prediction | PA_athaliana | Sequence Regression | 8,703 | 1,088 | 1,087 | 1,128 | 3,456 |
|  | PA_dmelanogaster | Sequence Regression | 7,854 | 982 | 981 | 1,173 | 4,287 |
|  | PA_ecoli | Sequence Regression | 2,756 | 344 | 344 | 805 | 2,145 |
|  | PA_hsapiens | Sequence Regression | 8,959 | 1,120 | 1,120 | 1,203 | 4,251 |
|  | PA_scervisiae | Sequence Regression | 3,751 | 469 | 469 | 1,182 | 3,702 |
|  | TA_athaliana | Sequence Regression | 9,625 | 1,203 | 1,202 | 1,035 | 3,156 |
|  | TA_dmelanogaster | Sequence Regression | 7,474 | 934 | 934 | 1,221 | 4,857 |
|  | TA_ecoli | Sequence Regression | 2,680 | 335 | 335 | 819 | 2,622 |
|  | TA_hsapiens | Sequence Regression | 4,168 | 521 | 520 | 1,083 | 4,500 |
|  | TA_hvolcanii | Sequence Regression | 2,424 | 303 | 303 | 747 | 2,079 |
|  | TA_ppastoris | Sequence Regression | 3,557 | 445 | 444 | 1,191 | 3,351 |
|  | TA_scervisiae | Sequence Regression | 4,141 | 517 | 517 | 1,080 | 3,510 |
| Non-coding RNA Classification | NoncodingRNAFamily | Sequence Classification (13 classes) | 5,670 | 2,600 | 650 | 128 | 1,182 |
| Expression Level and Translation Efficiency Regulation Prediction | EL_HEK | Sequence Regression | 11,528 | 1,441 | 1,441 | 100 | 100 |
|  | EL_Muscle | Sequence Regression | 1,005 | 126 | 126 | 100 | 100 |
|  | EL_pc3 | Sequence Regression | 10,063 | 1,258 | 1,258 | 100 | 100 |
|  | TE_HEK | Sequence Regression | 11,528 | 1,441 | 1,441 | 100 | 100 |
|  | TE_Muscle | Sequence Regression | 1,005 | 126 | 126 | 100 | 100 |
|  | TE_pc3 | Sequence Regression | 10,063 | 1,258 | 1,258 | 100 | 100 |
| Modification Prediction | Modification | Sequence Multilabel Classification (12 classes) | 247,568 | 30,946 | 30,946 | 101 | 101 |
| APA Isoform Prediction | Isoform | Sequence Regression | 182,710 | 22,839 | 22,839 | 186 | 186 |
| SARS-CoV-2 Vaccine Degradation Prediction | CoV_Vaccine_Degradation | Sequence Regression | 1,920 | 240 | 240 | 81 | 81 |
| Programmable RNA Switches Prediction | ProgrammableRNASwitches | Sequence Regression | 73,227 | 9,154 | 9,153 | 148 | 148 |
| Tc-Riboswitches Prediction | Tc_Riboswitches | Sequence Regression | 284 | 36 | 36 | 72 | 75 |
| mRFP Expression Prediction | mRFP_Expression | Sequence Regression | 1,167 | 146 | 146 | 678 | 678 |
| Protein |  |  |  |  |  |  |  |
| Secondary Structure Analysis | Structure Prediction | Sequence Classification (3 classes) | 1,347 | 337 | 422 | 2,160 | 2,160 |
| Transmembrane Region Prediction | Transmembrane | Sequence Classification (6 classes) | 1,241 | 311 | 389 | 2,160 | 2,160 |
| Protein Variant Classification | Variant | Sequence Classification (2 classes) | 2,544 | 636 | 796 | 2,160 | 2,160 |
| Post-translational Modifications Prediction | PTM Prediction | Sequence Classification (3 classes) | 2,635 | 659 | 824 | 2,160 | 2,160 |
| Protein Domain Classification | Domain | Sequence Classification (4 classes) | 351 | 88 | 110 | 2,160 | 2,160 |
| Enzyme Classification | Enzyme Classification | Sequence Classification (4 classes) | 723 | 181 | 226 | 2,160 | 2,160 |
| Beta-Lactamase Activity Prediction | beta_lactamase_complete | Sequence Regression | 9,202 | 1,080 | 2,814 | 858 | 858 |
|  | beta_lactamase_unique | Sequence Regression | 3,457 | 1,080 | 865 | 858 | 858 |
| Fluorescence Prediction | Fluorescence | Sequence Regression | 21,464 | 27,217 | 5,366 | 714 | 714 |
| Protein Stability Prediction | Stability | Sequence Regression | 53,700 | 12,851 | 2,512 | 135 | 135 |
| Melting Point Prediction | Melting Point | Sequence Regression | 9,432 | 1,648 | 1,064 | 1,176 | 1,176 |

**Table 3.**
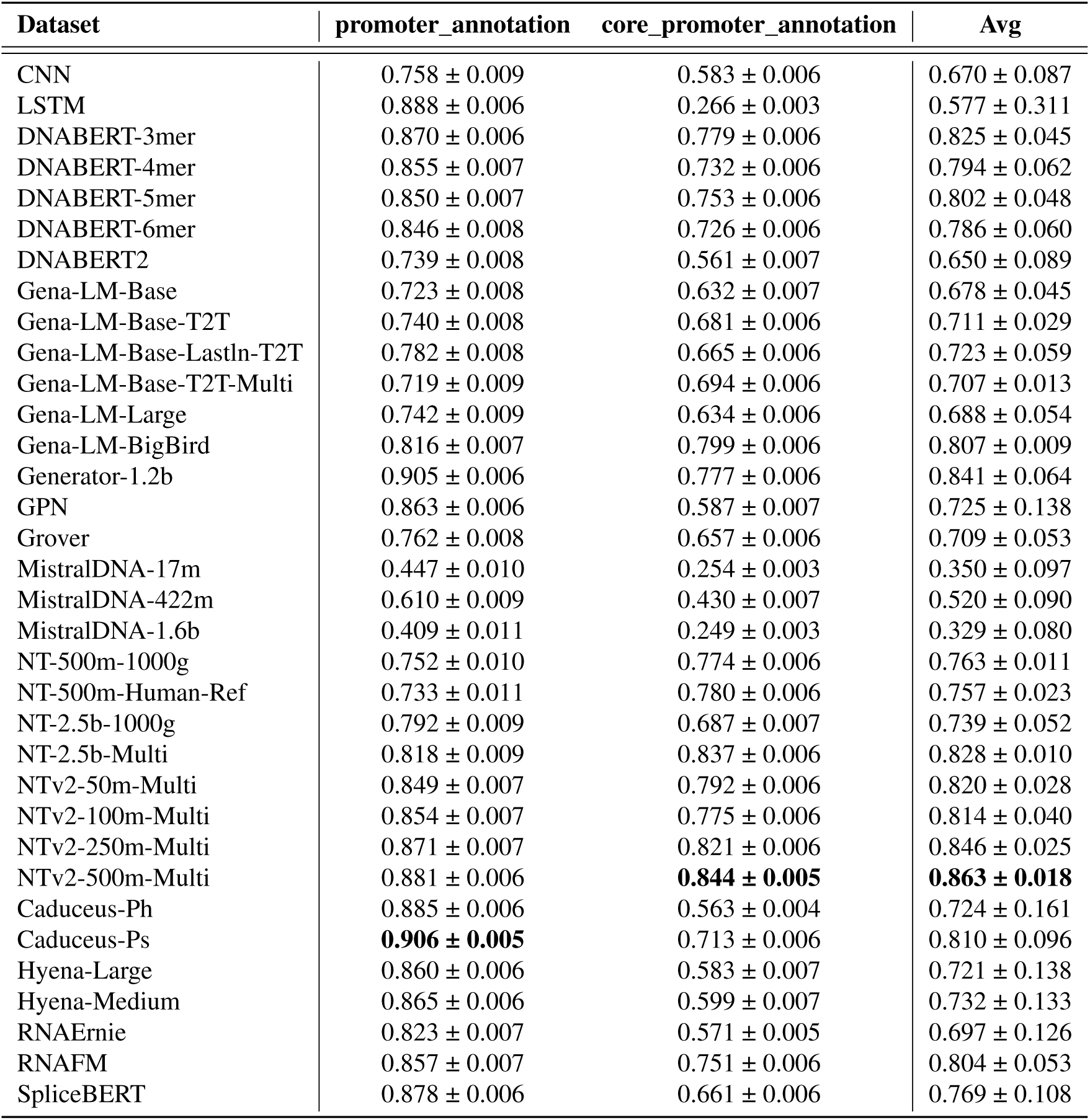
F1 score of the evaluated models on Promoter Annotation Tasks.

**Table 4.** F1 score of the evaluated models on Enhancer Types Classification.

| Dataset | enhancers_types |
| --- | --- |
| CNN | 0.500 ± 0.011 |
| LSTM | 0.482 ± 0.009 |
| DNABERT-3mer | 0.488 ± 0.010 |
| DNABERT-4mer | 0.501 ± 0.012 |
| DNABERT-5mer | 0.482 ± 0.006 |
| DNABERT-6mer | 0.522 ± 0.013 |
| DNABERT2 | **0.552 ± 0.016** |
| Gena-LM-Base | 0.494 ± 0.010 |
| Gena-LM-Base-T2T | 0.539 ± 0.014 |
| Gena-LM-Base-LastIn-T2T | 0.513 ± 0.012 |
| Gena-LM-Base-T2T-Multi | 0.518 ± 0.013 |
| Gena-LM-Large | 0.482 ± 0.006 |
| Gena-LM-BigBird | 0.537 ± 0.013 |
| Generator-1.2b | 0.505 ± 0.012 |
| GPN | 0.530 ± 0.014 |
| Grover | 0.486 ± 0.005 |
| MistralDNA-17m | 0.479 ± 0.008 |
| MistralDNA-422m | 0.483 ± 0.005 |
| MistralDNA-1.6b | 0.461 ± 0.008 |
| NT-500m-1000g | 0.519 ± 0.014 |
| NT-500m-Human-Ref | 0.482 ± 0.006 |
| NT-2.5b-1000g | 0.531 ± 0.015 |
| NT-2.5b-Multi | 0.543 ± 0.017 |
| NTv2-50m-Multi | 0.525 ± 0.014 |
| NTv2-100m-Multi | 0.516 ± 0.014 |
| NTv2-250m-Multi | 0.533 ± 0.011 |
| NTv2-500m-Multi | 0.493 ± 0.008 |
| Caduceus-Ph | 0.502 ± 0.012 |
| Caduceus-Ps | 0.518 ± 0.011 |
| Hyena-Large | 0.495 ± 0.005 |
| Hyena-Medium | 0.484 ± 0.005 |
| RNAErnie | 0.479 ± 0.005 |
| RNAFM | 0.502 ± 0.009 |
| SpliceBERT | 0.493 ± 0.009 |

**Splice Sites Prediction**. Splicing is a fundamental process in eukaryotic gene expression, where non-coding introns are removed and coding exons are joined together to form mature mRNA. Accurate recognition of splice sites is essential for understanding the diversity of transcript isoforms and alternative splicing, which contribute to functional complexity in the genome^26^. We focus on evaluating gLMs’ ability to identify splice donor and acceptor sites within genomic sequences. As shown in Fig. 3a, in the splice site prediction task, most models perform exceptionally well, with SpliceBERT achieving the highest F1 score of 0.918. This strong performance is likely attributable to its pretraining on splice site–related sequences, which directly aligns with the evaluation objective. Several other models—including NTv2-250m-Multi, NTv2-500m-Multi, and DNABERT variants—also achieve scores above 0.910, reflecting both the tractability of the task and the models’ capacity to capture splicing-relevant sequence features. In contrast, as illustrated in Table 5, MistralDNA-1.6b (0.604) performs notably worse, suggesting that large model size alone does not guarantee success in tasks requiring fine-grained sequence resolution and specialized signal capture.

**Table 5.** F1 score of the evaluated models on Splice Sites Prediction.

| Dataset | splice_sites_acceptor | splice_sites_donor | splice_sites_all | Avg |
| --- | --- | --- | --- | --- |
| CNN | 0.890 ± 0.003 | 0.847 ± 0.004 | 0.894 ± 0.003 | 0.877 ± 0.022 |
| LSTM | 0.840 ± 0.004 | 0.797 ± 0.005 | 0.844 ± 0.003 | 0.827 ± 0.021 |
| DNABERT-3mer | 0.923 ± 0.003 | 0.891 ± 0.003 | 0.928 ± 0.002 | 0.914 ± 0.017 |
| DNABERT-4mer | 0.921 ± 0.003 | 0.892 ± 0.003 | 0.924 ± 0.002 | 0.912 ± 0.015 |
| DNABERT-5mer | 0.918 ± 0.003 | 0.888 ± 0.004 | 0.925 ± 0.002 | 0.910 ± 0.016 |
| DNABERT-6mer | 0.918 ± 0.003 | 0.888 ± 0.003 | 0.927 ± 0.002 | 0.911 ± 0.017 |
| DNABERT2 | 0.807 ± 0.005 | 0.771 ± 0.005 | 0.820 ± 0.004 | 0.799 ± 0.021 |
| Gena-LM-Base | 0.812 ± 0.004 | 0.780 ± 0.005 | 0.834 ± 0.003 | 0.809 ± 0.022 |
| Gena-LM-Base-T2T | 0.857 ± 0.004 | 0.820 ± 0.004 | 0.862 ± 0.003 | 0.846 ± 0.019 |
| Gena-LM-Base-LastIn-T2T | 0.847 ± 0.004 | 0.818 ± 0.004 | 0.859 ± 0.003 | 0.841 ± 0.017 |
| Gena-LM-Base-T2T-Multi | 0.850 ± 0.004 | 0.811 ± 0.004 | 0.859 ± 0.003 | 0.840 ± 0.021 |
| Gena-LM-Large | 0.850 ± 0.004 | 0.828 ± 0.004 | 0.867 ± 0.003 | 0.849 ± 0.016 |
| Gena-LM-BigBird | 0.867 ± 0.004 | 0.839 ± 0.004 | 0.882 ± 0.003 | 0.862 ± 0.018 |
| Generator-1.2b | 0.918 ± 0.003 | 0.876 ± 0.004 | 0.930 ± 0.002 | 0.908 ± 0.023 |
| GPN | 0.853 ± 0.004 | 0.824 ± 0.004 | 0.868 ± 0.003 | 0.848 ± 0.018 |
| Grover | 0.848 ± 0.004 | 0.813 ± 0.005 | 0.863 ± 0.003 | 0.841 ± 0.021 |
| MistralDNA-17m | 0.680 ± 0.006 | 0.638 ± 0.006 | 0.693 ± 0.005 | 0.670 ± 0.023 |
| MistralDNA-422m | 0.739 ± 0.005 | 0.679 ± 0.006 | 0.763 ± 0.004 | 0.727 ± 0.036 |
| MistralDNA-1.6b | 0.616 ± 0.006 | 0.570 ± 0.007 | 0.624 ± 0.005 | 0.604 ± 0.024 |
| NT-500m-1000g | 0.902 ± 0.003 | 0.864 ± 0.004 | 0.917 ± 0.002 | 0.894 ± 0.022 |
| NT-500m-Human-Ref | 0.898 ± 0.003 | 0.871 ± 0.004 | 0.918 ± 0.002 | 0.895 ± 0.020 |
| NT-2.5b-1000g | 0.915 ± 0.003 | 0.879 ± 0.004 | 0.920 ± 0.002 | 0.905 ± 0.018 |
| NT-2.5b-Multi | 0.925 ± 0.003 | 0.893 ± 0.004 | 0.933 ± 0.002 | 0.917 ± 0.017 |
| NTv2-50m-Multi | 0.922 ± 0.003 | 0.892 ± 0.003 | 0.929 ± 0.002 | 0.914 ± 0.016 |
| NTv2-100m-Multi | 0.924 ± 0.003 | 0.884 ± 0.004 | 0.926 ± 0.002 | 0.911 ± 0.019 |
| NTv2-250m-Multi | 0.922 ± 0.003 | 0.896 ± 0.003 | **0.934 ± 0.002** | 0.917 ± 0.016 |
| NTv2-500m-Multi | 0.927 ± 0.003 | 0.892 ± 0.004 | 0.932 ± 0.002 | 0.917 ± 0.018 |
| Caduceus-Ph | 0.880 ± 0.004 | 0.778 ± 0.005 | 0.799 ± 0.004 | 0.819 ± 0.044 |
| Caduceus-Ps | 0.875 ± 0.004 | 0.821 ± 0.005 | 0.865 ± 0.003 | 0.854 ± 0.024 |
| Hyena-Large | 0.879 ± 0.004 | 0.870 ± 0.004 | 0.891 ± 0.003 | 0.880 ± 0.009 |
| Hyena-Medium | 0.908 ± 0.003 | 0.864 ± 0.004 | 0.895 ± 0.003 | 0.889 ± 0.018 |
| RNAErnie | 0.913 ± 0.003 | 0.877 ± 0.004 | 0.919 ± 0.002 | 0.903 ± 0.019 |
| RNAFM | 0.917 ± 0.003 | 0.888 ± 0.003 | 0.920 ± 0.002 | 0.908 ± 0.015 |
| SpliceBERT | **0.927 ± 0.003** | **0.899 ± 0.003** | 0.928 ± 0.002 | **0.918 ± 0.014** |

**Epigenetic Marks Prediction** is a pivotal computational task that leverages the representations learned by gLMs to decipher the complex regulatory networks modulating gene expression. By accurately predicting the presence of these marks on genomic sequences, we can gain insights into how different histone modifications interact, and ultimately reveal patterns related to gene expression and disease mechanisms^27^. As shown in Fig. 3a, in the epigenetic marks prediction task, overall model performance is relatively low, with the top model NTv2-500m-Multi achieving an F1 score of 0.380. Gena-LM-Large (0.374) and NT-2.5b-Multi (0.372) follow closely, while the majority of models fall below 0.350, indicating the challenging nature of this task. Notably, as demonstrated in Table 6, several models including RNAErnie (0.067), MistralDNA-1.6b (0.133), and CNN (0.133) perform poorly, highlighting substantial room for improvement in capturing epigenetic regulatory patterns. These results suggest that Epigenetic Marks Prediction remains a highly challenging task for gLMs.

**Table 6.** F1 score of the evaluated models on Epigenetic Marks Prediction.

| Dataset | Histone_annotation |
| --- | --- |
| CNN | 0.133 ± 0.003 |
| LSTM | 0.178 ± 0.004 |
| DNABERT-3mer | 0.250 ± 0.004 |
| DNABERT-4mer | 0.282 ± 0.004 |
| DNABERT-5mer | 0.240 ± 0.004 |
| DNABERT-6mer | 0.269 ± 0.004 |
| DNABERT2 | 0.301 ± 0.004 |
| Gena-LM-Base | 0.288 ± 0.004 |
| Gena-LM-Base-T2T | 0.334 ± 0.004 |
| Gena-LM-Base-LastIn-T2T | 0.324 ± 0.004 |
| Gena-LM-Base-T2T-Multi | 0.344 ± 0.004 |
| Gena-LM-Large | 0.374 ± 0.004 |
| Gena-LM-BigBird | 0.357 ± 0.004 |
| Generator-1.2b | 0.333 ± 0.004 |
| GPN | 0.125 ± 0.003 |
| Grover | 0.350 ± 0.004 |
| MistralDNA-17m | 0.123 ± 0.003 |
| MistralDNA-422m | 0.233 ± 0.004 |
| MistralDNA-1.6b | 0.133 ± 0.003 |
| NT-500m-1000g | 0.242 ± 0.004 |
| NT-500m-Human-Ref | 0.264 ± 0.004 |
| NT-2.5b-1000g | 0.323 ± 0.004 |
| NT-2.5b-Multi | 0.372 ± 0.004 |
| NTv2-50m-Multi | 0.289 ± 0.004 |
| NTv2-100m-Multi | 0.302 ± 0.004 |
| NTv2-250m-Multi | 0.327 ± 0.004 |
| NTv2-500m-Multi | **0.380 ± 0.004** |
| Caduceus-Ph | 0.328 ± 0.004 |
| Caduceus-Ps | 0.354 ± 0.004 |
| Hyena-Large | 0.225 ± 0.004 |
| Hyena-Medium | 0.224 ± 0.004 |
| RNAErnie | 0.067 ± 0.003 |
| RNAFM | 0.187 ± 0.004 |
| SpliceBERT | 0.334 ± 0.004 |

**Table 7.** F1 score of the evaluated models on CpG Methylation Prediction.

| Dataset | cpg_methylation |
| --- | --- |
| CNN | 0.933 ± 0.002 |
| LSTM | 0.944 ± 0.002 |
| DNABERT-3mer | 0.945 ± 0.002 |
| DNABERT-4mer | 0.946 ± 0.002 |
| DNABERT-5mer | 0.944 ± 0.002 |
| DNABERT-6mer | 0.944 ± 0.002 |
| DNABERT2 | 0.943 ± 0.002 |
| Gena-LM-Base | 0.946 ± 0.002 |
| Gena-LM-Base-T2T | 0.953 ± 0.001 |
| Gena-LM-Base-LastIn-T2T | 0.953 ± 0.002 |
| Gena-LM-Base-T2T-Multi | 0.950 ± 0.002 |
| Gena-LM-Large | 0.952 ± 0.001 |
| Gena-LM-BigBird | 0.948 ± 0.002 |
| Generator-1.2b | 0.954 ± 0.001 |
| GPN | 0.934 ± 0.002 |
| Grover | 0.950 ± 0.002 |
| MistralDNA-17m | 0.928 ± 0.002 |
| MistralDNA-422m | 0.935 ± 0.002 |
| MistralDNA-1.6b | 0.921 ± 0.002 |
| NT-500m-1000g | 0.946 ± 0.002 |
| NT-500m-Human-Ref | 0.945 ± 0.002 |
| NT-2.5b-1000g | 0.942 ± 0.002 |
| NT-2.5b-Multi | 0.945 ± 0.002 |
| NTv2-50m-Multi | 0.940 ± 0.002 |
| NTv2-100m-Multi | 0.946 ± 0.002 |
| NTv2-250m-Multi | 0.954 ± 0.002 |
| NTv2-500m-Multi | **0.957 ± 0.001** |
| Caduceus-Ph | 0.955 ± 0.002 |
| Caduceus-Ps | 0.950 ± 0.002 |
| Hyena-Large | 0.944 ± 0.002 |
| Hyena-Medium | 0.951 ± 0.002 |
| RNAErnie | 0.922 ± 0.002 |
| RNAFM | 0.952 ± 0.002 |
| SpliceBERT | 0.954 ± 0.001 |

**CpG Methylation Prediction** focuses on identifying the methylation status of CpG sites—critical epigenetic markers that influence gene expression without altering the underlying DNA sequence. DNA methylation, especially in CpG contexts, plays a vital role in a wide range of biological processes, including transcriptional regulation, embryonic development, genomic imprinting, and X-chromosome inactivation^28^. As shown in Fig. 3a, in the CpG methylation prediction task, nearly all models perform exceptionally well, with F1 scores tightly clustered above 0.920. The top-performing model, NTv2-500m-Multi, achieves an F1 score of 0.957, followed closely by Caduceus-Ph (0.955) and Generator-1.2b (0.954). The narrow performance range suggests that this task is relatively easy for current gLMs, and that many architectures are capable of effectively capturing the sequence features relevant to CpG methylation status.

**Regulation Element Prediction** aims at identifying core regulatory elements in the genome, including promoters, enhancers, CTCF binding sites, and open chromatin regions. These elements play pivotal roles in controlling gene expression by modulating the accessibility and activity of transcriptional machinery. Promoters initiate transcription, enhancers amplify gene expression in a context-specific manner, CTCF binding sites help establish chromatin architecture and insulation, while open chromatin regions mark accessible sites of potential regulatory activity. Accurate prediction of these regulatory elements is essential for decoding the gene regulatory landscape, understanding transcriptional regulation, and identifying functional non-coding regions that may underlie disease-associated genetic variants^29^. As shown in Fig. 3a, in the regulatory element prediction task, performance varies more widely across models. The best-performing model, Caduceus-Ps, achieves an F1 score of 0.720, followed closely by Caduceus-Ph (0.715) and SpliceBERT (0.709). While many top-tier gLMs cluster between 0.660 and 0.700, lower-performing models such as MistralDNA-1.6b (0.474), RNAErnie (0.525), and MistralDNA-17m (0.544) lag significantly behind (Table 8). This performance spread indicates that regulatory element modeling remains a moderately challenging task and may benefit more from architecture and pretraining design tailored to regulatory sequence patterns.

**Table 8.** F1 score of the evaluated models on Regulation Element Prediction.

| Dataset | regulation | regulation_multilabel | Avg |
| --- | --- | --- | --- |
| CNN | 0.620 ± 0.005 | 0.583 ± 0.002 | 0.602 ± 0.019 |
| LSTM | 0.648 ± 0.006 | 0.671 ± 0.003 | 0.660 ± 0.011 |
| DNABERT-3mer | 0.681 ± 0.006 | 0.675 ± 0.003 | 0.678 ± 0.003 |
| DNABERT-4mer | 0.666 ± 0.006 | 0.667 ± 0.003 | 0.667 ± 0.001 |
| DNABERT-5mer | 0.671 ± 0.006 | 0.671 ± 0.003 | 0.671 ± 0.000 |
| DNABERT-6mer | 0.675 ± 0.006 | 0.678 ± 0.003 | 0.677 ± 0.002 |
| DNABERT2 | 0.637 ± 0.006 | 0.686 ± 0.003 | 0.661 ± 0.024 |
| Gena-LM-Base | 0.617 ± 0.006 | 0.691 ± 0.003 | 0.654 ± 0.037 |
| Gena-LM-Base-T2T | 0.680 ± 0.006 | 0.708 ± 0.003 | 0.694 ± 0.014 |
| Gena-LM-Base-LastIn-T2T | 0.689 ± 0.006 | 0.665 ± 0.002 | 0.677 ± 0.012 |
| Gena-LM-Base-T2T-Multi | 0.663 ± 0.006 | 0.695 ± 0.003 | 0.679 ± 0.016 |
| Gena-LM-Large | 0.689 ± 0.006 | 0.670 ± 0.003 | 0.679 ± 0.009 |
| Gena-LM-BigBird | 0.682 ± 0.006 | 0.714 ± 0.003 | 0.698 ± 0.016 |
| Generator-1.2b | 0.709 ± 0.006 | 0.674 ± 0.003 | 0.692 ± 0.018 |
| GPN | 0.603 ± 0.006 | 0.581 ± 0.002 | 0.592 ± 0.011 |
| Grover | 0.675 ± 0.006 | 0.680 ± 0.003 | 0.677 ± 0.002 |
| MistralDNA-17m | 0.503 ± 0.006 | 0.586 ± 0.002 | 0.544 ± 0.042 |
| MistralDNA-422m | 0.613 ± 0.006 | 0.653 ± 0.003 | 0.633 ± 0.020 |
| MistralDNA-1.6b | 0.407 ± 0.006 | 0.541 ± 0.002 | 0.474 ± 0.067 |
| NT-500m-1000g | 0.694 ± 0.006 | 0.651 ± 0.003 | 0.673 ± 0.022 |
| NT-500m-Human-Ref | 0.669 ± 0.006 | 0.664 ± 0.003 | 0.667 ± 0.003 |
| NT-2.5b-1000g | 0.687 ± 0.006 | 0.641 ± 0.002 | 0.664 ± 0.023 |
| NT-2.5b-Multi | 0.684 ± 0.006 | 0.693 ± 0.003 | 0.689 ± 0.005 |
| NTv2-50m-Multi | 0.676 ± 0.005 | 0.664 ± 0.003 | 0.670 ± 0.006 |
| NTv2-100m-Multi | 0.698 ± 0.006 | 0.675 ± 0.003 | 0.686 ± 0.012 |
| NTv2-250m-Multi | 0.690 ± 0.006 | 0.680 ± 0.003 | 0.685 ± 0.005 |
| NTv2-500m-Multi | 0.707 ± 0.006 | 0.677 ± 0.003 | 0.692 ± 0.015 |
| Caduceus-Ph | 0.724 ± 0.006 | 0.706 ± 0.003 | 0.715 ± 0.009 |
| Caduceus-Ps | 0.719 ± 0.006 | **0.720 ± 0.003** | **0.720 ± 0.001** |
| Hyena-Large | 0.643 ± 0.006 | 0.664 ± 0.003 | 0.654 ± 0.011 |
| Hyena-Medium | 0.595 ± 0.006 | 0.645 ± 0.002 | 0.620 ± 0.025 |
| RNAErnie | 0.521 ± 0.005 | 0.529 ± 0.002 | 0.525 ± 0.004 |
| RNAFM | 0.620 ± 0.006 | 0.616 ± 0.003 | 0.618 ± 0.002 |
| SpliceBERT | **0.727 ± 0.006** | 0.691 ± 0.003 | 0.709 ± 0.018 |

**Species Classification** aims to distinguish subtle yet informative sequence differences across species, offering insights into evolutionary patterns and species-specific mutational signatures^30^. Despite high genomic similarity—for example, between humans and non-human primates—long DNA sequences contain distinct variations that enable accurate species identification. This task holds significant biological relevance for comparative genomics, species identification, and ancient DNA analysis, enhancing our understanding of evolutionary relationships and regulatory divergence across species. As shown in Table 9, in the species classification task, overall F1 score is relatively low across models, with the best-performing model, Gena-LM-Base-T2T-Multi, reaching only 0.462. Most models cluster between 0.430 and 0.450, indicating limited cross-species generalization. Notably, several models—including CNN (0.287), MistralDNA-1.6b (0.284), and RNAErnie (0.193)—perform substantially worse, suggesting that current gLMs struggle to distinguish species-level patterns and may require dedicated pretraining strategies or taxonomic priors to improve performance in this challenging task.

**Table 9.** F1 score of the evaluated models on Species Classification.

| Dataset | species_classification_128 | species_classification_256 | species_classification_512 | species_classification_1024 | species_classification_2048 | Avg |
| --- | --- | --- | --- | --- | --- | --- |
| CNN | 0.229 ± 0.006 | 0.242 ± 0.006 | 0.276 ± 0.006 | 0.310 ± 0.006 | 0.381 ± 0.007 | 0.287 ± 0.055 |
| LSTM | 0.269 ± 0.006 | 0.299 ± 0.006 | 0.363 ± 0.007 | 0.435 ± 0.007 | 0.563 ± 0.007 | 0.386 ± 0.105 |
| DNABERT-3mer | 0.250 ± 0.006 | 0.276 ± 0.007 | 0.337 ± 0.007 | 0.397 ± 0.007 | 0.473 ± 0.007 | 0.347 ± 0.081 |
| DNABERT-4mer | 0.257 ± 0.006 | 0.279 ± 0.007 | 0.328 ± 0.007 | 0.400 ± 0.007 | 0.524 ± 0.007 | 0.358 ± 0.097 |
| DNABERT-5mer | 0.258 ± 0.006 | 0.296 ± 0.006 | 0.324 ± 0.007 | 0.405 ± 0.007 | 0.513 ± 0.006 | 0.359 ± 0.091 |
| DNABERT-6mer | 0.263 ± 0.006 | 0.299 ± 0.006 | 0.340 ± 0.007 | 0.404 ± 0.007 | 0.526 ± 0.007 | 0.366 ± 0.093 |
| DNABERT2 | 0.311 ± 0.007 | 0.367 ± 0.007 | 0.443 ± 0.006 | 0.512 ± 0.007 | 0.618 ± 0.006 | 0.450 ± 0.108 |
| Gena-LM-Base | 0.305 ± 0.007 | 0.328 ± 0.006 | 0.406 ± 0.007 | 0.504 ± 0.007 | 0.607 ± 0.007 | 0.430 ± 0.113 |
| Gena-LM-Base-T2T | 0.303 ± 0.007 | 0.344 ± 0.006 | 0.409 ± 0.007 | 0.516 ± 0.007 | 0.632 ± 0.006 | 0.441 ± 0.120 |
| Gena-LM-Base-LastIn-T2T | 0.304 ± 0.007 | 0.351 ± 0.007 | 0.429 ± 0.007 | 0.504 ± 0.007 | 0.637 ± 0.006 | 0.445 ± 0.118 |
| **Gena-LM-Base-T2T-Multi** | **0.322 ± 0.007** | **0.373 ± 0.007** | 0.444 ± 0.007 | 0.530 ± 0.007 | **0.641 ± 0.006** | **0.462 ± 0.114** |
| Gena-LM-Large | 0.308 ± 0.007 | 0.347 ± 0.007 | 0.410 ± 0.007 | 0.509 ± 0.007 | 0.608 ± 0.006 | 0.437 ± 0.109 |
| Gena-LM-BigBird | 0.296 ± 0.007 | 0.356 ± 0.007 | 0.423 ± 0.007 | 0.515 ± 0.007 | 0.604 ± 0.006 | 0.439 ± 0.110 |
| Generator-1.2b | 0.316 ± 0.007 | 0.368 ± 0.007 | **0.447 ± 0.007** | **0.531 ± 0.007** | 0.626 ± 0.006 | 0.457 ± 0.111 |
| GPN | 0.257 ± 0.006 | 0.298 ± 0.007 | 0.366 ± 0.007 | 0.366 ± 0.007 | 0.374 ± 0.007 | 0.332 ± 0.047 |
| Grover | 0.288 ± 0.006 | 0.335 ± 0.007 | 0.388 ± 0.007 | 0.471 ± 0.007 | 0.600 ± 0.007 | 0.416 ± 0.110 |
| MistralDNA-17m | 0.214 ± 0.006 | 0.239 ± 0.006 | 0.270 ± 0.006 | 0.334 ± 0.007 | 0.445 ± 0.007 | 0.300 ± 0.083 |
| MistralDNA-422m | 0.239 ± 0.006 | 0.285 ± 0.006 | 0.349 ± 0.007 | 0.445 ± 0.007 | 0.544 ± 0.006 | 0.372 ± 0.110 |
| MistralDNA-1.6b | 0.218 ± 0.006 | 0.231 ± 0.006 | 0.266 ± 0.006 | 0.304 ± 0.007 | 0.400 ± 0.007 | 0.284 ± 0.065 |
| NT-500m-1000g | 0.239 ± 0.006 | 0.301 ± 0.007 | 0.369 ± 0.007 | 0.441 ± 0.007 | 0.552 ± 0.007 | 0.380 ± 0.109 |
| NT-500m-Human-Ref | 0.257 ± 0.006 | 0.325 ± 0.007 | 0.366 ± 0.007 | 0.461 ± 0.007 | 0.578 ± 0.006 | 0.398 ± 0.112 |
| NT-2.5b-1000g | 0.295 ± 0.007 | 0.324 ± 0.007 | 0.373 ± 0.007 | 0.471 ± 0.006 | 0.613 ± 0.006 | 0.415 ± 0.116 |
| NT-2.5b-Multi | 0.301 ± 0.006 | 0.372 ± 0.007 | 0.413 ± 0.007 | 0.508 ± 0.006 | 0.632 ± 0.006 | 0.445 ± 0.115 |
| NTv2-50m-Multi | 0.266 ± 0.007 | 0.307 ± 0.006 | 0.378 ± 0.007 | 0.472 ± 0.007 | 0.588 ± 0.006 | 0.402 ± 0.116 |
| NTv2-100m-Multi | 0.260 ± 0.007 | 0.325 ± 0.006 | 0.403 ± 0.007 | 0.501 ± 0.007 | 0.594 ± 0.006 | 0.417 ± 0.120 |
| NTv2-250m-Multi | 0.293 ± 0.007 | 0.367 ± 0.006 | 0.437 ± 0.007 | 0.526 ± 0.007 | 0.622 ± 0.006 | 0.449 ± 0.116 |
| NTv2-500m-Multi | 0.281 ± 0.006 | 0.355 ± 0.007 | 0.421 ± 0.007 | 0.516 ± 0.007 | 0.620 ± 0.006 | 0.439 ± 0.119 |
| Caduceus-Ph | 0.315 ± 0.007 | 0.360 ± 0.007 | 0.442 ± 0.007 | 0.503 ± 0.006 | 0.616 ± 0.006 | 0.447 ± 0.107 |
| Caduceus-Ps | 0.300 ± 0.007 | 0.340 ± 0.007 | 0.426 ± 0.007 | 0.508 ± 0.006 | 0.610 ± 0.006 | 0.437 ± 0.113 |
| Hyena-Large | 0.263 ± 0.007 | 0.292 ± 0.007 | 0.355 ± 0.007 | 0.413 ± 0.007 | 0.560 ± 0.007 | 0.377 ± 0.105 |
| Hyena-Medium | 0.262 ± 0.006 | 0.299 ± 0.007 | 0.353 ± 0.007 | 0.431 ± 0.007 | 0.545 ± 0.007 | 0.378 ± 0.101 |
| RNAErnie | 0.173 ± 0.005 | 0.199 ± 0.006 | 0.197 ± 0.005 | 0.198 ± 0.006 | 0.197 ± 0.006 | 0.193 ± 0.010 |
| RNAFM | 0.298 ± 0.007 | 0.334 ± 0.007 | 0.389 ± 0.007 | 0.381 ± 0.007 | 0.394 ± 0.007 | 0.360 ± 0.037 |
| SpliceBERT | 0.301 ± 0.006 | 0.359 ± 0.007 | 0.417 ± 0.006 | 0.500 ± 0.007 | 0.504 ± 0.006 | 0.416 ± 0.079 |

#### Regulatory Mechanism Modeling

**Regulatory Activity Prediction** centers on predicting the quantitative levels of chromatin accessibility, specifically through DNase-Seq read counts. DNase-Seq is a method used to map regions of open chromatin, which are critical for understanding gene regulation, as these regions are accessible to transcription factors and other regulatory proteins^31^. By predicting chromatin accessibility from DNA sequence data, this task provides insights into the regulatory landscape of the genome. Understanding chromatin accessibility is essential for identifying key regulatory regions involved in gene expression, cell differentiation, and disease mechanisms. As shown in Fig. 4a, in the regulatory activity prediction task, performance varies widely across models. NTv2-500m-Multi achieves the highest Spearman score of 0.782, closely followed by NT-2.5b-Multi (0.774) and NTv2-250m-Multi (0.764), indicating strong capacity for modeling regulatory activity. However, a substantial performance drop is observed among lower-ranked models, with RNAErnie (0.134), GPN (0.266), and RNAFM (0.284) performing poorly (Table 10). This wide spread highlights the difficulty of the task and the importance of model architecture and pretraining strategies in capturing quantitative regulatory patterns.

**Figure 4.**
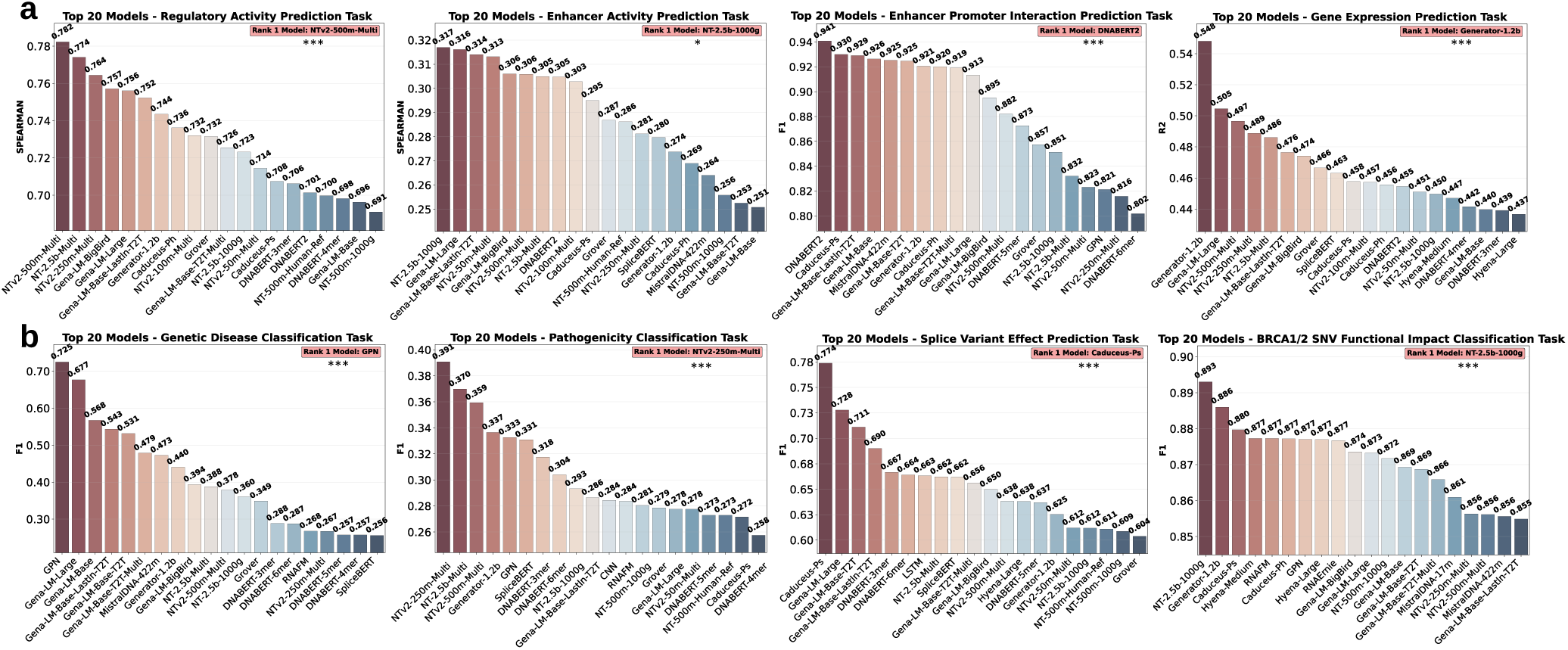
Model Performance on Regulatory Mechanism Modeling Tasks and Genetic Variant Effect Prediction Tasks. **a**, Top-20 models on Regulatory Mechanism Modeling tasks, including regulatory activity prediction, enhancer activity prediction, enhancer–promoter interaction prediction, and gene expression prediction (***: *P <* 0.001). **b**, Performance on Genetic Variant Effect Prediction tasks, including genetic disease classification, pathogenicity classification, splice variant effect prediction, and BRCA1/2 SNV function impact classification (***: *P <* 0.001).

**Table 10.**
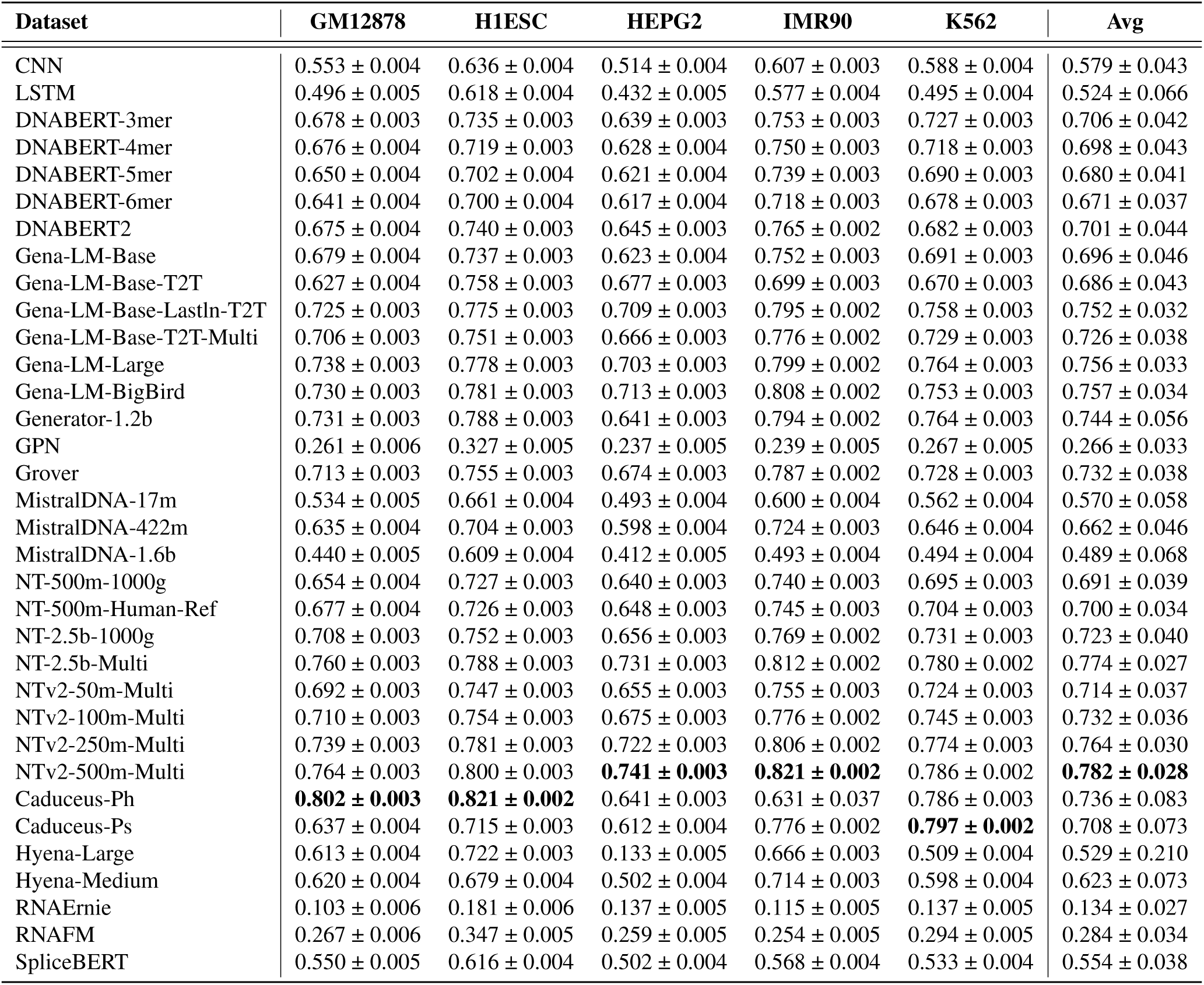
Spearman score of the evaluated models on Regulatory Activity Prediction.

**Enhancer Activity Prediction** focuses on quantifying the regulatory strength of enhancer regions, which are critical DNA sequences in the human genome that amplify the transcription of target genes. Enhancers play a pivotal role in gene expression by interacting with transcription factors and cofactors, thereby fine-tuning spatial and temporal gene regulation during development and cellular processes. Accurately predicting enhancer activity is biologically significant as it enhances our understanding of the molecular mechanisms driving gene regulation, sheds light on how enhancer dysfunction contributes to diseases such as cancer, and supports the development of targeted therapeutic strategies by identifying key regulatory elements in the genome^32^. As shown in Fig. 4a, in the enhancer activity prediction task, overall performance is modest, with the best model, NT-2.5b-1000g, achieving a Spearman score of 0.317. Other top models including Gena-LM-Large and Gena-LM-Base-Lastln-T2T perform similarly (0.316 and 0.314, respectively). Most models fall within a narrow range between 0.250 and 0.310, indicating limited capacity to explain enhancer activity variance. Notably, as summarized in Table 11, several models such as RNAErnie (0.114) and CNN (0.100) perform poorly, suggesting that enhancer activity prediction remains a challenging regression task requiring more specialized modeling of regulatory sequence features.

**Table 11.**
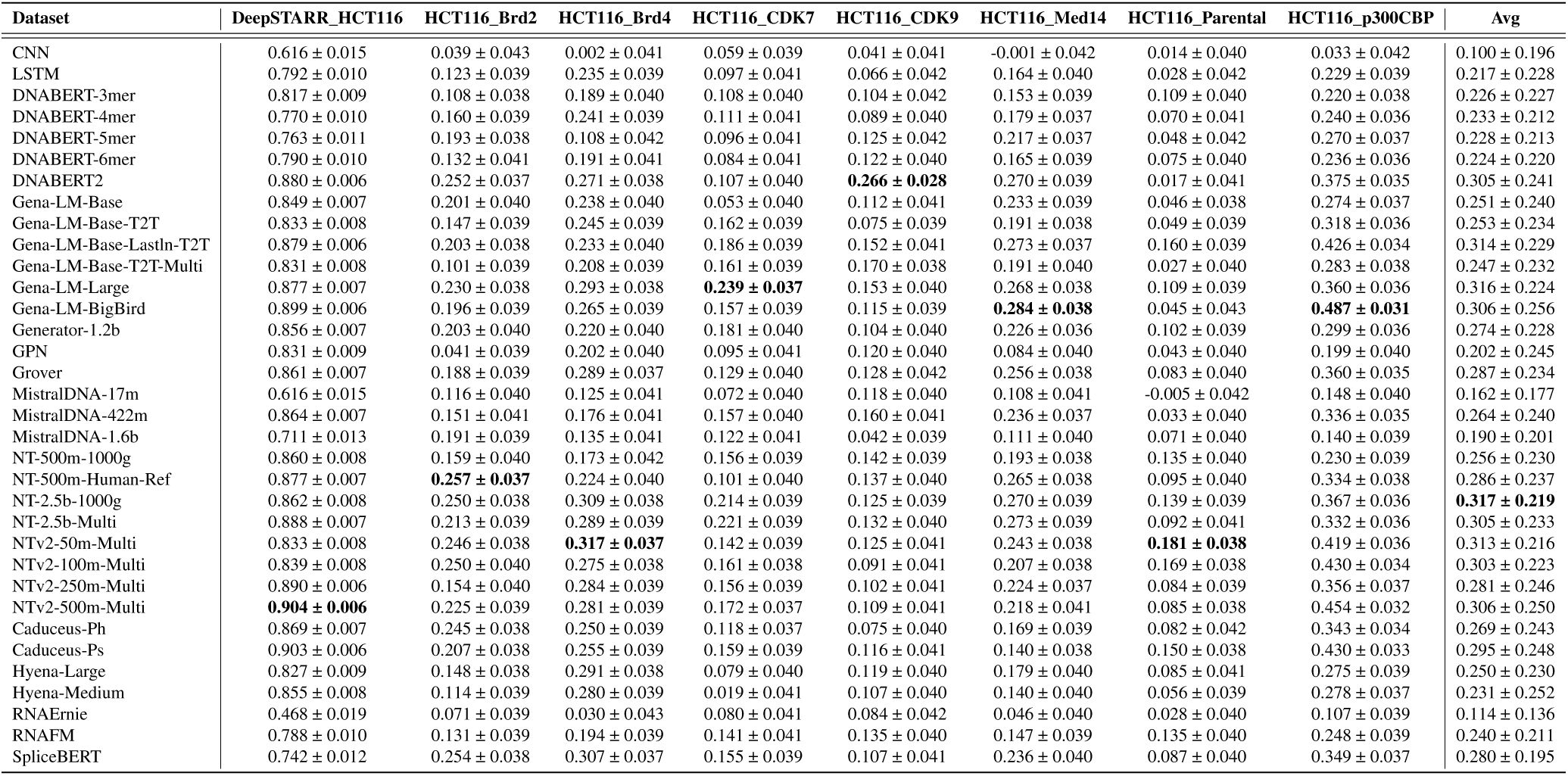
Spearman score of the evaluated models on Enhancer Activity Prediction.

| Dataset | DeepSTARR_HCT116 | HCT116_Brd2 | HCT116_Brd4 | HCT116_CDK7 | HCT116_CDK9 | HCT116_Med14 | HCT116_Parental | HCT116_p300CBP | Avg |
| --- | --- | --- | --- | --- | --- | --- | --- | --- | --- |
| CNN | 0.616 ± 0.015 | 0.039 ± 0.043 | 0.002 ± 0.041 | 0.059 ± 0.039 | 0.041 ± 0.041 | -0.001 ± 0.042 | 0.014 ± 0.040 | 0.033 ± 0.042 | 0.100 ± 0.196 |
| LSTM | 0.792 ± 0.010 | 0.123 ± 0.039 | 0.235 ± 0.039 | 0.097 ± 0.041 | 0.066 ± 0.042 | 0.164 ± 0.040 | 0.028 ± 0.042 | 0.229 ± 0.039 | 0.217 ± 0.228 |
| DNABERT-3mer | 0.817 ± 0.009 | 0.108 ± 0.038 | 0.189 ± 0.040 | 0.108 ± 0.040 | 0.104 ± 0.042 | 0.153 ± 0.039 | 0.109 ± 0.040 | 0.220 ± 0.038 | 0.226 ± 0.227 |
| DNABERT-4mer | 0.770 ± 0.010 | 0.160 ± 0.039 | 0.241 ± 0.039 | 0.111 ± 0.041 | 0.089 ± 0.040 | 0.179 ± 0.037 | 0.070 ± 0.041 | 0.240 ± 0.036 | 0.233 ± 0.212 |
| DNABERT-5mer | 0.763 ± 0.011 | 0.193 ± 0.038 | 0.108 ± 0.042 | 0.096 ± 0.041 | 0.125 ± 0.042 | 0.217 ± 0.037 | 0.048 ± 0.042 | 0.270 ± 0.037 | 0.228 ± 0.213 |
| DNABERT-6mer | 0.790 ± 0.010 | 0.132 ± 0.041 | 0.191 ± 0.041 | 0.084 ± 0.041 | 0.122 ± 0.040 | 0.165 ± 0.039 | 0.075 ± 0.040 | 0.236 ± 0.036 | 0.224 ± 0.220 |
| DNABERT2 | 0.880 ± 0.006 | 0.252 ± 0.037 | 0.271 ± 0.038 | 0.107 ± 0.040 | **0.266 ± 0.028** | 0.270 ± 0.039 | 0.017 ± 0.041 | 0.375 ± 0.035 | 0.305 ± 0.241 |
| Gena-LM-Base | 0.849 ± 0.007 | 0.201 ± 0.040 | 0.238 ± 0.040 | 0.053 ± 0.040 | 0.112 ± 0.041 | 0.233 ± 0.039 | 0.046 ± 0.038 | 0.274 ± 0.037 | 0.251 ± 0.240 |
| Gena-LM-Base-T2T | 0.833 ± 0.008 | 0.147 ± 0.039 | 0.245 ± 0.039 | 0.162 ± 0.039 | 0.075 ± 0.039 | 0.191 ± 0.038 | 0.049 ± 0.039 | 0.318 ± 0.036 | 0.253 ± 0.234 |
| Gena-LM-Base-LastIn-T2T | 0.879 ± 0.006 | 0.203 ± 0.038 | 0.233 ± 0.040 | 0.186 ± 0.039 | 0.152 ± 0.041 | 0.273 ± 0.037 | 0.160 ± 0.039 | 0.426 ± 0.034 | 0.314 ± 0.229 |
| Gena-LM-Base-T2T-Multi | 0.831 ± 0.008 | 0.101 ± 0.039 | 0.208 ± 0.039 | 0.161 ± 0.039 | 0.170 ± 0.038 | 0.191 ± 0.040 | 0.027 ± 0.040 | 0.283 ± 0.038 | 0.247 ± 0.232 |
| Gena-LM-Large | 0.877 ± 0.007 | 0.230 ± 0.038 | 0.293 ± 0.038 | **0.239 ± 0.037** | 0.153 ± 0.040 | 0.268 ± 0.038 | 0.109 ± 0.039 | 0.360 ± 0.036 | 0.316 ± 0.224 |
| Gena-LM-BigBird | 0.899 ± 0.006 | 0.196 ± 0.039 | 0.265 ± 0.039 | 0.157 ± 0.039 | 0.115 ± 0.039 | **0.284 ± 0.038** | 0.045 ± 0.043 | **0.487 ± 0.031** | 0.306 ± 0.256 |
| Generator-1.2b | 0.856 ± 0.007 | 0.203 ± 0.040 | 0.220 ± 0.040 | 0.181 ± 0.040 | 0.104 ± 0.040 | 0.226 ± 0.036 | 0.102 ± 0.039 | 0.299 ± 0.036 | 0.274 ± 0.228 |
| GPN | 0.831 ± 0.009 | 0.041 ± 0.039 | 0.202 ± 0.040 | 0.095 ± 0.041 | 0.120 ± 0.040 | 0.084 ± 0.040 | 0.043 ± 0.040 | 0.199 ± 0.040 | 0.202 ± 0.245 |
| Grover | 0.861 ± 0.007 | 0.188 ± 0.039 | 0.289 ± 0.037 | 0.129 ± 0.040 | 0.128 ± 0.042 | 0.256 ± 0.038 | 0.083 ± 0.040 | 0.360 ± 0.035 | 0.287 ± 0.234 |
| MistralDNA-17m | 0.616 ± 0.015 | 0.116 ± 0.040 | 0.125 ± 0.041 | 0.072 ± 0.040 | 0.118 ± 0.040 | 0.108 ± 0.041 | -0.005 ± 0.042 | 0.148 ± 0.040 | 0.162 ± 0.177 |
| MistralDNA-422m | 0.864 ± 0.007 | 0.151 ± 0.041 | 0.176 ± 0.041 | 0.157 ± 0.040 | 0.160 ± 0.041 | 0.236 ± 0.037 | 0.033 ± 0.040 | 0.336 ± 0.035 | 0.264 ± 0.240 |
| MistralDNA-1.6b | 0.711 ± 0.013 | 0.191 ± 0.039 | 0.135 ± 0.041 | 0.122 ± 0.041 | 0.042 ± 0.039 | 0.111 ± 0.040 | 0.071 ± 0.040 | 0.140 ± 0.039 | 0.190 ± 0.201 |
| NT-500m-1000g | 0.860 ± 0.008 | 0.159 ± 0.040 | 0.173 ± 0.042 | 0.156 ± 0.039 | 0.142 ± 0.039 | 0.193 ± 0.038 | 0.135 ± 0.040 | 0.230 ± 0.039 | 0.256 ± 0.230 |
| NT-500m-Human-Ref | 0.877 ± 0.007 | **0.257 ± 0.037** | 0.224 ± 0.040 | 0.101 ± 0.040 | 0.137 ± 0.040 | 0.265 ± 0.038 | 0.095 ± 0.040 | 0.334 ± 0.038 | 0.286 ± 0.237 |
| NT-2.5b-1000g | 0.862 ± 0.008 | 0.250 ± 0.038 | 0.309 ± 0.038 | 0.214 ± 0.039 | 0.125 ± 0.039 | 0.270 ± 0.039 | 0.139 ± 0.039 | 0.367 ± 0.036 | **0.317 ± 0.219** |
| NT-2.5b-Multi | 0.888 ± 0.007 | 0.213 ± 0.039 | 0.289 ± 0.039 | 0.221 ± 0.039 | 0.132 ± 0.040 | 0.273 ± 0.039 | 0.092 ± 0.041 | 0.332 ± 0.036 | 0.305 ± 0.233 |
| NTv2-50m-Multi | 0.833 ± 0.008 | 0.246 ± 0.038 | **0.317 ± 0.037** | 0.142 ± 0.039 | 0.125 ± 0.041 | 0.243 ± 0.038 | **0.181 ± 0.038** | 0.419 ± 0.036 | 0.313 ± 0.216 |
| NTv2-100m-Multi | 0.839 ± 0.008 | 0.250 ± 0.040 | 0.275 ± 0.038 | 0.161 ± 0.038 | 0.091 ± 0.041 | 0.207 ± 0.038 | 0.169 ± 0.038 | 0.430 ± 0.034 | 0.303 ± 0.223 |
| NTv2-250m-Multi | 0.890 ± 0.006 | 0.154 ± 0.040 | 0.284 ± 0.039 | 0.156 ± 0.039 | 0.102 ± 0.041 | 0.224 ± 0.037 | 0.084 ± 0.039 | 0.356 ± 0.037 | 0.281 ± 0.246 |
| NTv2-500m-Multi | **0.904 ± 0.006** | 0.225 ± 0.039 | 0.281 ± 0.039 | 0.172 ± 0.037 | 0.109 ± 0.041 | 0.218 ± 0.041 | 0.085 ± 0.038 | 0.454 ± 0.032 | 0.306 ± 0.250 |
| Caduceus-Ph | 0.869 ± 0.007 | 0.245 ± 0.038 | 0.250 ± 0.039 | 0.118 ± 0.037 | 0.075 ± 0.040 | 0.169 ± 0.039 | 0.082 ± 0.042 | 0.343 ± 0.034 | 0.269 ± 0.243 |
| Caduceus-Ps | 0.903 ± 0.006 | 0.207 ± 0.038 | 0.255 ± 0.039 | 0.159 ± 0.039 | 0.116 ± 0.041 | 0.140 ± 0.038 | 0.150 ± 0.038 | 0.430 ± 0.033 | 0.295 ± 0.248 |
| Hyena-Large | 0.827 ± 0.009 | 0.148 ± 0.038 | 0.291 ± 0.038 | 0.079 ± 0.040 | 0.119 ± 0.040 | 0.179 ± 0.040 | 0.085 ± 0.041 | 0.275 ± 0.039 | 0.250 ± 0.230 |
| Hyena-Medium | 0.855 ± 0.008 | 0.114 ± 0.039 | 0.280 ± 0.039 | 0.019 ± 0.041 | 0.107 ± 0.040 | 0.140 ± 0.040 | 0.056 ± 0.039 | 0.278 ± 0.037 | 0.231 ± 0.252 |
| RNAErnie | 0.468 ± 0.019 | 0.071 ± 0.039 | 0.030 ± 0.043 | 0.080 ± 0.041 | 0.084 ± 0.042 | 0.046 ± 0.040 | 0.028 ± 0.040 | 0.107 ± 0.039 | 0.114 ± 0.136 |
| RNAFM | 0.788 ± 0.010 | 0.131 ± 0.039 | 0.194 ± 0.039 | 0.141 ± 0.041 | 0.135 ± 0.040 | 0.147 ± 0.039 | 0.135 ± 0.040 | 0.248 ± 0.039 | 0.240 ± 0.211 |
| SpliceBERT | 0.742 ± 0.012 | 0.254 ± 0.038 | 0.307 ± 0.037 | 0.155 ± 0.039 | 0.107 ± 0.041 | 0.236 ± 0.040 | 0.087 ± 0.040 | 0.349 ± 0.037 | 0.280 ± 0.195 |

**Enhancer Promoter Interaction Prediction**. Enhancer–promoter interactions lie at the core of gene regulation: as distal regulatory elements, enhancers make physical contact with promoters through three-dimensional chromatin structures, thereby influencing when, where, and how strongly genes are transcribed^33^. Because multiple enhancers can target the same promoter, and one enhancer can simultaneously control different genes, those interactions are highly complex and cell-type-specific. Consequently, identifying and understanding enhancer–promoter interactions is pivotal for deciphering gene regulatory networks and unraveling disease-related molecular mechanisms^34^. As shown in Fig. 4a, in the enhancer–promoter interaction prediction task, most models achieve high F1 score, with DNABERT2 leading at 0.941, followed closely by Caduceus-Ps (0.930) and Gena-LM-Base-Lastln-T2T (0.929). A large group of models, including MistralDNA-422m, Generator-1.2b, and Gena-LM-Large, perform above 0.900, indicating that many gLMs are well-suited for this task. However, as indicated in Table 12, lower-performing models such as RNAErnie (0.525) and Hyena-Medium (0.394) show a substantial gap, highlighting that model architecture and pretraining remain critical for achieving robust interaction modeling.

**Table 12.** F1 score of the evaluated models on Enhancer Promoter Interaction Prediction.

| Dataset | GM12878 | HUVEC | HeLa-S3 | IMR90 | K562 | NHEK | Avg |
| --- | --- | --- | --- | --- | --- | --- | --- |
| CNN | 0.613 ± 0.011 | 0.609 ± 0.011 | 0.642 ± 0.011 | 0.656 ± 0.011 | 0.579 ± 0.011 | 0.662 ± 0.011 | 0.627 ± 0.029 |
| LSTM | 0.594 ± 0.012 | 0.707 ± 0.010 | 0.763 ± 0.009 | 0.834 ± 0.008 | 0.755 ± 0.010 | 0.745 ± 0.010 | 0.733 ± 0.073 |
| DNABERT-3mer | 0.651 ± 0.011 | 0.710 ± 0.010 | 0.642 ± 0.011 | 0.840 ± 0.008 | 0.701 ± 0.010 | 0.674 ± 0.011 | 0.703 ± 0.066 |
| DNABERT-4mer | 0.655 ± 0.011 | 0.744 ± 0.010 | 0.724 ± 0.010 | 0.790 ± 0.009 | 0.729 ± 0.010 | 0.896 ± 0.007 | 0.756 ± 0.074 |
| DNABERT-5mer | 0.814 ± 0.008 | 0.868 ± 0.008 | 0.886 ± 0.007 | 0.860 ± 0.008 | 0.870 ± 0.008 | 0.938 ± 0.005 | 0.873 ± 0.037 |
| DNABERT-6mer | 0.735 ± 0.010 | 0.803 ± 0.009 | 0.804 ± 0.009 | 0.824 ± 0.009 | 0.761 ± 0.009 | 0.885 ± 0.007 | 0.802 ± 0.047 |
| DNABERT2 | 0.916 ± 0.006 | 0.925 ± 0.006 | 0.954 ± 0.005 | 0.963 ± 0.004 | **0.933 ± 0.006** | 0.955 ± 0.005 | **0.941 ± 0.017** |
| Gena-LM-Base | 0.855 ± 0.008 | 0.919 ± 0.006 | 0.945 ± 0.005 | 0.975 ± 0.003 | 0.885 ± 0.007 | **0.980 ± 0.003** | 0.926 ± 0.046 |
| Gena-LM-Base-T2T | 0.882 ± 0.007 | 0.927 ± 0.006 | 0.941 ± 0.005 | 0.930 ± 0.006 | 0.891 ± 0.007 | 0.977 ± 0.003 | 0.925 ± 0.032 |
| Gena-LM-Base-LastIn-T2T | 0.885 ± 0.007 | 0.931 ± 0.006 | 0.945 ± 0.005 | 0.960 ± 0.004 | 0.885 ± 0.007 | 0.969 ± 0.004 | 0.929 ± 0.033 |
| Gena-LM-Base-T2T-Multi | 0.897 ± 0.007 | 0.912 ± 0.006 | 0.940 ± 0.005 | 0.912 ± 0.007 | 0.882 ± 0.007 | 0.972 ± 0.004 | 0.919 ± 0.029 |
| Gena-LM-Large | 0.842 ± 0.008 | 0.928 ± 0.006 | 0.937 ± 0.006 | 0.948 ± 0.005 | 0.856 ± 0.008 | 0.968 ± 0.004 | 0.913 ± 0.047 |
| Gena-LM-BigBird | 0.809 ± 0.009 | 0.913 ± 0.006 | 0.935 ± 0.006 | 0.910 ± 0.006 | 0.875 ± 0.007 | 0.929 ± 0.006 | 0.895 ± 0.043 |
| Generator-1.2b | 0.889 ± 0.007 | 0.931 ± 0.005 | 0.919 ± 0.006 | 0.912 ± 0.006 | 0.912 ± 0.006 | 0.960 ± 0.004 | 0.921 ± 0.022 |
| GPN | 0.777 ± 0.009 | 0.792 ± 0.009 | 0.817 ± 0.009 | 0.839 ± 0.008 | 0.812 ± 0.009 | 0.891 ± 0.007 | 0.821 ± 0.037 |
| Grover | 0.848 ± 0.008 | 0.877 ± 0.007 | 0.801 ± 0.009 | 0.849 ± 0.008 | 0.829 ± 0.008 | 0.941 ± 0.005 | 0.857 ± 0.044 |
| MistralDNA-17m | 0.633 ± 0.011 | 0.688 ± 0.010 | 0.674 ± 0.011 | 0.707 ± 0.010 | 0.622 ± 0.011 | 0.685 ± 0.010 | 0.668 ± 0.031 |
| MistralDNA-422m | 0.881 ± 0.007 | 0.937 ± 0.006 | 0.938 ± 0.005 | 0.939 ± 0.006 | 0.888 ± 0.007 | 0.970 ± 0.004 | 0.925 ± 0.031 |
| MistralDNA-1.6b | 0.754 ± 0.010 | 0.773 ± 0.010 | 0.802 ± 0.009 | 0.811 ± 0.009 | 0.720 ± 0.010 | 0.861 ± 0.008 | 0.787 ± 0.045 |
| NT-500m-1000g | 0.743 ± 0.010 | 0.745 ± 0.010 | 0.766 ± 0.010 | 0.851 ± 0.008 | 0.685 ± 0.011 | 0.830 ± 0.008 | 0.770 ± 0.056 |
| NT-500m-Human-Ref | 0.698 ± 0.010 | 0.810 ± 0.009 | 0.783 ± 0.009 | 0.855 ± 0.008 | 0.767 ± 0.010 | 0.760 ± 0.010 | 0.779 ± 0.048 |
| NT-2.5b-1000g | 0.709 ± 0.010 | 0.808 ± 0.009 | 0.881 ± 0.008 | 0.912 ± 0.006 | 0.860 ± 0.008 | 0.938 ± 0.005 | 0.851 ± 0.076 |
| NT-2.5b-Multi | 0.746 ± 0.010 | 0.798 ± 0.009 | 0.899 ± 0.007 | 0.878 ± 0.007 | 0.749 ± 0.010 | 0.924 ± 0.006 | 0.832 ± 0.071 |
| NTv2-50m-Multi | 0.786 ± 0.009 | 0.843 ± 0.008 | 0.791 ± 0.010 | 0.830 ± 0.009 | 0.818 ± 0.009 | 0.871 ± 0.007 | 0.823 ± 0.029 |
| NTv2-100m-Multi | 0.812 ± 0.009 | 0.769 ± 0.009 | 0.763 ± 0.010 | 0.807 ± 0.009 | 0.778 ± 0.009 | 0.824 ± 0.009 | 0.792 ± 0.023 |
| NTv2-250m-Multi | 0.794 ± 0.009 | 0.750 ± 0.010 | 0.822 ± 0.009 | 0.846 ± 0.008 | 0.810 ± 0.009 | 0.872 ± 0.008 | 0.816 ± 0.038 |
| NTv2-500m-Multi | 0.805 ± 0.009 | 0.868 ± 0.008 | 0.882 ± 0.007 | 0.945 ± 0.005 | 0.842 ± 0.008 | 0.951 ± 0.005 | 0.882 ± 0.053 |
| Caduceus-Ph | 0.864 ± 0.008 | **0.962 ± 0.004** | **0.965 ± 0.004** | **0.982 ± 0.003** | 0.806 ± 0.009 | 0.943 ± 0.005 | 0.920 ± 0.064 |
| Caduceus-Ps | **0.916 ± 0.006** | 0.953 ± 0.005 | 0.963 ± 0.004 | 0.974 ± 0.004 | 0.796 ± 0.009 | 0.978 ± 0.003 | 0.930 ± 0.063 |
| Hyena-Large | 0.636 ± 0.011 | 0.718 ± 0.010 | 0.669 ± 0.011 | 0.719 ± 0.010 | 0.664 ± 0.011 | 0.633 ± 0.011 | 0.673 ± 0.035 |
| Hyena-Medium | 0.464 ± 0.011 | 0.338 ± 0.005 | 0.342 ± 0.005 | 0.343 ± 0.005 | 0.532 ± 0.011 | 0.342 ± 0.005 | 0.394 ± 0.076 |
| RNAErnie | 0.512 ± 0.011 | 0.544 ± 0.011 | 0.494 ± 0.011 | 0.538 ± 0.011 | 0.524 ± 0.011 | 0.542 ± 0.011 | 0.525 ± 0.018 |
| RNAFM | 0.618 ± 0.011 | 0.673 ± 0.010 | 0.668 ± 0.011 | 0.705 ± 0.010 | 0.621 ± 0.011 | 0.676 ± 0.011 | 0.660 ± 0.031 |
| SpliceBERT | 0.738 ± 0.010 | 0.761 ± 0.010 | 0.769 ± 0.010 | 0.744 ± 0.010 | 0.664 ± 0.011 | 0.774 ± 0.009 | 0.742 ± 0.037 |

**Gene Expression Prediction** seeks to model the relationship between DNA sequence and gene activity levels, revealing how genomic elements regulate transcription. This task is biologically meaningful because gene expression underlies cellular identity, development, and response to environmental cues. Accurate prediction of expression levels from sequence can uncover cis-regulatory logic, aid in interpreting non-coding variants, and support studies in disease genomics and precision medicine^35^. It also contributes to a deeper understanding of how genetic variation influences phenotypic diversity and cellular function. As shown in Fig. 4a, in the gene expression prediction task, overall performance is moderate. Generator-1.2b set the performance ceiling with an *R*^2^ of 0.548, substantially outperforming the next-best models, Gena-LM-Large (0.505) and NTv2-500m-Multi (0.497). Most models cluster between 0.430 and 0.480, suggesting comparable effectiveness in capturing expression-related signals. However, several models—including RNAErnie (0.295) and MistralDNA-1.6b (0.354)—exhibit significantly lower performance (Table 13), indicating that modeling gene expression remains a challenging regression task that may require deeper integration of regulatory and contextual sequence information.

**Table 13.**
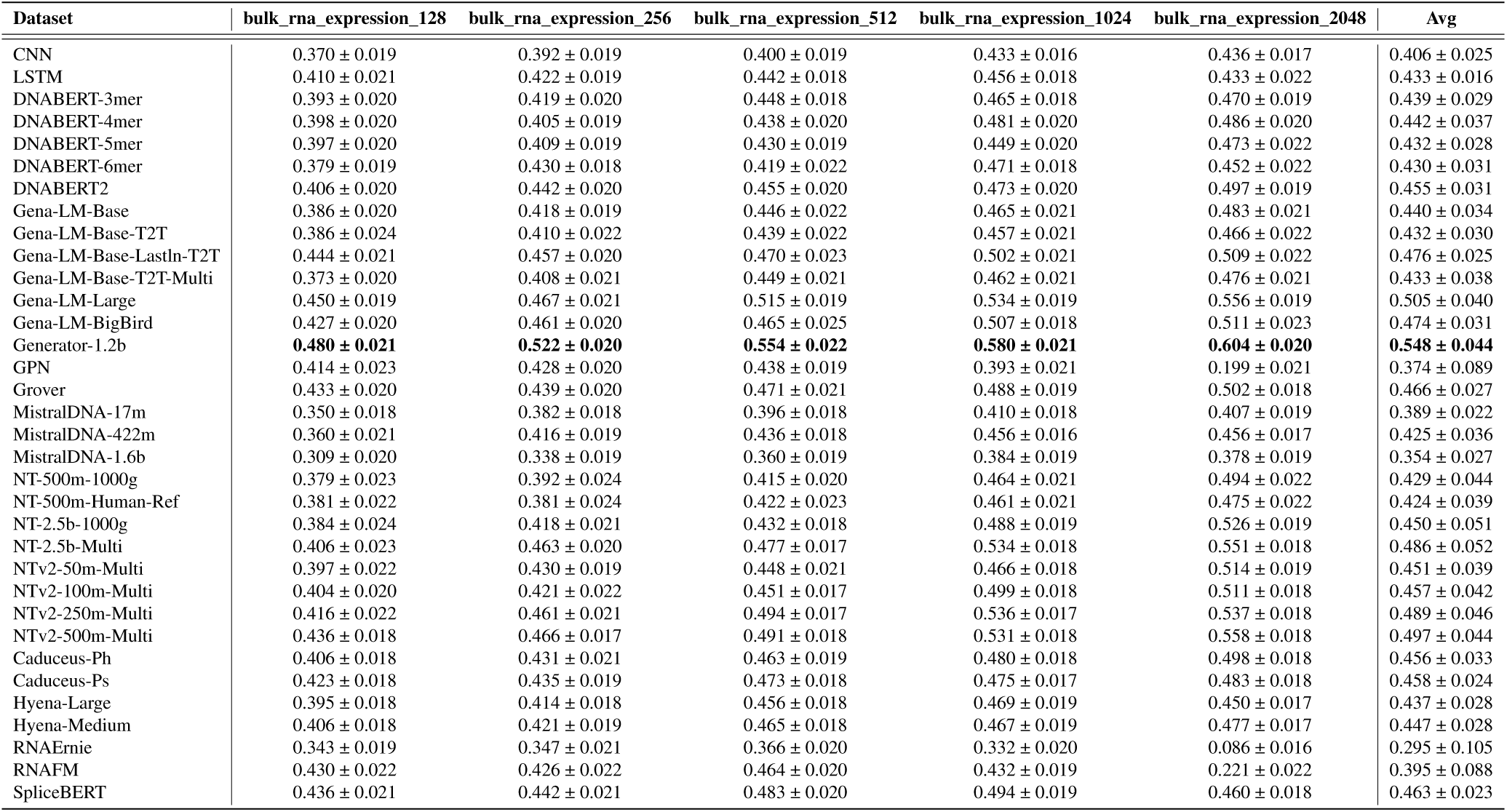
***R*^2^ score of the evaluated models on Gene Expression Prediction**

#### Genetic Variant Effect Prediction

**Genetic Disease Classification**. We first investigated the ability of gLMs to link specific genetic variants to distinct disease categories. This task relies on curated single nucleotide variants (SNVs) from ClinVar^36^ and utilizes sequences centered on the variant sites, enabling the models to learn predictive sequence-level features. As shown in Fig. 4b, in the genetic disease classification task, GPN significantly outperforms all other models with an F1 score of 0.725, followed by Gena-LM-Large (0.677) and Gena-LM-Base (0.568). The performance gap between the top and bottom models is substantial, with DNABERT-3mer and DNABERT-6mer achieving only 0.288 and 0.287 (Table 14), respectively. This wide disparity highlights the importance of model architecture and pretraining design in capturing disease-relevant sequence features, and suggests that many existing gLMs still struggle with complex genotype-to-phenotype classification tasks.

**Table 14.** F1 score of the evaluated models on Genetic Disease Classification.

| Dataset | breast_disease_type_classification | cardiovascular_disorders_classification | kidney_disease_type_classification | Avg |
| --- | --- | --- | --- | --- |
| CNN | 0.237 ± 0.009 | 0.106 ± 0.004 | 0.146 ± 0.008 | 0.163 ± 0.055 |
| LSTM | 0.267 ± 0.010 | 0.137 ± 0.003 | 0.123 ± 0.005 | 0.176 ± 0.065 |
| DNABERT-3mer | 0.366 ± 0.014 | 0.235 ± 0.005 | 0.264 ± 0.013 | 0.288 ± 0.056 |
| DNABERT-4mer | 0.234 ± 0.010 | 0.276 ± 0.008 | 0.262 ± 0.016 | 0.257 ± 0.018 |
| DNABERT-5mer | 0.304 ± 0.013 | 0.236 ± 0.005 | 0.232 ± 0.011 | 0.257 ± 0.033 |
| DNABERT-6mer | 0.360 ± 0.014 | 0.223 ± 0.005 | 0.279 ± 0.013 | 0.287 ± 0.056 |
| DNABERT2 | 0.192 ± 0.006 | 0.061 ± 0.002 | 0.094 ± 0.002 | 0.116 ± 0.056 |
| Gena-LM-Base | 0.483 ± 0.010 | 0.645 ± 0.009 | 0.574 ± 0.018 | 0.568 ± 0.066 |
| Gena-LM-Base-T2T | 0.467 ± 0.013 | 0.576 ± 0.010 | 0.551 ± 0.014 | 0.531 ± 0.046 |
| Gena-LM-Base-LastIn-T2T | 0.500 ± 0.012 | 0.603 ± 0.009 | 0.527 ± 0.015 | 0.543 ± 0.043 |
| Gena-LM-Base-T2T-Multi | 0.424 ± 0.012 | 0.502 ± 0.009 | 0.510 ± 0.017 | 0.479 ± 0.039 |
| Gena-LM-Large | **0.558 ± 0.014** | 0.790 ± 0.008 | 0.683 ± 0.019 | 0.677 ± 0.095 |
| Gena-LM-BigBird | 0.515 ± 0.013 | 0.045 ± 0.001 | 0.621 ± 0.017 | 0.394 ± 0.250 |
| Generator-1.2b | 0.452 ± 0.012 | 0.422 ± 0.006 | 0.447 ± 0.013 | 0.440 ± 0.013 |
| GPN | 0.519 ± 0.011 | **0.812 ± 0.008** | **0.845 ± 0.015** | **0.725 ± 0.146** |
| Grover | 0.431 ± 0.012 | 0.376 ± 0.008 | 0.240 ± 0.012 | 0.349 ± 0.080 |
| MistralDNA-17m | 0.291 ± 0.006 | 0.141 ± 0.005 | 0.107 ± 0.008 | 0.180 ± 0.080 |
| MistralDNA-422m | 0.413 ± 0.011 | 0.526 ± 0.009 | 0.479 ± 0.012 | 0.473 ± 0.046 |
| MistralDNA-1.6b | 0.311 ± 0.007 | 0.150 ± 0.004 | 0.185 ± 0.010 | 0.216 ± 0.069 |
| NT-500m-1000g | 0.357 ± 0.011 | 0.149 ± 0.004 | 0.253 ± 0.014 | 0.253 ± 0.085 |
| NT-500m-Human-Ref | 0.401 ± 0.012 | 0.188 ± 0.005 | 0.172 ± 0.008 | 0.254 ± 0.104 |
| NT-2.5b-1000g | 0.381 ± 0.011 | 0.359 ± 0.007 | 0.341 ± 0.013 | 0.360 ± 0.016 |
| NT-2.5b-Multi | 0.435 ± 0.012 | 0.391 ± 0.007 | 0.338 ± 0.014 | 0.388 ± 0.039 |
| NTv2-50m-Multi | 0.318 ± 0.009 | 0.234 ± 0.005 | 0.151 ± 0.011 | 0.234 ± 0.068 |
| NTv2-100m-Multi | 0.355 ± 0.009 | 0.231 ± 0.005 | 0.153 ± 0.011 | 0.246 ± 0.083 |
| NTv2-250m-Multi | 0.379 ± 0.011 | 0.265 ± 0.005 | 0.156 ± 0.007 | 0.267 ± 0.091 |
| NTv2-500m-Multi | 0.377 ± 0.010 | 0.328 ± 0.006 | 0.430 ± 0.015 | 0.378 ± 0.042 |
| Caduceus-Ph | 0.141 ± 0.003 | 0.043 ± 0.001 | 0.094 ± 0.002 | 0.093 ± 0.040 |
| Caduceus-Ps | 0.238 ± 0.009 | 0.121 ± 0.003 | 0.091 ± 0.002 | 0.150 ± 0.063 |
| Hyena-Large | 0.141 ± 0.003 | 0.115 ± 0.003 | 0.094 ± 0.002 | 0.117 ± 0.019 |
| Hyena-Medium | 0.141 ± 0.003 | 0.116 ± 0.004 | 0.094 ± 0.002 | 0.117 ± 0.019 |
| RNAErnie | 0.250 ± 0.008 | 0.173 ± 0.005 | 0.214 ± 0.014 | 0.212 ± 0.032 |
| RNAFM | 0.329 ± 0.012 | 0.243 ± 0.006 | 0.231 ± 0.012 | 0.268 ± 0.043 |
| SpliceBERT | 0.294 ± 0.010 | 0.262 ± 0.006 | 0.212 ± 0.010 | 0.256 ± 0.034 |

**Pathogenicity Classification**. In addition to disease classification, we also evaluated the performance of gLMs to predict the pathogenic potential of genetic variants by categorizing them into four groups—likely benign, benign, pathogenic, and likely pathogenic—based on their associations with human diseases. Using curated genetic data from ClinVar, our experiments assessed how well the models can identify mutations that may contribute to disease onset and progression. Such evaluations are critical for genetic diagnosis, therapeutic decision making and personalized medicine, as accurately determining a variant’s pathogenicity is essential for understanding its role in disease etiology and selecting appropriate treatment strategies^37^. As shown in Fig. 4b, in the pathogenicity classification task, overall model performance remains relatively low, with the top-performing model NTv2-250m-Multi achieving an F1 score of only 0.391. NT-2.5b-Multi (0.370) and NTv2-500m-Multi (0.359) follow closely, suggesting modest improvements from larger model sizes. The majority of other models exhibited similar or lower performance (Table 15), including Generator-1.2b (0.337) and GPN (0.333). This widespread struggle to reliably distinguish pathogenic from benign variants underscores the need for models better tailored to subtle, variant-level prediction.

**Table 15.** F1 score of the evaluated models on Pathogenicity Classification.

| Dataset | pathogenicity_classification_512 |
| --- | --- |
| CNN | 0.284 ± 0.001 |
| LSTM | 0.237 ± 0.006 |
| DNABERT-3mer | 0.318 ± 0.008 |
| DNABERT-4mer | 0.258 ± 0.007 |
| DNABERT-5mer | 0.273 ± 0.007 |
| DNABERT-6mer | 0.304 ± 0.008 |
| DNABERT2 | 0.213 ± 0.001 |
| Gena-LM-Base | 0.256 ± 0.006 |
| Gena-LM-Base-T2T | 0.235 ± 0.005 |
| Gena-LM-Base-Lastln-T2T | 0.286 ± 0.009 |
| Gena-LM-Base-T2T-Multi | 0.235 ± 0.006 |
| Gena-LM-Large | 0.278 ± 0.008 |
| Gena-LM-BigBird | 0.245 ± 0.005 |
| Generator-1.2b | 0.337 ± 0.007 |
| GPN | 0.333 ± 0.007 |
| Grover | 0.279 ± 0.006 |
| MistralDNA-17m | 0.248 ± 0.005 |
| MistralDNA-422m | 0.253 ± 0.006 |
| MistralDNA-1.6b | 0.221 ± 0.004 |
| NT-500m-1000g | 0.281 ± 0.006 |
| NT-500m-Human-Ref | 0.273 ± 0.006 |
| NT-2.5b-1000g | 0.293 ± 0.006 |
| NT-2.5b-Multi | 0.370 ± 0.008 |
| NTv2-50m-Multi | 0.278 ± 0.006 |
| NTv2-100m-Multi | 0.243 ± 0.005 |
| NTv2-250m-Multi | **0.391 ± 0.008** |
| NTv2-500m-Multi | 0.359 ± 0.008 |
| Caduceus-Ph | 0.213 ± 0.001 |
| Caduceus-Ps | 0.272 ± 0.007 |
| Hyena-Large | 0.235 ± 0.005 |
| Hyena-Medium | 0.223 ± 0.004 |
| RNAErnie | 0.256 ± 0.006 |
| RNAFM | 0.284 ± 0.007 |
| SpliceBERT | 0.331 ± 0.007 |

**Splice Variant Effect Prediction**. Moreover, to assess the performance of gLMs on a broader and more challenging set of functional noncoding variants^38^, we selected the Splice Variant Effect Prediction task from the SpliceVarDB dataset. Splice variants, by altering mRNA splicing patterns, can lead to abnormal transcript structures or protein dysfunction, playing a significant role in genetic disorders and complex diseases^39^. Accurately predicting the effects of splice variants holds profound biological significance, as it elucidates how mutations disrupt gene expression and protein production, offering critical insights into the molecular mechanisms of genetic diseases and supporting the development of diagnostics and therapeutic strategies tailored to individual genomic profiles in precision medicine. As shown in Fig. 4b, in the splice variant effect prediction task, Caduceus-Ps achieves the highest F1 score of 0.774, followed by Gena-LM-Large (0.728) and Gena-LM-Base-T2T (0.711), indicating strong performance from top-tier gLMs. Most other models cluster in the 0.650–0.700 range, including DNABERT variants and LSTM, suggesting moderate but consistent effectiveness. Overall, the task is well handled by several models, though room for further gains remains in capturing subtle splicing impacts of genetic variants.

**BRCA1/2 SNV Functional Impact Classification**. Last but not least, to mitigate curation bias and address incomplete experimental coverage of both coding and noncoding variants, we focused on a dataset that measures the functional consequences of variants across the coding and noncoding regions of the BRCA1 and BRCA2 genes^40,41^. The BRCA1/2 SNV functional impact classification task evaluates the functional effects of SNVs in these critical genes, which play essential roles in DNA repair and maintaining genomic stability. Accurate classification of BRCA1/2 variants is of paramount biological importance, as it reveals how these mutations impair DNA damage repair processes, thereby providing a molecular basis for assessing the risk of hereditary cancers, such as breast and ovarian cancer, and guiding genotype-based strategies for cancer prevention and treatment^42^. Our evaluation showed that most gLMs find this task highly tractable, with F1 scores tightly clustered above 0.860 (Fig. 4b). The best-performing model, NT-2.5b-1000g, achieves an F1 of 0.893, followed by Generator-1.2b (0.886) and Caduceus-Ps (0.880). The performance difference between the top and bottom models is minimal (only 2.7%), indicating that this task is well-handled by most gLMs and may not be sufficiently discriminative for evaluating model differences at the high end of performance.

***Stratified Assessment of Genomic Language Models Across Functional, Regulatory, and Variant-Level Tasks*** Building on our categorization of DNA-related tasks into three biologically meaningful groups, we further provide a structured analysis of gLMs’ performance across these categories: Genomic Function Annotation, Genetic Variant Effect Prediction, and Regulatory Mechanism Modeling. This taxonomy captures distinct levels of biological reasoning and modeling requirements. Genomic Function Annotation tasks emphasize recognition of known sequence features and yield relatively consistent performance across models. Regulatory Mechanism Modeling tasks, which require capturing dynamic, long-range regulatory interactions, result in the most pronounced divergence in model capabilities. Genetic Variant Effect Prediction tasks involve fine-grained changes and exhibit larger variation in model accuracy, reflecting differences in local sequence sensitivity. This stratified analysis offers a biologically grounded framework for evaluating the strengths and limitations of gLMs and highlights key areas for future methodological improvement.

**Performance Stratification of gLMs in Genomic Function Annotation Tasks.** In Genomic Function Annotation tasks, models generally exhibit stable and competitive performance across a broad range of architectures and training strategies. Tasks in this category typically rely on the identification of conserved sequence features or well-characterized regulatory motifs, which are often shared across species and supported by abundant experimental evidence. As a result, even relatively simple models such as CNN and LSTM can achieve performance comparable to that of more sophisticated pretrained gLMs.

Performance across six representative genomic annotation tasks reveals substantial variation in modeling difficulty and architectural sensitivity. Splice site prediction is the most tractable task, with several models—including NTv2 variants and DNABERT—achieving F1 scores above 0.910. SpliceBERT leads with 0.918, a result likely driven by its task-specific pretraining on splice junction–enriched sequences. In contrast, larger models such as MistralDNA underperform, suggesting that model size alone is insufficient for capturing localized signals. CpG methylation prediction shows similarly high and consistent performance across models (> 0.940), reflecting strong signal clarity and low task ambiguity.

Other tasks—such as promoter annotation, enhancer type classification, and regulatory element prediction—display more pronounced performance stratification, with F1 score gaps exceeding 0.1 between top and bottom models. These tasks, while foundational, remain non-trivial and offer effective resolution for assessing model quality. Epigenetic mark prediction further accentuates this divergence, yielding lower overall scores and greater architectural spread. Collectively, these findings emphasize that genomic function annotation tasks vary widely in complexity and discriminative power, and that robust, multi-task benchmarks are essential for evaluating model generalization in biologically meaningful settings.

In addition to architectural factors, we also observe a strong influence of input sequence length on classification performance. As shown in (Fig. 3b), increasing the input sequence length leads to a clear and consistent improvement in model performance. F1 scores steadily rise as the sequence length increases from 128 to over 8,000 nucleotides, with performance gains eventually plateauing near the upper input limits of each model. Model input length limits vary significantly due to architectural differences—Transformer-based models like DNABERT2 and NTv2-500m-Multi are capped at 8,192 tokens, whereas SSM-based models such as Hyena-Large and Caduceus-Ph support up to 32,768. Interestingly, although Hyena-Large underperforms below 8,192 tokens, its ability to handle longer inputs allows it to eventually outperform NTv2-500m-Multi at extended sequence lengths. The results underscore the importance of long-range genomic context in functional annotation and species-level inference tasks. Longer sequences contain richer evolutionary and structural signals, enabling models to capture broader patterns beyond local motifs. These findings emphasize that supporting long input contexts is critical for improving generalization in genomics, and should be a core consideration in future gLM design.

**Dissecting gLM Performance in Regulatory Mechanism Tasks.** Regulatory mechanism modeling tasks differ substantially from genomic function annotation and genetic variant effect prediction in terms of modeling requirements. These tasks emphasize understanding complex genomic regulatory relationships such as transcription factor binding, chromatin accessibility, and enhancer–promoter interactions. Unlike functional annotation tasks that often focus on well-characterized and conserved genomic elements, regulatory modeling involves context-dependent and dynamic regulatory components that vary across cell types, environmental conditions, and species. Moreover, rather than evaluating the effect of isolated variants, these tasks demand a more holistic understanding of regulatory networks and how different regions of the genome coordinate to influence gene expression.

Benchmark results reveal substantial variability in model performance across these tasks. For instance, in the gene expression prediction task, the best-performing model (Generator-1.2b) achieves an *R*^2^ of 0.548, while many others fall below 0.450. Enhancer activity and regulatory activity prediction tasks show similarly broad performance spreads, with Spearman scores ranging from 0.690 to 0.780, highlighting differences in models’ ability to resolve subtle regulatory variation. In contrast, the enhancer–promoter interaction task shows compressed F1 score distributions above 0.880, suggesting shared structural features that may be more uniformly captured across models.

Notably, model rankings vary substantially across tasks within this category. No single model demonstrates consistently superior performance, and task-specific advantages often depend on unique architectural choices or the nature of pretraining data. This inconsistency suggests that existing gLMs still face notable limitations in capturing the nuanced regulatory logic embedded within DNA sequences.

Taken together, regulatory mechanism modeling tasks provide a meaningful lens through which to evaluate a model’s ability to handle long-range dependencies, biological context sensitivity, and cross-regional coordination. These tasks highlight critical gaps in current model capabilities and represent an important direction for future improvements in genomic language modeling.

**Performance Profiling of gLMs in Variant Effect Prediction.**Genetic Variant Effect Prediction tasks are characterized by a requirement for high-resolution modeling, where models must distinguish subtle sequence changes, such as single-nucleotide variations, that can lead to significant functional consequences in gene regulation, transcriptional activity, or chromatin interactions. Unlike Genomic Function Annotation tasks, which often benefit from conserved sequence motifs and redundant evolutionary signals, these tasks typically involve loci with limited homologous support and high individual-specific variability. This lack of contextual redundancy increases the demand for precise, context-aware modeling of localized sequence patterns.

Despite these challenges, many gLMs exhibit strong capabilities on tasks that involve variant effect prediction, owing to their pretraining strategies based on masked language modeling, which encourage fine-grained sequence representation learning. Nevertheless, benchmark results reveal more diverse and spread-out performance profiles across models for this category. While some gLMs achieve consistently high scores, others demonstrate notable declines in performance, suggesting that variant-sensitive modeling imposes stronger differentiation among architectures and training regimes.

Some models, such as GPN, NT-2.5b-1000g, and Caduceus-Ps, achieve top-tier performance in specific tasks, but no single model consistently dominates across all benchmarks. For instance, GPN ranks first in genetic disease classification (F1 = 0.725), while NTv2 and Caduceus variants perform best on pathogenicity and splice effect prediction, respectively. The BRCA1/2 classification task yields the highest overall scores (up to F1 = 0.893), yet even here, the performance gap between top and bottom models remains substantial. This variability highlights the sensitivity of variant-level tasks to architectural differences, pretraining data, and sequence encoding strategies.

Taken together, these results suggest that tasks in this category serve as a sensitive probe for assessing a model’s ability to capture biologically meaningful variant effects. They provide valuable insights into how different gLMs handle localized sequence interpretation, long-range interactions, and the integration of weak regulatory signals, making them a critical component in evaluating the generalization and fine-tuning capabilities of genomic models.

### Genomic Language Models Show Robust Performance on RNA Tasks

To align with the DNA-based tokenizer, we replaced all *uracils (U)* with *thymines (T)* in RNA sequences before feeding them to the gLMs (Fig. 5a). gLMs pretrained on DNA data consistently surpass RNA-pretrained models across nearly all RNA benchmarks, highlighting the broad transferability of DNA-based contextual learning. As shown in Fig. 5b, DNA-pretrained models with similar parameter counts outperform RNA-pretrained models on RNA tasks.

**Figure 5.**
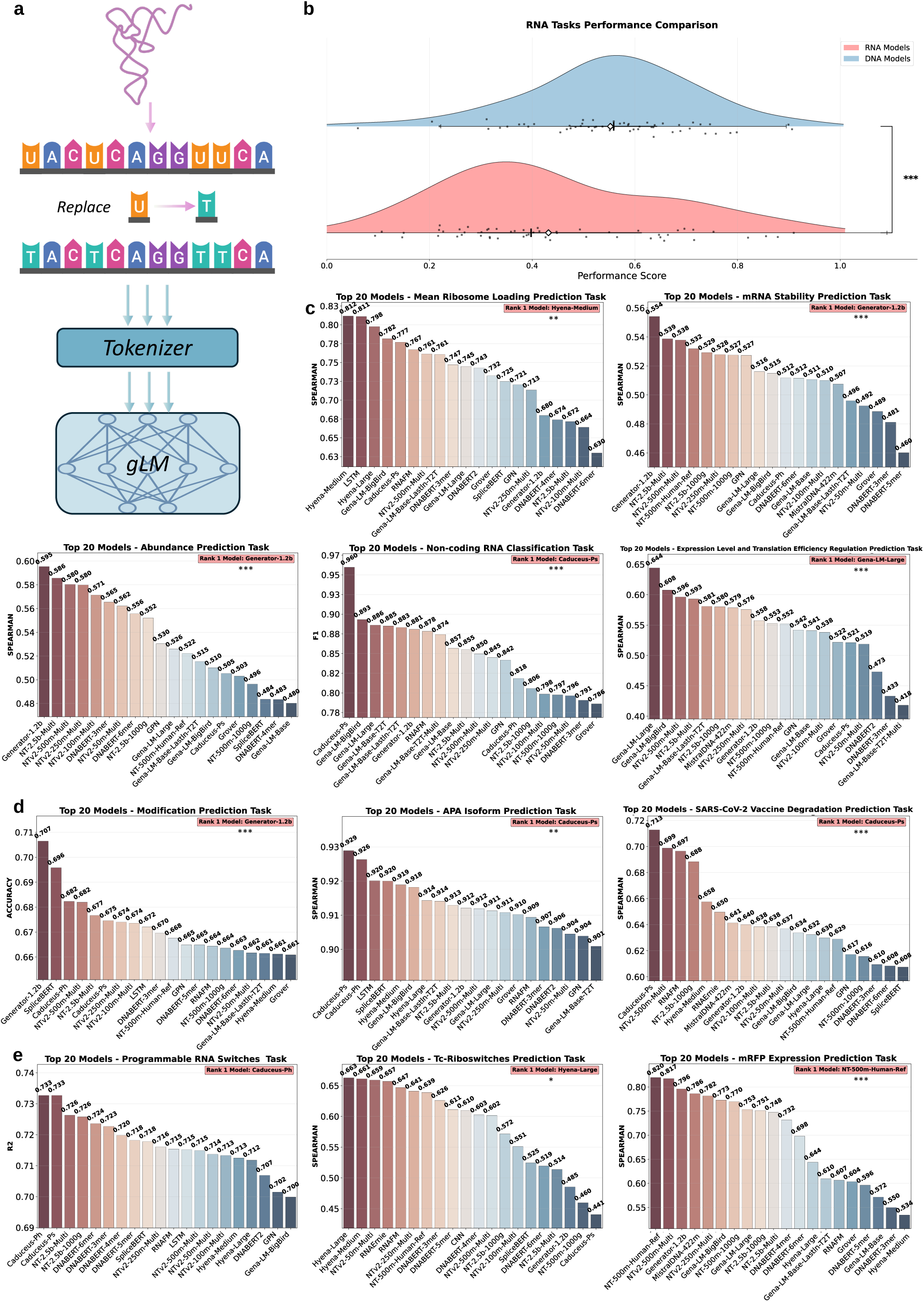
Model Performance across Diverse RNA-level Tasks. **a**, Schematic of RNA-to-DNA conversion for RNA-level genomic language modeling, where *uracil (U)* is replaced with *thymine (T)* prior to tokenization and model input. **b**, Performance comparison between RNA- and DNA-pretrained models on RNA tasks. DNA models (blue) outperform RNA models (red) with high statistical significance (***: *P <* 0.001). To ensure a fair comparison, we selected models with similar parameter counts: RNA models RNAErnie and RNAFM, and DNA models DNABERT2 and NTv2-100m-Multi—each with approximately 100 million parameters. **c**, Top-20 model performance on RNA functional studies tasks including mean ribosome loading prediction, mRNA stability prediction, abundance prediction, non-coding RNA classification, and translation regulation (***: *P <* 0.001, **: *P <* 0.01). **d**, Model performance on Post-transcriptional Regulation tasks including modification prediction, APA isoform prediction, and Sars-Cov-2 Vaccine degradation prediction (***: *P <* 0.001, **: *P <* 0.01). **e**, Model performance on RNA Engineering Applications tasks such as programmable switches prediction, Tc-riboswitches prediction, and mRFP expression prediction (***: *P <* 0.001, *: *P <* 0.05).

#### RNA Functional Studies

**Mean Ribosome Loading Prediction** involves predicting translation efficiency by estimating ribosome loading on a given mRNA sequence, with a particular focus on the 5’ UTR^43^. Ribosome loading, defined as the number of ribosomes actively translating an mRNA at any given time, reflects how efficiently the mRNA is converted into protein. Understanding how the 5’ UTR sequence influences ribosome loading is essential for optimizing protein expression, making this task highly relevant to biotechnology and therapeutic protein production. As shown in Fig. 5c, in the mean ribosome loading prediction task, Hyena-Medium (0.812) and LSTM (0.811) outperform all other models in terms of the Spearman score, followed closely by Hyena-Large (0.798), indicating that models with strong temporal modeling capabilities may better capture translational efficiency. Most gLMs achieve moderate performance in the 0.650–0.750 range, while models such as RNAErnie (0.265) and MistralDNA-1.6b (0.236) fall significantly behind (Table 18). The wide performance spread highlights the complexity of ribosome loading dynamics and suggests that specialized modeling of regulatory and structural features is necessary to improve predictive accuracy.

**Table 16.** F1 score of the evaluated models on Splice Variant Effect Prediction.

| Dataset | Exonic | Intronic | Avg |
| --- | --- | --- | --- |
| CNN | 0.573 ± 0.019 | 0.543 ± 0.019 | 0.558 ± 0.015 |
| LSTM | 0.767 ± 0.013 | 0.560 ± 0.020 | 0.663 ± 0.104 |
| DNABERT-3mer | 0.812 ± 0.011 | 0.521 ± 0.021 | 0.667 ± 0.146 |
| DNABERT-4mer | 0.784 ± 0.012 | 0.191 ± 0.020 | 0.488 ± 0.296 |
| DNABERT-5mer | 0.787 ± 0.012 | 0.487 ± 0.020 | 0.637 ± 0.150 |
| DNABERT-6mer | 0.785 ± 0.012 | 0.543 ± 0.019 | 0.664 ± 0.121 |
| DNABERT2 | 0.784 ± 0.012 | 0.403 ± 0.022 | 0.594 ± 0.190 |
| Gena-LM-Base | 0.772 ± 0.013 | 0.415 ± 0.023 | 0.594 ± 0.178 |
| Gena-LM-Base-T2T | 0.803 ± 0.012 | 0.619 ± 0.019 | 0.711 ± 0.092 |
| Gena-LM-Base-LastIn-T2T | 0.773 ± 0.013 | 0.607 ± 0.019 | 0.690 ± 0.083 |
| Gena-LM-Base-T2T-Multi | 0.809 ± 0.012 | 0.503 ± 0.022 | 0.656 ± 0.153 |
| Gena-LM-Large | 0.782 ± 0.012 | 0.673 ± 0.018 | 0.728 ± 0.054 |
| Gena-LM-BigBird | 0.784 ± 0.012 | 0.515 ± 0.021 | 0.650 ± 0.134 |
| Generator-1.2b | 0.626 ± 0.019 | 0.625 ± 0.020 | 0.625 ± 0.001 |
| GPN | 0.784 ± 0.012 | 0.000 ± 0.000 | 0.392 ± 0.392 |
| Grover | 0.652 ± 0.018 | 0.556 ± 0.020 | 0.604 ± 0.048 |
| MistralDNA-17m | 0.687 ± 0.015 | 0.475 ± 0.018 | 0.581 ± 0.106 |
| MistralDNA-422m | 0.649 ± 0.017 | 0.522 ± 0.018 | 0.585 ± 0.064 |
| MistralDNA-1.6b | 0.576 ± 0.018 | 0.426 ± 0.019 | 0.501 ± 0.075 |
| NT-500m-1000g | 0.655 ± 0.017 | 0.562 ± 0.019 | 0.609 ± 0.046 |
| NT-500m-Human-Ref | 0.660 ± 0.017 | 0.561 ± 0.019 | 0.611 ± 0.050 |
| NT-2.5b-1000g | 0.633 ± 0.018 | 0.591 ± 0.020 | 0.612 ± 0.021 |
| NT-2.5b-Multi | 0.683 ± 0.017 | 0.641 ± 0.019 | 0.662 ± 0.021 |
| NTv2-50m-Multi | 0.612 ± 0.018 | 0.612 ± 0.018 | 0.612 ± 0.000 |
| NTv2-100m-Multi | 0.592 ± 0.019 | 0.542 ± 0.020 | 0.567 ± 0.025 |
| NTv2-250m-Multi | 0.562 ± 0.020 | 0.578 ± 0.020 | 0.570 ± 0.008 |
| NTv2-500m-Multi | 0.620 ± 0.018 | 0.657 ± 0.019 | 0.638 ± 0.018 |
| Caduceus-Ph | 0.784 ± 0.011 | 0.000 ± 0.000 | 0.392 ± 0.392 |
| Caduceus-Ps | **0.835 ± 0.011** | **0.713 ± 0.016** | **0.774 ± 0.061** |
| Hyena-Large | 0.775 ± 0.013 | 0.501 ± 0.021 | 0.638 ± 0.137 |
| Hyena-Medium | 0.784 ± 0.012 | 0.387 ± 0.022 | 0.586 ± 0.198 |
| RNAErnie | 0.784 ± 0.012 | 0.000 ± 0.000 | 0.392 ± 0.392 |
| RNAFM | 0.784 ± 0.012 | 0.000 ± 0.000 | 0.392 ± 0.392 |
| SpliceBERT | 0.709 ± 0.016 | 0.615 ± 0.019 | 0.662 ± 0.047 |

**Table 17.**
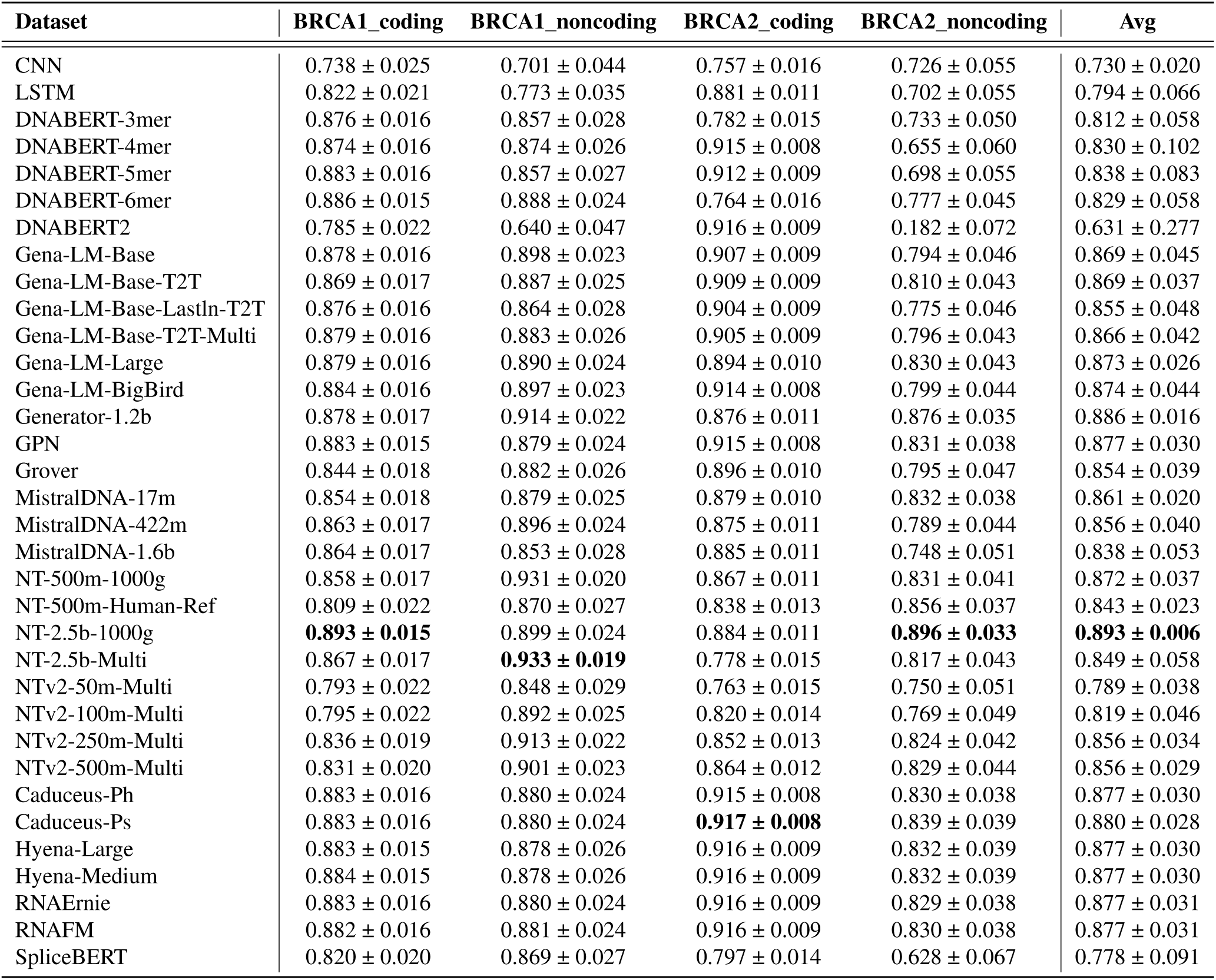
F1 score of the evaluated models on BRCA1/2 SNV Functional Impact Classification.

**Table 18.** Spearman score of the evaluated models on Mean Ribosome Loading Prediction.

| Dataset | MeanRibosomeLoading | MeanRibosomeLoading_Human | Avg |
| --- | --- | --- | --- |
| CNN | 0.718 ± 0.007 | 0.256 ± 0.036 | 0.487 ± 0.231 |
| LSTM | 0.818 ± 0.005 | 0.804 ± 0.019 | 0.811 ± 0.007 |
| DNABERT-3mer | 0.780 ± 0.006 | 0.714 ± 0.022 | 0.747 ± 0.033 |
| DNABERT-4mer | 0.777 ± 0.006 | 0.571 ± 0.027 | 0.674 ± 0.103 |
| DNABERT-5mer | 0.753 ± 0.006 | 0.474 ± 0.032 | 0.613 ± 0.139 |
| DNABERT-6mer | 0.753 ± 0.006 | 0.507 ± 0.032 | 0.630 ± 0.123 |
| DNABERT2 | 0.758 ± 0.006 | 0.728 ± 0.023 | 0.743 ± 0.015 |
| Gena-LM-Base | 0.711 ± 0.007 | 0.494 ± 0.030 | 0.603 ± 0.109 |
| Gena-LM-Base-T2T | 0.721 ± 0.007 | 0.473 ± 0.028 | 0.597 ± 0.124 |
| Gena-LM-Base-LastIn-T2T | 0.774 ± 0.006 | 0.748 ± 0.021 | 0.761 ± 0.013 |
| Gena-LM-Base-T2T-Multi | 0.726 ± 0.006 | 0.466 ± 0.029 | 0.596 ± 0.130 |
| Gena-LM-Large | 0.765 ± 0.006 | 0.725 ± 0.022 | 0.745 ± 0.020 |
| Gena-LM-BigBird | 0.792 ± 0.006 | 0.771 ± 0.018 | 0.782 ± 0.010 |
| Generator-1.2b | 0.754 ± 0.006 | 0.606 ± 0.027 | 0.680 ± 0.074 |
| GPN | 0.804 ± 0.005 | 0.637 ± 0.025 | 0.721 ± 0.084 |
| Grover | 0.767 ± 0.006 | 0.698 ± 0.020 | 0.733 ± 0.035 |
| MistralDNA-17m | 0.489 ± 0.010 | 0.187 ± 0.035 | 0.338 ± 0.151 |
| MistralDNA-422m | 0.661 ± 0.007 | 0.456 ± 0.032 | 0.559 ± 0.103 |
| MistralDNA-1.6b | 0.287 ± 0.011 | 0.184 ± 0.036 | 0.236 ± 0.052 |
| NT-500m-1000g | 0.649 ± 0.008 | 0.487 ± 0.031 | 0.568 ± 0.081 |
| NT-500m-Human-Ref | 0.546 ± 0.009 | 0.313 ± 0.034 | 0.429 ± 0.117 |
| NT-2.5b-1000g | 0.688 ± 0.007 | 0.519 ± 0.029 | 0.604 ± 0.085 |
| NT-2.5b-Multi | 0.721 ± 0.007 | 0.622 ± 0.026 | 0.672 ± 0.049 |
| NTv2-50m-Multi | 0.746 ± 0.006 | 0.514 ± 0.029 | 0.630 ± 0.116 |
| NTv2-100m-Multi | 0.737 ± 0.006 | 0.591 ± 0.028 | 0.664 ± 0.073 |
| NTv2-250m-Multi | 0.752 ± 0.006 | 0.675 ± 0.024 | 0.714 ± 0.039 |
| NTv2-500m-Multi | 0.800 ± 0.005 | 0.723 ± 0.020 | 0.761 ± 0.039 |
| Caduceus-Ph | 0.564 ± 0.009 | 0.456 ± 0.033 | 0.510 ± 0.054 |
| Caduceus-Ps | **0.841 ± 0.005** | 0.713 ± 0.022 | 0.777 ± 0.064 |
| Hyena-Large | 0.809 ± 0.006 | 0.786 ± 0.019 | 0.798 ± 0.012 |
| Hyena-Medium | 0.816 ± 0.005 | **0.807 ± 0.017** | **0.812 ± 0.005** |
| RNAErnie | 0.275 ± 0.011 | 0.254 ± 0.034 | 0.265 ± 0.010 |
| RNAFM | 0.821 ± 0.005 | 0.714 ± 0.022 | 0.767 ± 0.053 |
| SpliceBERT | 0.784 ± 0.006 | 0.666 ± 0.025 | 0.725 ± 0.059 |

**mRNA Stability Prediction**. Messenger RNA (mRNA) stability substantially impacts steady-state gene expression levels in a cell, governing how long transcripts remain available for translation. Because mRNA stability is strongly influenced by codon composition in a translation-dependent manner (referred to as codon optimality), it shapes protein abundance and modulates cellular function^44,45^. Accurate prediction of mRNA stability can therefore shed light on fundamental gene regulatory networks and assist in the design of more efficient expression systems. As a result, mRNA stability prediction serves as an important task for gLMs, testing their ability to capture the nuanced relationship between sequence composition, codon usage, and transcript degradation. As shown in Fig. 5c, in the mRNA stability prediction task, Generator-1.2b achieves the highest Spearman score of 0.554, followed closely by NT-2.5b-Multi (0.539) and NTv2-500m-Multi (0.538), indicating moderate success in capturing sequence determinants of transcript stability. The majority of models perform within a narrow range of 0.500-0.530, suggesting broadly similar capabilities among well-trained gLMs. However, as indicated in Table 19, performance drops significantly for models such as Hyena-Large (0.045), LSTM (0.256), and RNAErnie (0.258), highlighting the challenge of this regression task and the need for more specialized modeling to capture post-transcriptional regulatory features.

**Table 19.** Spearman score of the evaluated models on mRNA Stability Prediction.

| Dataset | mRNA_Stability |
| --- | --- |
| CNN | 0.297 ± 0.011 |
| LSTM | 0.256 ± 0.011 |
| DNABERT-3mer | 0.481 ± 0.009 |
| DNABERT-4mer | 0.454 ± 0.010 |
| DNABERT-5mer | 0.460 ± 0.010 |
| DNABERT-6mer | 0.512 ± 0.009 |
| DNABERT2 | 0.454 ± 0.010 |
| Gena-LM-Base | 0.511 ± 0.009 |
| Gena-LM-Base-T2T | 0.451 ± 0.010 |
| Gena-LM-Base-LastIn-T2T | 0.496 ± 0.009 |
| Gena-LM-Base-T2T-Multi | 0.429 ± 0.010 |
| Gena-LM-Large | 0.516 ± 0.009 |
| Gena-LM-BigBird | 0.515 ± 0.009 |
| Generator-1.2b | **0.554 ± 0.009** |
| GPN | 0.527 ± 0.009 |
| Grover | 0.488 ± 0.009 |
| MistralDNA-17m | 0.332 ± 0.011 |
| MistralDNA-422m | 0.507 ± 0.009 |
| MistralDNA-1.6b | 0.327 ± 0.011 |
| NT-500m-1000g | 0.527 ± 0.009 |
| NT-500m-Human-Ref | 0.532 ± 0.009 |
| NT-2.5b-1000g | 0.529 ± 0.009 |
| NT-2.5b-Multi | 0.539 ± 0.009 |
| NTv2-50m-Multi | 0.492 ± 0.010 |
| NTv2-100m-Multi | 0.510 ± 0.009 |
| NTv2-250m-Multi | 0.528 ± 0.009 |
| NTv2-500m-Multi | 0.538 ± 0.009 |
| Caduceus-Ph | 0.512 ± 0.009 |
| Caduceus-Ps | 0.357 ± 0.011 |
| Hyena-Large | 0.045 ± 0.012 |
| Hyena-Medium | 0.286 ± 0.011 |
| RNAErnie | 0.258 ± 0.011 |
| RNAFM | 0.263 ± 0.011 |
| SpliceBERT | 0.406 ± 0.010 |

**Abundance Prediction**. Transcript abundance refers to the concentration of mRNA molecules generated from a specific gene within a cell, which reflects gene expression levels at the transcriptional stage^46^. In contrast, protein abundance indicates the quantity of the corresponding protein present in the cell, representing the ultimate outcome of gene expression at the translational stage^46,47^. We benchmark gLMs to explore whether they can be used to predict transcript and protein abundance as measured by transcriptomics and proteomics experiments. As shown in Fig. 5c, in the abundance prediction task, Generator-1.2b achieves the highest Spearman score of 0.595, followed by NT-2.5b-Multi (0.586) and NTv2-500m-Multi (0.580), indicating strong performance from large-scale pretraining. Most models score between 0.500 and 0.560, reflecting moderate predictive capability. However, performance drops substantially for models such as RNAErnie (0.328) and MistralDNA-1.6b (0.293) (Table 20), highlighting the challenge of accurately modeling transcript and protein abundance and the importance of pretraining objectives aligned with expression-level signals.

**Table 20.**
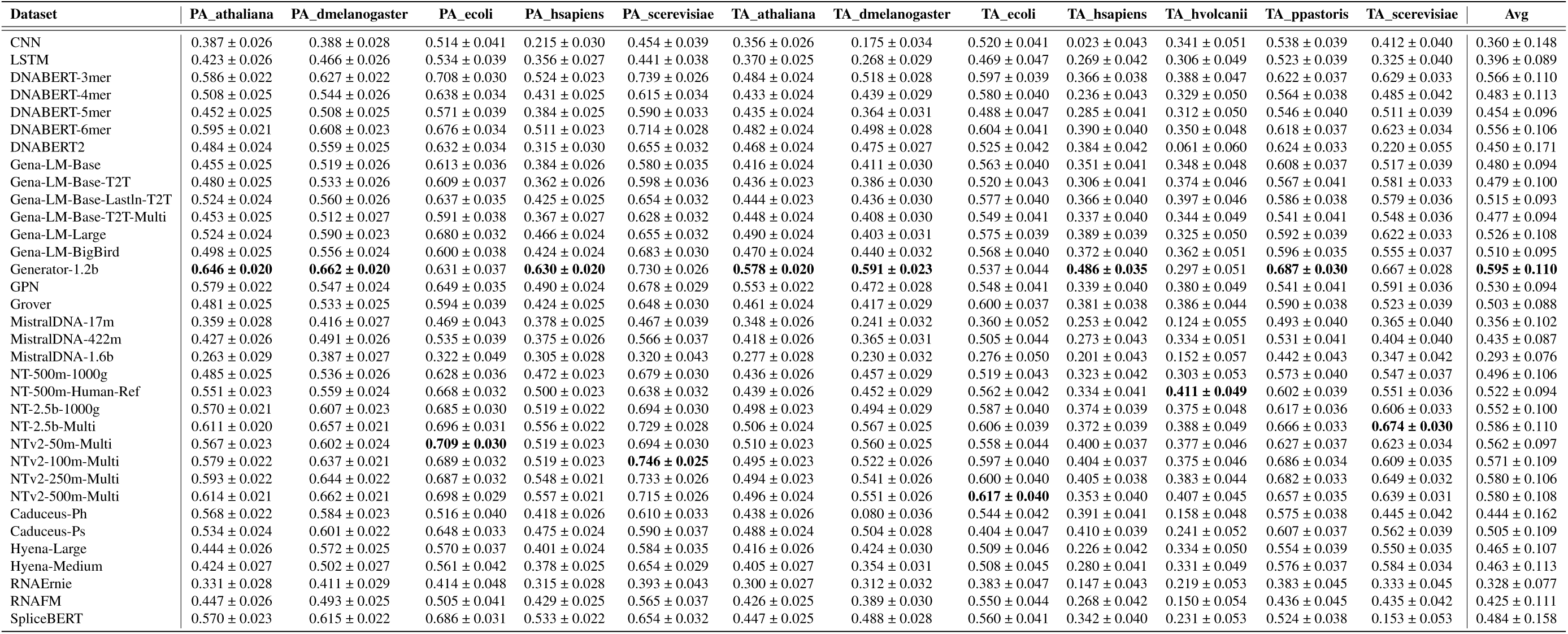
Spearman score of the evaluated models on Abundance Prediction.

**Non-coding RNA Classification** focuses on classifying non-coding RNA (ncRNA) sequences into functional categories. ncRNAs are RNA molecules that do not encode proteins but play critical roles in various cellular processes such as regulation of gene expression, RNA splicing, and chromatin remodeling^48,49^. gLMs can help uncover the regulatory roles of different ncRNA classes, such as miRNAs, lncRNAs, and siRNAs, in cellular functions. As shown in Fig. 5c, in the non-coding RNA classification task, Caduceus-Ps stands out with an F1 score of 0.960, significantly ahead of the next-best models, Gena-LM-BigBird (0.893) and Gena-LM-Large (0.886). Most models perform strongly, with over 20 models achieving F1 score above 0.800, indicating that this task is well captured by current gLMs. However, as illustrated in Table 21, a sharp decline is observed in lower-tier models such as MistralDNA-1.6b (0.235) and MistralDNA-17m (0.338).

**Table 21.** F1 score of the evaluated models on Non-coding RNA Classification.

| Dataset | NoncodingRNAFamily |
| --- | --- |
| CNN | 0.632 ± 0.009 |
| LSTM | 0.354 ± 0.008 |
| DNABERT-3mer | 0.791 ± 0.008 |
| DNABERT-4mer | 0.753 ± 0.008 |
| DNABERT-5mer | 0.714 ± 0.009 |
| DNABERT-6mer | 0.760 ± 0.008 |
| DNABERT2 | 0.786 ± 0.008 |
| Gena-LM-Base | 0.857 ± 0.007 |
| Gena-LM-Base-T2T | 0.885 ± 0.006 |
| Gena-LM-Base-LastIn-T2T | 0.883 ± 0.006 |
| Gena-LM-Base-T2T-Multi | 0.874 ± 0.006 |
| Gena-LM-Large | 0.886 ± 0.006 |
| Gena-LM-BigBird | 0.893 ± 0.005 |
| Generator-1.2b | 0.881 ± 0.006 |
| GPN | 0.842 ± 0.007 |
| Grover | 0.786 ± 0.008 |
| MistralDNA-17m | 0.338 ± 0.008 |
| MistralDNA-422m | 0.747 ± 0.008 |
| MistralDNA-1.6b | 0.235 ± 0.007 |
| NT-500m-1000g | 0.798 ± 0.008 |
| NT-500m-Human-Ref | 0.769 ± 0.008 |
| NT-2.5b-1000g | 0.806 ± 0.008 |
| NT-2.5b-Multi | 0.855 ± 0.007 |
| NTv2-50m-Multi | 0.796 ± 0.007 |
| NTv2-100m-Multi | 0.799 ± 0.008 |
| NTv2-250m-Multi | 0.845 ± 0.007 |
| NTv2-500m-Multi | 0.850 ± 0.007 |
| Caduceus-Ph | 0.818 ± 0.007 |
| **Caduceus-Ps** | **0.960 ± 0.004** |
| Hyena-Large | 0.598 ± 0.009 |
| Hyena-Medium | 0.744 ± 0.008 |
| RNAErnie | 0.753 ± 0.008 |
| RNAFM | 0.878 ± 0.006 |
| SpliceBERT | 0.675 ± 0.009 |

**Expression Level and Translation Efficiency Regulation Prediction**. We challenge gLMs with a task of profound biological importance: predicting both mRNA expression levels (EL) and translation efficiency (TE) from 5’ UTR sequences. This task, using datasets from diverse human tissues and cell lines^50^, directly assesses a model’s ability to bridge the gap between transcriptomic and proteomic outcomes by deciphering the complex regulatory interplay governing protein expression. As shown in Fig. 5c, in the expression level and translation efficiency regulation task, Gena-LM-Large achieves the top Spearman score of 0.644, indicating a strong capacity to capture rank-order relationships in gene expression and translation efficiency. Gena-LM-BigBird (0.608) and NTv2-500m-Multi (0.596) also perform competitively, while a gradual performance decline is observed across the majority of models. Lower-performing models, including LSTM (0.197) and CNN (0.175), highlight the difficulty of this task for architectures less capable of modeling complex regulatory signals (Table 22). The results underscore the importance of context-aware genomic modeling for capturing nuanced regulatory variation in transcriptomics.

**Table 22.** Spearman score of the evaluated models on Expression Level and Translation Efficiency Regulation Prediction.

| Dataset | HEK_EL | HEK_TE | Muscle_EL | Muscle_TE | pc3_EL | pc3_TE | Avg |
| --- | --- | --- | --- | --- | --- | --- | --- |
| CNN | 0.171 ± 0.027 | 0.233 ± 0.025 | 0.068 ± 0.094 | 0.077 ± 0.091 | 0.183 ± 0.029 | 0.317 ± 0.026 | 0.175 ± 0.087 |
| LSTM | 0.155 ± 0.027 | 0.230 ± 0.026 | 0.169 ± 0.083 | 0.171 ± 0.085 | 0.164 ± 0.028 | 0.293 ± 0.025 | 0.197 ± 0.049 |
| DNABERT-3mer | 0.420 ± 0.023 | 0.454 ± 0.022 | 0.378 ± 0.079 | 0.492 ± 0.075 | 0.364 ± 0.026 | 0.491 ± 0.023 | 0.433 ± 0.050 |
| DNABERT-4mer | 0.331 ± 0.025 | 0.421 ± 0.022 | 0.342 ± 0.080 | 0.378 ± 0.086 | 0.340 ± 0.027 | 0.456 ± 0.024 | 0.378 ± 0.046 |
| DNABERT-5mer | 0.316 ± 0.025 | 0.390 ± 0.025 | 0.401 ± 0.081 | 0.356 ± 0.080 | 0.354 ± 0.026 | 0.458 ± 0.025 | 0.379 ± 0.045 |
| DNABERT-6mer | 0.341 ± 0.024 | 0.433 ± 0.024 | 0.410 ± 0.081 | 0.343 ± 0.081 | 0.339 ± 0.027 | 0.459 ± 0.025 | 0.387 ± 0.049 |
| DNABERT2 | 0.537 ± 0.021 | 0.352 ± 0.025 | 0.337 ± 0.085 | 0.473 ± 0.074 | 0.552 ± 0.021 | 0.588 ± 0.021 | 0.473 ± 0.097 |
| Gena-LM-Base | 0.529 ± 0.021 | 0.536 ± 0.022 | 0.475 ± 0.081 | 0.574 ± 0.066 | 0.568 ± 0.021 | 0.565 ± 0.021 | 0.541 ± 0.034 |
| Gena-LM-Base-T2T | 0.450 ± 0.023 | 0.422 ± 0.023 | 0.234 ± 0.089 | 0.263 ± 0.092 | 0.392 ± 0.027 | 0.473 ± 0.023 | 0.372 ± 0.092 |
| Gena-LM-Base-LastIn-T2T | 0.565 ± 0.021 | 0.609 ± 0.021 | 0.504 ± 0.073 | 0.560 ± 0.071 | 0.587 ± 0.021 | 0.661 ± 0.019 | 0.581 ± 0.048 |
| Gena-LM-Base-T2T-Multi | 0.481 ± 0.021 | 0.461 ± 0.022 | 0.392 ± 0.081 | 0.300 ± 0.087 | 0.411 ± 0.026 | 0.464 ± 0.024 | 0.418 ± 0.062 |
| Gena-LM-Large | **0.673 ± 0.017** | **0.637 ± 0.020** | 0.576 ± 0.067 | 0.610 ± 0.067 | **0.676 ± 0.019** | **0.693 ± 0.018** | **0.644 ± 0.041** |
| Gena-LM-BigBird | 0.620 ± 0.020 | 0.615 ± 0.021 | 0.550 ± 0.072 | 0.647 ± 0.062 | 0.568 ± 0.022 | 0.647 ± 0.020 | 0.608 ± 0.037 |
| Generator-1.2b | 0.579 ± 0.020 | 0.556 ± 0.022 | 0.552 ± 0.071 | 0.542 ± 0.077 | 0.532 ± 0.024 | 0.584 ± 0.022 | 0.558 ± 0.019 |
| GPN | 0.596 ± 0.020 | 0.576 ± 0.021 | 0.367 ± 0.084 | 0.515 ± 0.068 | 0.578 ± 0.022 | 0.621 ± 0.020 | 0.542 ± 0.085 |
| Grover | 0.567 ± 0.021 | 0.554 ± 0.021 | 0.379 ± 0.086 | 0.503 ± 0.074 | 0.527 ± 0.023 | 0.600 ± 0.021 | 0.522 ± 0.071 |
| MistralDNA-17m | 0.218 ± 0.027 | 0.263 ± 0.025 | 0.269 ± 0.084 | 0.338 ± 0.084 | 0.222 ± 0.028 | 0.349 ± 0.027 | 0.276 ± 0.051 |
| MistralDNA-422m | 0.582 ± 0.021 | 0.548 ± 0.023 | 0.526 ± 0.078 | 0.653 ± 0.055 | 0.573 ± 0.023 | 0.589 ± 0.021 | 0.579 ± 0.039 |
| MistralDNA-1.6b | 0.366 ± 0.025 | 0.300 ± 0.025 | 0.113 ± 0.085 | 0.087 ± 0.088 | 0.249 ± 0.028 | 0.333 ± 0.026 | 0.241 ± 0.106 |
| NT-500m-1000g | 0.567 ± 0.021 | 0.542 ± 0.022 | 0.517 ± 0.079 | 0.549 ± 0.076 | 0.542 ± 0.024 | 0.601 ± 0.022 | 0.553 ± 0.026 |
| NT-500m-Human-Ref | 0.529 ± 0.022 | 0.550 ± 0.022 | 0.530 ± 0.076 | 0.575 ± 0.076 | 0.527 ± 0.024 | 0.602 ± 0.022 | 0.552 ± 0.028 |
| NT-2.5b-1000g | 0.586 ± 0.020 | 0.572 ± 0.021 | 0.584 ± 0.063 | 0.567 ± 0.066 | 0.552 ± 0.023 | 0.621 ± 0.020 | 0.580 ± 0.021 |
| NT-2.5b-Multi | 0.588 ± 0.020 | 0.580 ± 0.021 | 0.568 ± 0.071 | 0.660 ± 0.057 | 0.541 ± 0.024 | 0.623 ± 0.020 | 0.593 ± 0.039 |
| NTv2-50m-Multi | 0.510 ± 0.023 | 0.521 ± 0.023 | 0.529 ± 0.074 | 0.515 ± 0.075 | 0.491 ± 0.024 | 0.546 ± 0.023 | 0.519 ± 0.017 |
| NTv2-100m-Multi | 0.571 ± 0.021 | 0.517 ± 0.022 | 0.498 ± 0.076 | 0.559 ± 0.075 | 0.499 ± 0.025 | 0.587 ± 0.021 | 0.538 ± 0.035 |
| NTv2-250m-Multi | 0.569 ± 0.021 | 0.582 ± 0.021 | 0.511 ± 0.078 | 0.628 ± 0.064 | 0.563 ± 0.022 | 0.602 ± 0.021 | 0.576 ± 0.037 |
| NTv2-500m-Multi | 0.592 ± 0.020 | 0.588 ± 0.021 | **0.597 ± 0.067** | 0.618 ± 0.063 | 0.557 ± 0.024 | 0.628 ± 0.020 | 0.596 ± 0.023 |
| Caduceus-Ph | 0.128 ± 0.027 | 0.560 ± 0.020 | 0.113 ± 0.088 | **0.727 ± 0.052** | 0.036 ± 0.028 | 0.673 ± 0.019 | 0.373 ± 0.286 |
| Caduceus-Ps | 0.520 ± 0.022 | 0.540 ± 0.020 | 0.338 ± 0.082 | 0.622 ± 0.064 | 0.457 ± 0.025 | 0.651 ± 0.020 | 0.521 ± 0.104 |
| Hyena-Large | 0.215 ± 0.025 | 0.310 ± 0.024 | 0.305 ± 0.076 | 0.424 ± 0.074 | 0.235 ± 0.028 | 0.423 ± 0.024 | 0.319 ± 0.082 |
| Hyena-Medium | 0.192 ± 0.026 | 0.332 ± 0.025 | 0.415 ± 0.076 | 0.467 ± 0.078 | 0.178 ± 0.029 | 0.345 ± 0.026 | 0.322 ± 0.106 |
| RNAErnie | 0.094 ± 0.026 | 0.217 ± 0.026 | 0.227 ± 0.083 | 0.318 ± 0.082 | 0.107 ± 0.028 | 0.220 ± 0.027 | 0.197 ± 0.077 |
| RNAFM | 0.317 ± 0.025 | 0.403 ± 0.024 | 0.366 ± 0.088 | 0.359 ± 0.080 | 0.301 ± 0.026 | 0.448 ± 0.024 | 0.366 ± 0.050 |
| SpliceBERT | 0.240 ± 0.026 | 0.355 ± 0.024 | 0.207 ± 0.094 | 0.410 ± 0.075 | 0.188 ± 0.026 | 0.417 ± 0.025 | 0.303 ± 0.094 |

#### Post-transcriptional Regulation

**Modification Prediction** is a task focused on predicting the types of RNA modifications that occur on a precursor mRNA (pre-mRNA) sequence^50^. RNA modifications, such as m6A, m1A, m5C, and others, are post-transcriptional chemical changes that occur after the RNA is transcribed but before it matures. These modifications increase the structural and functional diversity of RNA molecules and regulate all stages of RNA’s life cycle. Precise identification of RNA modification sites is therefore of crucial importance in understanding the functions and regulatory mechanisms of various RNAs. As shown in Fig. 5d, in the modification prediction task, Generator-1.2b and SpliceBERT achieve the highest accuracy (0.707 and 0.696, respectively), closely followed by Caduceus-Ph (0.682), NTv2-500m-Multi (0.682) and NT-2.5b-Multi (0.677), while many large-scale gLMs, including some MistralDNA variants, perform substantially worse, with accuracy dropping below 0.550. This indicates that current large models may underperform on fine-grained base-level classification tasks without tailored pretraining objectives or architectures.

**APA Isoform Prediction**. Alternative polyadenylation (APA) is a key contributor to the rich transcriptomic diversity observed in human cells. In the APA process, genes can generate multiple mRNA isoforms by using different proximal polyadenylation signals (PAS). These isoforms can have distinct functional roles, influencing mRNA stability, translation efficiency, and interactions with other regulatory molecules. gLMs can be used to predict the usage ratio of proximal PAS in the 3’ UTR of mRNA^51^. As shown in Fig. 5d, in the APA isoform prediction task, nearly all models demonstrate excellent predictive performance, with Spearman scores exceeding 0.740. The top model, Caduceus-Ps, achieves the highest Spearman score of 0.929, followed closely by Caduceus-Ph and LSTM at 0.926 and 0.920, respectively. The narrow performance range among top-performing models indicates that the task is relatively well-solved, with even lightweight models such as LSTM performing on par with larger gLMs. Only a few models, particularly MistralDNA variants and CNN (Table 24), exhibit noticeable performance drops, falling below the 0.85 threshold.

**Table 23.** Accuracy of the evaluated models on Modification Prediction.

| Dataset | Modification |
| --- | --- |
| CNN | 0.604±0.003 |
| LSTM | 0.672±0.003 |
| DNABERT-3mer | 0.670±0.003 |
| DNABERT-4mer | 0.659±0.003 |
| DNABERT-5mer | 0.665±0.003 |
| DNABERT-6mer | 0.663±0.003 |
| DNABERT2 | 0.636±0.003 |
| Gena-LM-Base | 0.622±0.003 |
| Gena-LM-Base-T2T | 0.653±0.003 |
| Gena-LM-Base-Lastln-T2T | 0.662±0.003 |
| Gena-LM-Base-T2T-Multi | 0.644±0.003 |
| Gena-LM-Large | 0.635±0.003 |
| Gena-LM-BigBird | 0.659±0.003 |
| Generator-1.2b | **0.707±0.003** |
| GPN | 0.665±0.003 |
| Grover | 0.661±0.003 |
| NT-500m-1000g | 0.664±0.003 |
| NT-500m-Human-Ref | 0.668±0.003 |
| MistralDNA-17m | 0.512±0.003 |
| MistralDNA-422m | 0.601±0.003 |
| MistralDNA-1.6b | 0.504±0.003 |
| NT-2.5b-1000g | 0.659±0.003 |
| NT-2.5b-Multi | 0.677±0.003 |
| NTv2-50m-Multi | 0.662±0.003 |
| NTv2-100m-Multi | 0.674±0.003 |
| NTv2-250m-Multi | 0.674±0.003 |
| NTv2-500m-Multi | 0.682±0.003 |
| Caduceus-Ph | 0.682±0.003 |
| Caduceus-Ps | 0.675±0.003 |
| Hyena-Large | 0.650±0.003 |
| Hyena-Medium | 0.661±0.003 |
| RNAErnie | 0.631±0.003 |
| RNAFM | 0.664±0.003 |
| SpliceBERT | 0.696±0.003 |

**Table 24.** Spearman score of the evaluated models on APA Isoform Prediction.

| Dataset | Isoform |
| --- | --- |
| CNN | $0.844 \pm 0.002$ |
| LSTM | $0.920 \pm 0.001$ |
| DNABERT-3mer | $0.907 \pm 0.002$ |
| DNABERT-4mer | $0.900 \pm 0.002$ |
| DNABERT-5mer | $0.894 \pm 0.002$ |
| DNABERT-6mer | $0.898 \pm 0.002$ |
| DNABERT2 | $0.906 \pm 0.002$ |
| Gena-LM-Base | $0.896 \pm 0.002$ |
| Gena-LM-Base-T2T | $0.901 \pm 0.002$ |
| Gena-LM-Base-LastIn-T2T | $0.914 \pm 0.002$ |
| Gena-LM-Base-T2T-Multi | $0.891 \pm 0.002$ |
| Gena-LM-Large | $0.911 \pm 0.002$ |
| Gena-LM-BigBird | $0.918 \pm 0.001$ |
| Generator-1.2b | $0.912 \pm 0.002$ |
| GPN | $0.904 \pm 0.001$ |
| Grover | $0.910 \pm 0.001$ |
| MistralDNA-17m | $0.819 \pm 0.003$ |
| MistralDNA-422m | $0.869 \pm 0.002$ |
| MistralDNA-1.6b | $0.745 \pm 0.003$ |
| NT-500m-1000g | $0.893 \pm 0.002$ |
| NT-500m-Human-Ref | $0.889 \pm 0.002$ |
| NT-2.5b-1000g | $0.899 \pm 0.002$ |
| NT-2.5b-Multi | $0.913 \pm 0.001$ |
| NTv2-50m-Multi | $0.904 \pm 0.002$ |
| NTv2-100m-Multi | $0.899 \pm 0.002$ |
| NTv2-250m-Multi | $0.911 \pm 0.001$ |
| NTv2-500m-Multi | $0.912 \pm 0.002$ |
| Caduceus-Ph | $0.926 \pm 0.001$ |
| Caduceus-Ps | **$0.929 \pm 0.001$** |
| Hyena-Large | $0.914 \pm 0.002$ |
| Hyena-Medium | $0.919 \pm 0.001$ |
| RNAErnie | $0.856 \pm 0.002$ |
| RNAFM | $0.909 \pm 0.002$ |
| SpliceBERT | $0.920 \pm 0.001$ |

**SARS-CoV-2 Vaccine Degradation Prediction** focuses on predicting the hydrolysis rates of mRNA sequences. The degradation profiles for the first 68 nucleotides of these RNAs are measured using In-line-seq, a high-throughput method designed to characterize RNA degradation through in-line hydrolysis^52^. This task is crucial for addressing mRNA vaccine instability and designing stabilized RNA therapeutics. RNA instability, particularly due to in-line hydrolysis (the spontaneous chemical cleavage of the phosphodiester backbone), presents significant challenges for the storage, distribution, and efficacy of mRNA vaccines. As shown in Fig. 5d, in the SARS-CoV-2 vaccine degradation prediction task, Caduceus-Ps achieves the best performance with a Spearman score of 0.713, followed closely by NTv2-500m-Multi (0.699) and RNAFM (0.697). Overall, several models maintain strong correlation scores above 0.650, indicating a reasonable ability to capture degradation-related sequence features. However, performance varies significantly across models, with the lowest score dropping to 0.340 for Gena-LM-Base-T2T-Multi (Table 25), suggesting limited generalization capacity on this RNA degradation prediction task.

**Table 25.**
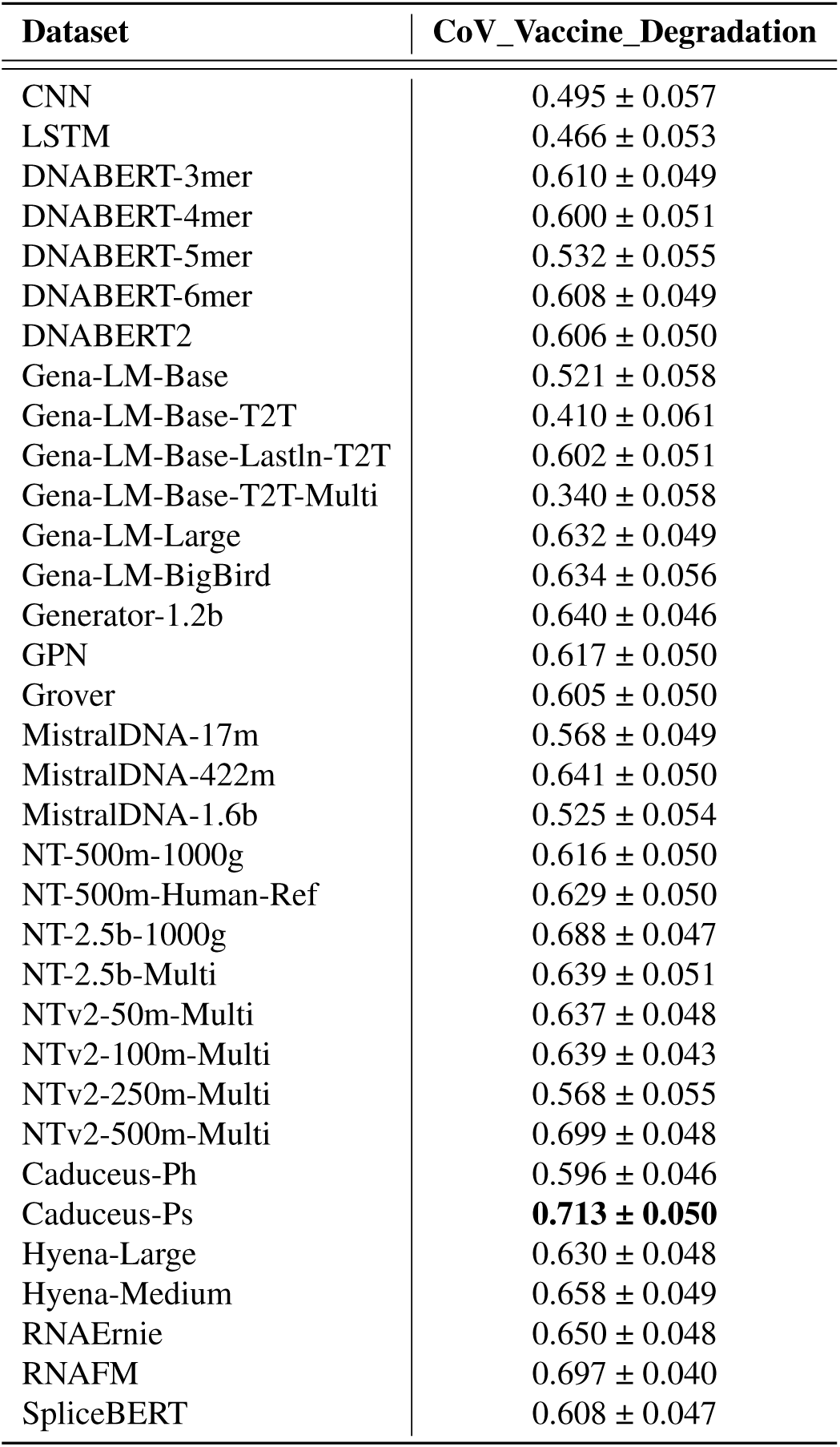
Spearman score of the evaluated models on SARS-CoV-2 Vaccine Degradation Prediction.

#### RNA Engineering Applications

**Programmable RNA Switches Prediction** represents a pivotal class of synthetic biology tools that enable precise detection and response to molecular signals (e.g., nucleic acids, proteins, or small molecules)^53^. These engineered RNA elements function as modular, sequence-specific biosensors, with applications ranging from genetic circuit design to diagnostic platforms. This task aims to predict the functional performance of programmable RNA switches by decoding the complex sequence-function relationships that govern their activity. As shown in Fig. 5e, for the programmable RNA switches prediction task, Caduceus-Ph and Caduceus-Ps achieve the highest *R*^2^ score of 0.733, followed closely by NT-2.5b-Multi (0.726) and NT-2.5b-1000g (0.726). A large number of models demonstrate highly similar performance clustered around the 0.700 mark, indicating consistent generalization in capturing RNA switch behavior. Nonetheless, several models underperform substantially, with MistralDNA-1.6b reaching only 0.258 (Table 26), revealing significant variance in model effectiveness on this structured RNA prediction task.

**Table 26.** ***R*^2^ score of the evaluated models on Programmable RNA Switches Prediction**

| Dataset | ProgrammableRNASwitches |
| --- | --- |
| CNN | $0.596 \pm 0.007$ |
| LSTM | $0.715 \pm 0.007$ |
| DNABERT-3mer | $0.723 \pm 0.007$ |
| DNABERT-4mer | $0.720 \pm 0.007$ |
| DNABERT-5mer | $0.718 \pm 0.007$ |
| DNABERT-6mer | $0.724 \pm 0.007$ |
| DNABERT2 | $0.707 \pm 0.007$ |
| Gena-LM-Base | $0.621 \pm 0.008$ |
| Gena-LM-Base-T2T | $0.668 \pm 0.008$ |
| Gena-LM-Base-Lastln-T2T | $0.690 \pm 0.007$ |
| Gena-LM-Base-T2T-Multi | $0.613 \pm 0.008$ |
| Gena-LM-Large | $0.699 \pm 0.007$ |
| Gena-LM-BigBird | $0.700 \pm 0.007$ |
| Generator-1.2b | $0.694 \pm 0.007$ |
| GPN | $0.702 \pm 0.007$ |
| Grover | $0.662 \pm 0.008$ |
| MistralDNA-17m | $0.372 \pm 0.009$ |
| MistralDNA-422m | $0.510 \pm 0.009$ |
| MistralDNA-1.6b | $0.258 \pm 0.009$ |
| NT-500m-1000g | $0.696 \pm 0.007$ |
| NT-500m-Human-Ref | $0.690 \pm 0.007$ |
| NT-2.5b-1000g | $0.726 \pm 0.007$ |
| NT-2.5b-Multi | $0.726 \pm 0.007$ |
| NTv2-50m-Multi | $0.714 \pm 0.006$ |
| NTv2-100m-Multi | $0.713 \pm 0.007$ |
| NTv2-250m-Multi | $0.716 \pm 0.007$ |
| NTv2-500m-Multi | $0.715 \pm 0.007$ |
| Caduceus-Ph | **$0.733 \pm 0.006$** |
| Caduceus-Ps | **$0.733 \pm 0.006$** |
| Hyena-Large | $0.712 \pm 0.007$ |
| Hyena-Medium | $0.713 \pm 0.007$ |
| RNAErnie | $0.684 \pm 0.008$ |
| RNAFM | $0.715 \pm 0.007$ |
| SpliceBERT | $0.718 \pm 0.007$ |

**Table 27.**
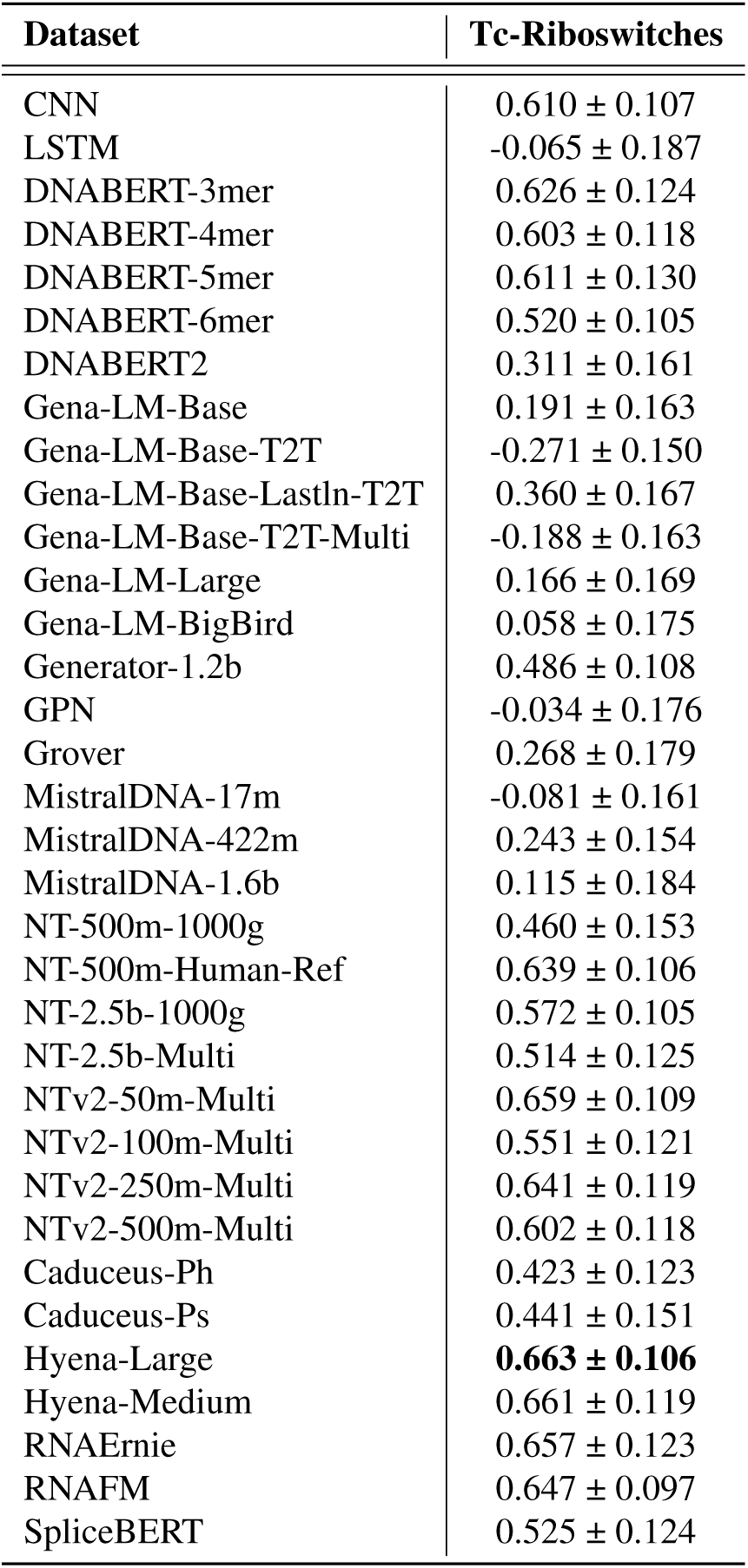
Spearman score of the evaluated models on Tc-Riboswitches Prediction.

**Tc-Riboswitches Prediction**. Riboswitches are RNA-based regulatory elements that control gene expression by undergoing conformational changes upon ligand binding, typically located in untranslated mRNA regions. The task challenges models to predict the regulatory performance of tetracycline (Tc) riboswitch variants, a capability crucial for the rational design of precisely tunable tools for synthetic biology^54^. As shown in Fig. 5e, on the Tc-Riboswitches prediction task, Hyena-Large achieves the highest Spearman score (0.663), marginally outperforming Hyena-Medium (0.661) and NTv2-50m-Multi (0.659). These top models exhibit superior capacity to capture riboswitch-mediated regulatory signals. In contrast, several models, including Gena-LM-Base-T2T and MistralDNA variants, perform poorly with negative correlations, suggesting systematic failure to generalize on this task. The wide spread of model performance highlights that successfully modeling synthetic, structured RNAs requires specific architectural or pretraining features that are not universally present in all models.

**mRFP Expression Prediction**. we challenge models to predict protein production levels from mRNA sequences that are identical in their amino acid translation but vary in their synonymous codon usage^55^. This task directly assesses a model’s ability to understand codon optimality, a key factor in tuning translation efficiency for synthetic biology and biotechnology applications. As shown in Fig. 5e, for the mRFP expression prediction task, NT-500m-Human-Ref achieves the highest Spearman score (0.820), followed closely by NTv2-500m-Multi (0.817) and Generator-1.2b (0.796), suggesting these models are particularly adept at capturing subtle sequence-expression relationships in synthetic regulatory elements. Conversely, as shown in Table 28, models such as SpliceBERT, CNN, and RNAErnie perform poorly, with RNAErnie even exhibiting a negative correlation (-0.169), indicating an inability to model the underlying biological signal effectively.

**Table 28.**
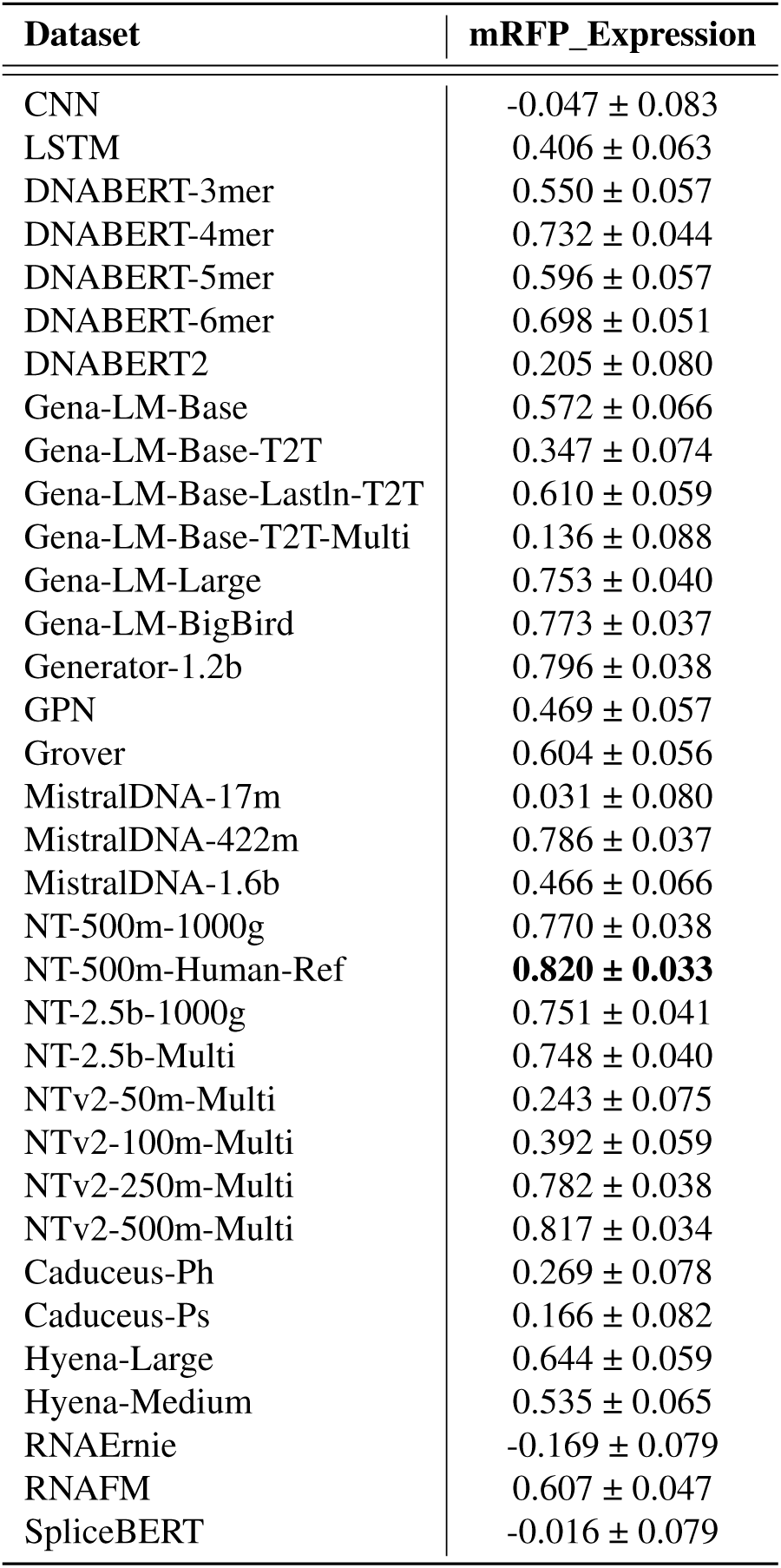
Spearman score of the evaluated models on mRFP Expression Prediction.

#### DNA-based gLMs Exhibit Superior Performance on RNA-level Tasks

The strong performance of DNA-pretrained gLMs on RNA benchmarks is a direct reflection of the central dogma’s first principle: transcription. Because RNA is transcribed directly from a DNA template, gLMs pretrained on genomic data are not just learning a similar sequence, but the very blueprint from which RNA is derived. This foundational knowledge is evident across different RNA-related tasks. RNA Functional Studies often require the ability to associate non-contiguous but functionally relevant sequence regions, which involves capturing long-range dependencies. This capability is well supported by the contextual learning mechanisms developed during DNA-based pretraining. Post-transcriptional Regulation tasks emphasize short-range sequence features, such as RNA-binding protein sites or splicing motifs near exon-intron boundaries. These features fall within the typical receptive field of models trained on DNA and are effectively handled by existing gLMs. RNA Engineering Applications place variable demands on the model, requiring both local motif detection and global sequence understanding depending on the task objectives.

Experimental results show that gLMs pretrained on DNA data outperform RNA-pretrained models across nearly all RNA tasks, as illustrated in Fig. 5. For example, the Generator-1.2b and Caduceus-Ps models rank first on several tasks and remains within the top three for most others, often with only small differences from the best-performing models. Even for models with a comparable number of parameters, RNA-pretrained models still underperform their DNA-pretrained counterparts (Fig. 5b). A key factor contributing to the relatively lower performance of RNA-pretrained models may be their reliance on character-level tokenization. While simple and biologically interpretable, this approach lacks the representational efficiency of k-mer or byte-pair encoding strategies commonly used in DNA models. These observations suggest that improvements in tokenization may enhance the performance of RNA-pretrained gLMs and contribute to more effective RNA modeling in future work.

### Genomic Language Models Excel in Protein-Level Predictions

We benchmarked gLMs on ten protein-centric tasks, using ESM-1b^5^ and ESM-2^56^ as protein-pretrained baselines to evaluate whether DNA pretraining captures protein-relevant patterns. For a comparison of pLMs with gLMs, we utilized curated CDS sequences as inputs for gLMs and their corresponding amino acid sequences for pLMs (Fig. 6a); the detailed procedure is described in the Materials and Methods section. As shown in Fig. 6b, we compared two 500M-parameter DNA models (NTv2-500m-Multi, NT-500m-1000g) with ESM-1b and ESM-2, which have comparable parameter counts. Despite being pretrained on genomic sequences, the gLMs outperform protein language models (pLMs) on most individual tasks and occupy the top ranks overall in the protein benchmark.

**Figure 6.**
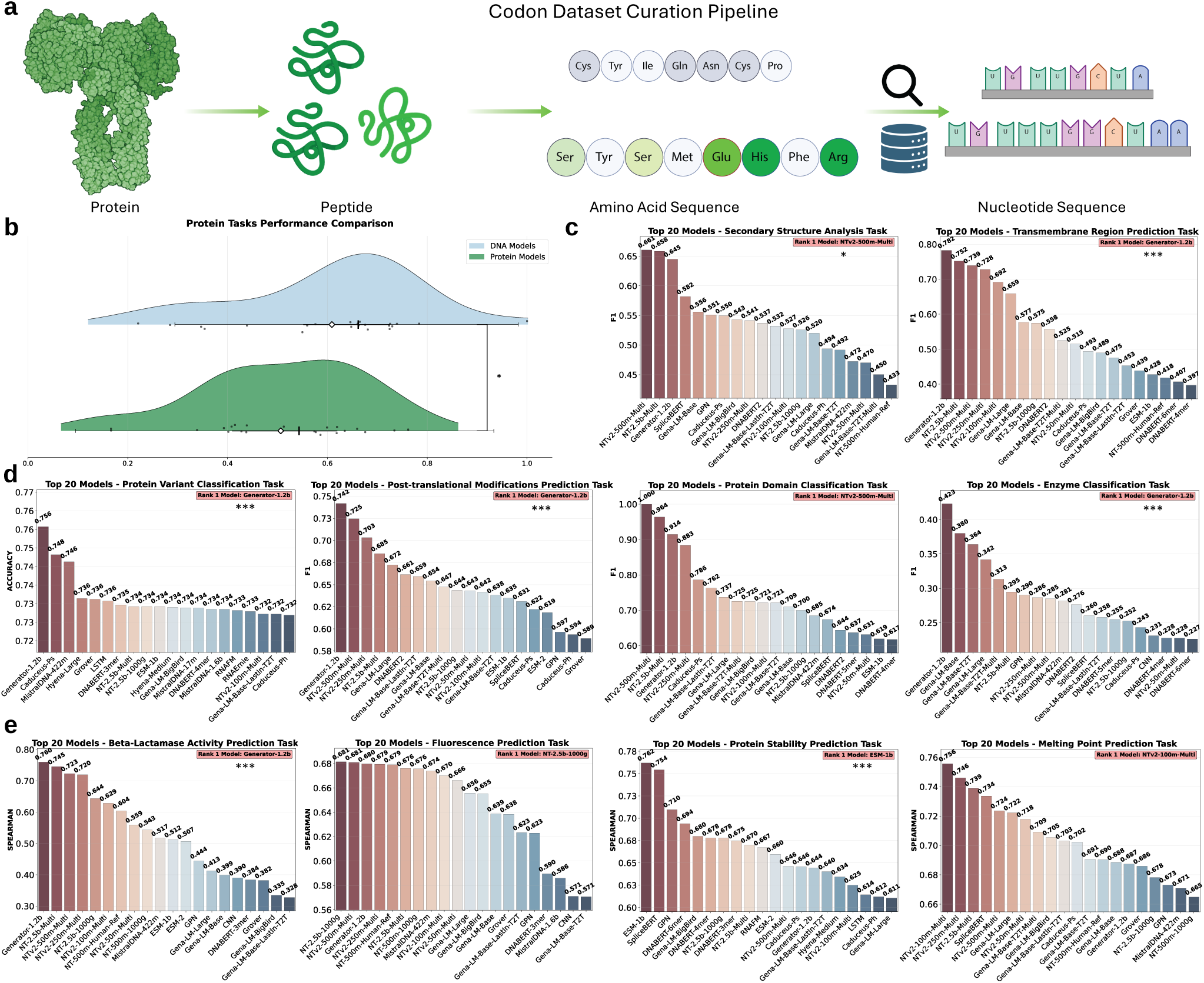
Model Performance on Curated Protein Level Benchmarks. **a**, Codon Dataset Curation Pipeline. Protein molecules are digested in silico to peptides, translated to amino-acid strings, back-translated to nucleotide (codon) sequences, and then filtered and annotated to produce task-ready datasets. **b**, Performance comparison between Protein- and DNA-pretrained models on Protein tasks. DNA models (blue) outperform Protein models (green) with high statistical significance (* *P <* 0.05). To ensure a fair comparison, we selected models with similar parameter budgets: NTv2-500m-Multi and NT-500m-1000G for the DNA models, and ESM-1b and ESM-2 for the protein models—all of which have approximately 500 million parameters. **c**, Top-20 model performance on Structural Analysis and Prediction tasks including secondary structure analysis and transmembrane region prediction (***: *P <* 0.001, *: *P <* 0.05). **d**, Top-20 Model performance on Protein Functional Annotation tasks including Protein variant classification, post-translational modifications prediction, protein domain classification and enzyme classification (***: *P <* 0.001). **e**, Top-20 Model performance on Protein Property Prediction tasks such as Beta-Lactamase activity prediction, fluorescence prediction, protein stability prediction and melting point prediction. (***: *P <* 0.001) gLMs to predict potential modification sites on a protein by analyzing the underlying genomic sequence that codes for it^60^. This capability is vital for understanding cellular regulation and developing novel therapeutics. As shown in Fig. 6d, in the post-translational modifications prediction task, Generator-1.2b achieves the best F1 score (0.742), followed by NTv2-500m-Multi (0.725) and NTv2-250m-Multi (0.703), indicating strong predictive power for modification sites. Other high-performing models include NT-2.5b-Multi (0.685) and Gena-LM-Large (0.672). These results highlight the effectiveness of large pretrained models in protein-level functional tasks.

#### Structural Analysis and Prediction

**Secondary Structure Analysis**. Understanding protein secondary structure is fundamental to elucidating protein function and folding mechanisms, as these local structural patterns dictate the overall three-dimensional conformation and biological activity of proteins. We employ the secondary structure analysis task to evaluate how effectively gLMs can predict and interpret local structural patterns in protein sequences. This task involves identifying structural elements such as *α*-helices, *β* -sheets, and random coils, which are fundamental units in protein folding processes^57^. Our evaluation shows that several gLMs can successfully perform this complex cross-modality prediction (Fig. 6c). NTv2-500m-Multi achieved the highest F1 score of 0.661, leading a top tier of models including NT-2.5b-Multi (0.658) and Generator-1.2b (0.645). In contrast, other architectures like Gena-LM-Large (0.520) and Caduceus-Ph (0.494) lagged behind (Table 29). This performance spread indicates that while the ability to infer protein structure from genomic context is achievable, it is highly dependent on the model’s capacity to learn the intricate rules of the genotype-to-structure relationship.

**Table 29.** F1 score of the evaluated models on Secondary Structure Analysis.

| Dataset | structure |
| --- | --- |
| CNN | 0.306 ± 0.023 |
| LSTM | 0.295 ± 0.017 |
| DNABERT-3mer | 0.306 ± 0.022 |
| DNABERT-4mer | 0.412 ± 0.024 |
| DNABERT-5mer | 0.409 ± 0.025 |
| DNABERT-6mer | 0.360 ± 0.023 |
| DNABERT2 | 0.537 ± 0.024 |
| Gena-LM-Base | 0.556 ± 0.024 |
| Gena-LM-Base-T2T | 0.492 ± 0.025 |
| Gena-LM-Base-LastIn-T2T | 0.532 ± 0.024 |
| Gena-LM-Base-T2T-Multi | 0.450 ± 0.024 |
| Gena-LM-Large | 0.520 ± 0.024 |
| Gena-LM-BigBird | 0.543 ± 0.025 |
| Generator-1.2b | 0.645 ± 0.023 |
| GPN | 0.551 ± 0.025 |
| Grover | 0.346 ± 0.022 |
| MistralDNA-17m | 0.274 ± 0.016 |
| MistralDNA-422m | 0.472 ± 0.025 |
| MistralDNA-1.6b | 0.258 ± 0.018 |
| NT-500m-1000g | 0.347 ± 0.024 |
| NT-500m-Human-Ref | 0.433 ± 0.024 |
| NT-2.5b-1000g | 0.526 ± 0.025 |
| NT-2.5b-Multi | 0.658 ± 0.024 |
| NTv2-50m-Multi | 0.470 ± 0.023 |
| NTv2-100m-Multi | 0.528 ± 0.024 |
| NTv2-250m-Multi | 0.541 ± 0.025 |
| NTv2-500m-Multi | **0.661 ± 0.022** |
| Caduceus-Ph | 0.494 ± 0.025 |
| Caduceus-Ps | 0.550 ± 0.026 |
| Hyena-Large | 0.404 ± 0.024 |
| Hyena-Medium | 0.392 ± 0.023 |
| RNAErnie | 0.272 ± 0.017 |
| RNAFM | 0.265 ± 0.017 |
| SpliceBERT | 0.582 ± 0.025 |
| ESM-1b | 0.339 ± 0.024 |
| ESM-2 | 0.402 ± 0.025 |

**Transmembrane Region Prediction**. Transmembrane proteins are essential components of cell membranes, functioning in cellular signal transduction and substance transport. Their misfolding or mutations may lead to various disorders, including cancer, diabetes, obesity, and neurodegenerative diseases^58^. To assess the abilities of gLMs to identify and locate transmembrane regions, we benchmarked their performance on carefully curated transmembrane region prediction datasets. This task effectively separated the models into distinct performance tiers (Fig. 6c). The top-performing gLMs, such as Generator-1.2b (F1 = 0.782) and NT-2.5b-Multi (0.752), proved highly capable of identifying the sequence features that define membrane-spanning domains. However, a significant performance gap was evident, with other architectures like the Gena-LM-Base variants failing to achieve F1 scores above 0.500 (Table 30). This wide disparity suggests that while many large models can learn general protein features, the ability to robustly identify specialized, biophysically constrained regions like transmembrane domains is a more challenging task that reveals key differences in architectural and pretraining effectiveness.

**Table 30.** F1 score of the evaluated models on Transmembrane Region Prediction.

| Dataset | transmembrane |
| --- | --- |
| CNN | 0.311 ± 0.022 |
| LSTM | 0.293 ± 0.016 |
| DNABERT-3mer | 0.345 ± 0.021 |
| DNABERT-4mer | 0.397 ± 0.022 |
| DNABERT-5mer | 0.385 ± 0.022 |
| DNABERT-6mer | 0.407 ± 0.023 |
| DNABERT2 | 0.558 ± 0.028 |
| Gena-LM-Base | 0.577 ± 0.030 |
| Gena-LM-Base-T2T | 0.476 ± 0.031 |
| Gena-LM-Base-LastIn-T2T | 0.453 ± 0.023 |
| Gena-LM-Base-T2T-Multi | 0.525 ± 0.033 |
| Gena-LM-Large | 0.659 ± 0.031 |
| Gena-LM-BigBird | 0.490 ± 0.019 |
| Generator-1.2b | **0.782 ± 0.029** |
| GPN | 0.332 ± 0.023 |
| Grover | 0.439 ± 0.020 |
| MistralDNA-17m | 0.150 ± 0.010 |
| MistralDNA-422m | 0.379 ± 0.024 |
| MistralDNA-1.6b | 0.213 ± 0.008 |
| NT-500m-1000g | 0.354 ± 0.022 |
| NT-500m-Human-Ref | 0.418 ± 0.024 |
| NT-2.5b-1000g | 0.575 ± 0.030 |
| NT-2.5b-Multi | 0.752 ± 0.029 |
| NTv2-50m-Multi | 0.515 ± 0.030 |
| NTv2-100m-Multi | 0.692 ± 0.030 |
| NTv2-250m-Multi | 0.728 ± 0.031 |
| NTv2-500m-Multi | 0.739 ± 0.030 |
| Caduceus-Ph | 0.254 ± 0.007 |
| Caduceus-Ps | 0.493 ± 0.026 |
| Hyena-Large | 0.379 ± 0.024 |
| Hyena-Medium | 0.375 ± 0.023 |
| RNAErnie | 0.177 ± 0.010 |
| RNAFM | 0.244 ± 0.016 |
| SpliceBERT | 0.342 ± 0.018 |
| ESM-1b | 0.428 ± 0.026 |
| ESM-2 | 0.393 ± 0.023 |

**Table 31.**
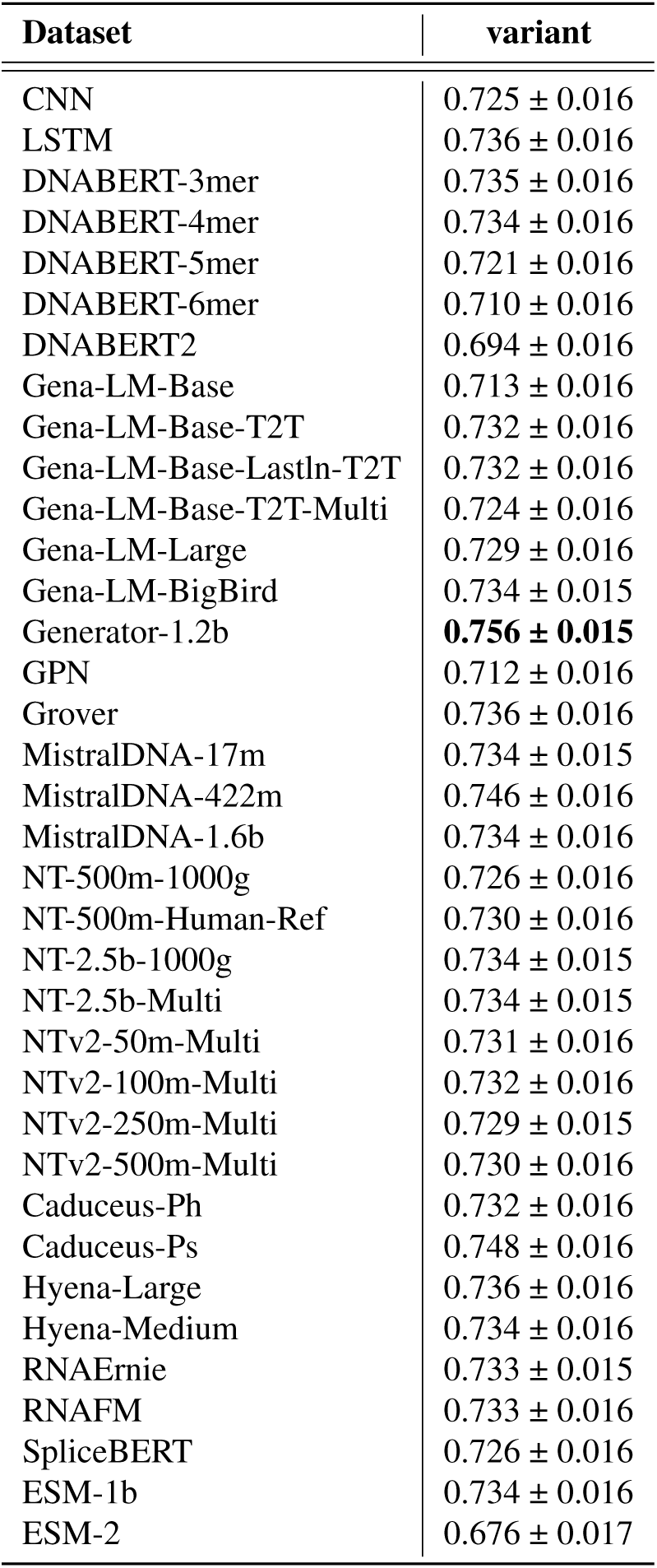
Accuracy of Protein Variant Classification.

**Table 32.** F1 score of the evaluated models on Post-translational Modifications Prediction.

| Dataset | ptm |
| --- | --- |
| CNN | 0.459 ± 0.017 |
| LSTM | 0.546 ± 0.017 |
| DNABERT-3mer | 0.567 ± 0.017 |
| DNABERT-4mer | 0.573 ± 0.017 |
| DNABERT-5mer | 0.563 ± 0.017 |
| DNABERT-6mer | 0.553 ± 0.017 |
| DNABERT2 | 0.662 ± 0.016 |
| Gena-LM-Base | 0.654 ± 0.017 |
| Gena-LM-Base-T2T | 0.638 ± 0.017 |
| Gena-LM-Base-LastIn-T2T | 0.659 ± 0.017 |
| Gena-LM-Base-T2T-Multi | 0.647 ± 0.017 |
| Gena-LM-Large | 0.672 ± 0.016 |
| Gena-LM-BigBird | 0.584 ± 0.017 |
| Generator-1.2b | **0.742 ± 0.016** |
| GPN | 0.597 ± 0.017 |
| Grover | 0.589 ± 0.017 |
| MistralDNA-17m | 0.453 ± 0.016 |
| MistralDNA-422m | 0.555 ± 0.017 |
| MistralDNA-1.6b | 0.490 ± 0.018 |
| NT-500m-1000g | 0.560 ± 0.018 |
| NT-500m-Human-Ref | 0.576 ± 0.016 |
| NT-2.5b-1000g | 0.644 ± 0.016 |
| NT-2.5b-Multi | 0.685 ± 0.015 |
| NTv2-50m-Multi | 0.643 ± 0.016 |
| NTv2-100m-Multi | 0.642 ± 0.016 |
| NTv2-250m-Multi | 0.703 ± 0.015 |
| NTv2-500m-Multi | 0.725 ± 0.015 |
| Caduceus-Ph | 0.594 ± 0.018 |
| Caduceus-Ps | 0.622 ± 0.018 |
| Hyena-Large | 0.562 ± 0.018 |
| Hyena-Medium | 0.563 ± 0.018 |
| RNAErnie | 0.404 ± 0.016 |
| RNAFM | 0.507 ± 0.018 |
| SpliceBERT | 0.631 ± 0.016 |
| ESM-1b | 0.635 ± 0.016 |
| ESM-2 | 0.619 ± 0.017 |

#### Protein Functional Annotation

**Protein Variant Classification**. Research has shown that protein variants can cause various hereditary disorders, including monogenic diseases and complex polygenic disorders^59^. Protein variant classification can help analyze and predict whether amino acid variations in protein sequences would lead to functional changes or disease phenotypes. Our benchmarking work demonstrates that gLMs are capable of capturing the subtle and intricate interactions among mutations, achieving predictive performance that approaches or even surpasses that of pLMs. As shown in Fig. 6d, in the protein variant classification task, Generator-1.2b outperforms all other models with an accuracy of 0.756, followed by Caduceus-Ps (0.748) and MistralDNA-422m (0.746). These models demonstrate strong capabilities in distinguishing functionally relevant protein variants. In contrast, pLMs such as ESM-1b (0.734) and ESM-2 (0.676) show relatively lower discriminative performance in this classification task. **Post-translational Modifications Prediction**. Post-translational modifications (PTMs) are critical for regulating protein function, influencing everything from signal transduction to cellular localization. In our PTM Prediction task, we challenge

**Protein Domain Classification** is critical for elucidating protein function by identifying and categorizing structurally and functionally autonomous regions within a protein^61^. These domains form stable three-dimensional structures through specific amino acid arrangements and frequently serve as the fundamental units for protein interactions and regulation. This task provides a robust benchmark for evaluating a gLM’s ability to recognize both local sequence motifs and long-range interactions that define domain boundaries and preserve structural integrity. As shown in Fig. 6d, for the protein domain classification task, NTv2-500m-Multi achieves perfect performance with an F1 score of 1.0, significantly outperforming all other models. It is followed by NT-2.5b-Multi (0.964) and Generator-1.2b (0.914), both of which also demonstrate strong classification capabilities. As summarized in Table 33, models like MistralDNA-17m (0.210) and MistralDNA-1.6b (0.343) perform considerably lower, indicating room for improvement in protein domain-level discrimination.

**Table 33.** F1 score of the evaluated models on Protein Domain Classification.

| Dataset | domain |
| --- | --- |
| CNN | $0.608 \pm 0.048$ |
| LSTM | $0.525 \pm 0.036$ |
| DNABERT-3mer | $0.597 \pm 0.041$ |
| DNABERT-4mer | $0.617 \pm 0.050$ |
| DNABERT-5mer | $0.637 \pm 0.047$ |
| DNABERT-6mer | $0.576 \pm 0.044$ |
| DNABERT2 | $0.644 \pm 0.053$ |
| Gena-LM-Base | $0.709 \pm 0.049$ |
| Gena-LM-Base-T2T | $0.721 \pm 0.049$ |
| Gena-LM-Base-LastIn-T2T | $0.763 \pm 0.046$ |
| Gena-LM-Base-T2T-Multi | $0.725 \pm 0.049$ |
| Gena-LM-Large | $0.737 \pm 0.050$ |
| Gena-LM-BigBird | $0.725 \pm 0.048$ |
| Generator-1.2b | $0.915 \pm 0.033$ |
| GPN | $0.248 \pm 0.031$ |
| Grover | $0.443 \pm 0.032$ |
| MistralDNA-17m | $0.210 \pm 0.025$ |
| MistralDNA-422m | $0.685 \pm 0.052$ |
| MistralDNA-1.6b | $0.343 \pm 0.032$ |
| NT-500m-1000g | $0.550 \pm 0.051$ |
| NT-500m-Human-Ref | $0.573 \pm 0.045$ |
| NT-2.5b-1000g | $0.700 \pm 0.047$ |
| NT-2.5b-Multi | $0.964 \pm 0.020$ |
| NTv2-50m-Multi | $0.632 \pm 0.042$ |
| NTv2-100m-Multi | $0.721 \pm 0.047$ |
| NTv2-250m-Multi | $0.883 \pm 0.036$ |
| NTv2-500m-Multi | **$1.000 \pm 0.000$** |
| Caduceus-Ph | $0.466 \pm 0.038$ |
| Caduceus-Ps | $0.786 \pm 0.049$ |
| Hyena-Large | $0.475 \pm 0.042$ |
| Hyena-Medium | $0.573 \pm 0.036$ |
| RNAErnie | $0.409 \pm 0.045$ |
| RNAFM | $0.363 \pm 0.038$ |
| SpliceBERT | $0.674 \pm 0.018$ |
| ESM-1b | $0.619 \pm 0.029$ |
| ESM-2 | $0.574 \pm 0.037$ |

**Table 34.** F1 score of the evaluated models on Enzyme Classification.

| Dataset | enzyme |
| --- | --- |
| CNN | 0.231 ± 0.019 |
| LSTM | 0.167 ± 0.007 |
| DNABERT-3mer | 0.171 ± 0.010 |
| DNABERT-4mer | 0.228 ± 0.018 |
| DNABERT-5mer | 0.255 ± 0.025 |
| DNABERT-6mer | 0.227 ± 0.024 |
| DNABERT2 | 0.276 ± 0.025 |
| Gena-LM-Base | 0.380 ± 0.040 |
| Gena-LM-Base-T2T | 0.364 ± 0.038 |
| Gena-LM-Base-LastIn-T2T | 0.258 ± 0.017 |
| Gena-LM-Base-T2T-Multi | 0.313 ± 0.033 |
| Gena-LM-Large | 0.342 ± 0.033 |
| Gena-LM-BigBird | 0.167 ± 0.008 |
| Generator-1.2b | **0.423 ± 0.040** |
| GPN | 0.290 ± 0.028 |
| Grover | 0.209 ± 0.023 |
| MistralDNA-17m | 0.166 ± 0.008 |
| MistralDNA-422m | 0.282 ± 0.018 |
| MistralDNA-1.6b | 0.167 ± 0.007 |
| NT-500m-1000g | 0.221 ± 0.017 |
| NT-500m-Human-Ref | 0.222 ± 0.017 |
| NT-2.5b-1000g | 0.252 ± 0.018 |
| NT-2.5b-Multi | 0.295 ± 0.021 |
| NTv2-50m-Multi | 0.228 ± 0.017 |
| NTv2-100m-Multi | 0.167 ± 0.007 |
| NTv2-250m-Multi | 0.287 ± 0.017 |
| NTv2-500m-Multi | 0.285 ± 0.017 |
| Caduceus-Ph | 0.167 ± 0.007 |
| Caduceus-Ps | 0.243 ± 0.017 |
| Hyena-Large | 0.167 ± 0.007 |
| Hyena-Medium | 0.167 ± 0.007 |
| RNAErnie | 0.167 ± 0.008 |
| RNAFM | 0.181 ± 0.013 |
| SpliceBERT | 0.260 ± 0.019 |
| ESM-1b | 0.167 ± 0.007 |
| ESM-2 | 0.166 ± 0.008 |

**Table 35.**
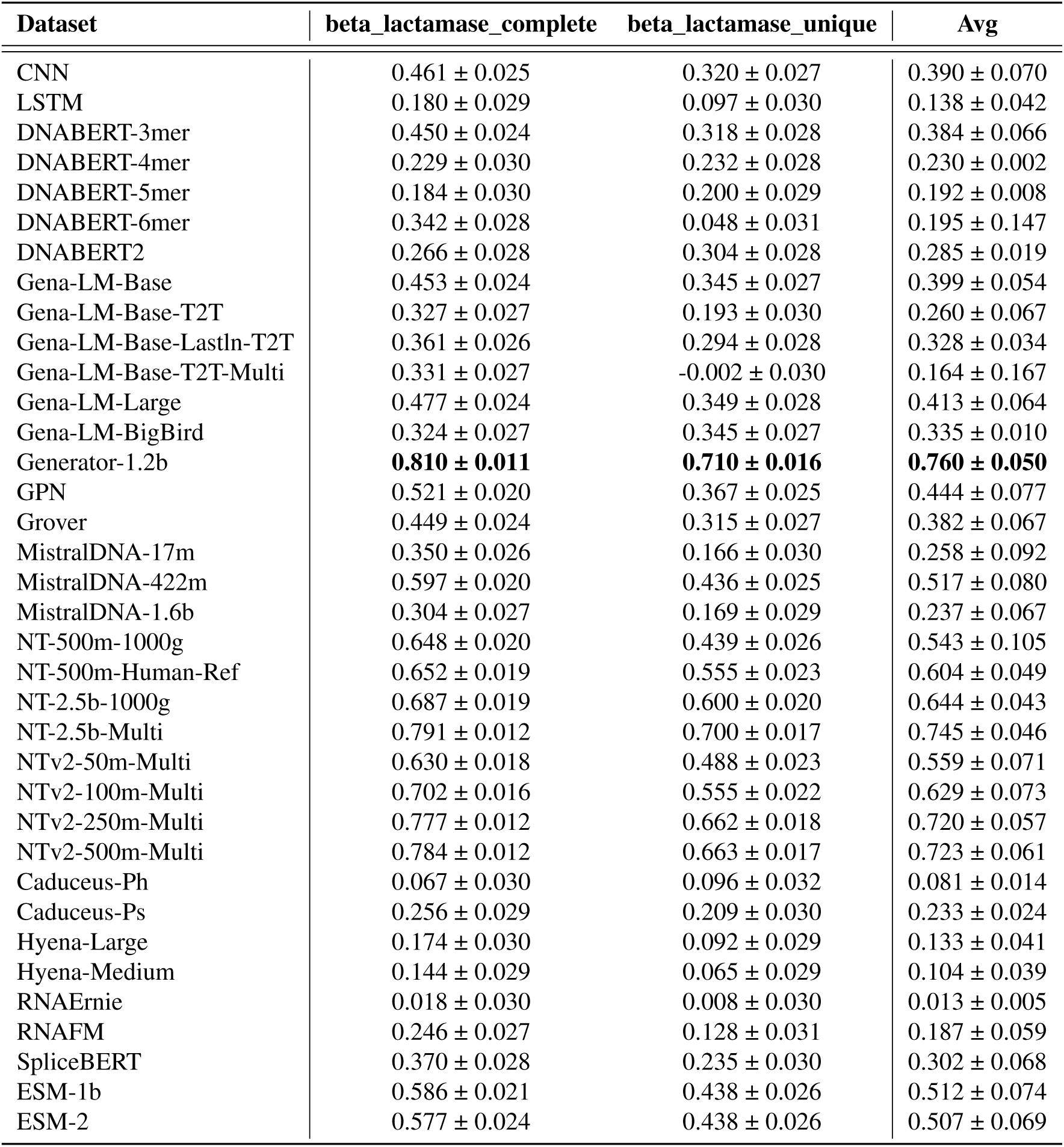
Spearman score of the evaluated models on Beta-Lactamase Activity Prediction.

**Table 36.** Spearman score of the evaluated models on Fluorescence Prediction.

| Dataset | fluorescence |
| --- | --- |
| CNN | $0.571 \pm 0.005$ |
| LSTM | $0.338 \pm 0.005$ |
| DNABERT-3mer | $0.590 \pm 0.004$ |
| DNABERT-4mer | $0.529 \pm 0.005$ |
| DNABERT-5mer | $0.540 \pm 0.005$ |
| DNABERT-6mer | $0.558 \pm 0.005$ |
| DNABERT2 | $0.548 \pm 0.004$ |
| Gena-LM-Base | $0.639 \pm 0.004$ |
| Gena-LM-Base-T2T | $0.571 \pm 0.005$ |
| Gena-LM-Base-LastIn-T2T | $0.623 \pm 0.004$ |
| Gena-LM-Base-T2T-Multi | $0.530 \pm 0.005$ |
| Gena-LM-Large | $0.656 \pm 0.004$ |
| Gena-LM-BigBird | $0.655 \pm 0.004$ |
| Generator-1.2b | $0.680 \pm 0.004$ |
| GPN | $0.623 \pm 0.004$ |
| Grover | $0.638 \pm 0.004$ |
| MistralDNA-17m | $0.468 \pm 0.005$ |
| MistralDNA-422m | $0.674 \pm 0.004$ |
| MistralDNA-1.6b | $0.586 \pm 0.005$ |
| NT-500m-1000g | $0.676 \pm 0.004$ |
| NT-500m-Human-Ref | $0.679 \pm 0.004$ |
| NT-2.5b-1000g | **$0.682 \pm 0.004$** |
| NT-2.5b-Multi | $0.676 \pm 0.004$ |
| NTv2-50m-Multi | $0.666 \pm 0.004$ |
| NTv2-100m-Multi | $0.670 \pm 0.004$ |
| NTv2-250m-Multi | $0.680 \pm 0.004$ |
| NTv2-500m-Multi | $0.681 \pm 0.004$ |
| Caduceus-Ph | $0.075 \pm 0.040$ |
| Caduceus-Ps | $0.075 \pm 0.040$ |
| Hyena-Large | $0.332 \pm 0.006$ |
| Hyena-Medium | $0.392 \pm 0.005$ |
| RNAErnie | $0.209 \pm 0.006$ |
| RNAFM | $0.341 \pm 0.005$ |
| SpliceBERT | $0.480 \pm 0.005$ |
| ESM-1b | $0.408 \pm 0.005$ |
| ESM-2 | $0.415 \pm 0.005$ |

**Table 37.** Spearman score of the evaluated models on Protein Stability Prediction.

| Dataset | stability |
| --- | --- |
| CNN | $0.467 \pm 0.006$ |
| LSTM | $0.614 \pm 0.006$ |
| DNABERT-3mer | $0.675 \pm 0.006$ |
| DNABERT-4mer | $0.678 \pm 0.006$ |
| DNABERT-5mer | $0.568 \pm 0.006$ |
| DNABERT-6mer | $0.694 \pm 0.005$ |
| DNABERT2 | $0.608 \pm 0.006$ |
| Gena-LM-Base | $0.458 \pm 0.007$ |
| Gena-LM-Base-T2T | $0.595 \pm 0.006$ |
| Gena-LM-Base-LastIn-T2T | $0.640 \pm 0.006$ |
| Gena-LM-Base-T2T-Multi | $0.557 \pm 0.006$ |
| Gena-LM-Large | $0.611 \pm 0.006$ |
| Gena-LM-BigBird | $0.680 \pm 0.005$ |
| Generator-1.2b | $0.644 \pm 0.005$ |
| GPN | $0.710 \pm 0.005$ |
| Grover | $0.463 \pm 0.007$ |
| MistralDNA-17m | $0.100 \pm 0.008$ |
| MistralDNA-422m | $0.332 \pm 0.008$ |
| MistralDNA-1.6b | $0.042 \pm 0.009$ |
| NT-500m-1000g | $0.576 \pm 0.007$ |
| NT-500m-Human-Ref | $0.538 \pm 0.007$ |
| NT-2.5b-1000g | $0.678 \pm 0.006$ |
| NT-2.5b-Multi | $0.670 \pm 0.006$ |
| NTv2-50m-Multi | $0.546 \pm 0.007$ |
| NTv2-100m-Multi | $0.625 \pm 0.006$ |
| NTv2-250m-Multi | $0.610 \pm 0.006$ |
| NTv2-500m-Multi | $0.646 \pm 0.006$ |
| Caduceus-Ph | $0.612 \pm 0.006$ |
| Caduceus-Ps | $0.646 \pm 0.006$ |
| Hyena-Large | $0.538 \pm 0.007$ |
| Hyena-Medium | $0.634 \pm 0.006$ |
| RNAErnie | $0.467 \pm 0.008$ |
| RNAFM | $0.667 \pm 0.005$ |
| SpliceBERT | $0.754 \pm 0.004$ |
| ESM-1b | **$0.762 \pm 0.004$** |
| ESM-2 | $0.660 \pm 0.005$ |

**Table 38.** Spearman score of the evaluated models on Melting Point Prediction.

| Dataset | melting_point |
| --- | --- |
| CNN | 0.454 ± 0.023 |
| LSTM | 0.535 ± 0.020 |
| DNABERT-3mer | 0.600 ± 0.018 |
| DNABERT-4mer | 0.444 ± 0.020 |
| DNABERT-5mer | 0.642 ± 0.016 |
| DNABERT-6mer | 0.664 ± 0.016 |
| DNABERT2 | 0.158 ± 0.022 |
| Gena-LM-Base | 0.688 ± 0.016 |
| Gena-LM-Base-T2T | 0.691 ± 0.016 |
| Gena-LM-Base-LastIn-T2T | 0.703 ± 0.015 |
| Gena-LM-Base-T2T-Multi | 0.709 ± 0.015 |
| Gena-LM-Large | 0.722 ± 0.014 |
| Gena-LM-BigBird | 0.705 ± 0.015 |
| Generator-1.2b | 0.687 ± 0.016 |
| GPN | 0.673 ± 0.017 |
| Grover | 0.686 ± 0.016 |
| MistralDNA-17m | 0.638 ± 0.018 |
| MistralDNA-422m | 0.671 ± 0.017 |
| MistralDNA-1.6b | 0.545 ± 0.021 |
| NT-500m-1000g | 0.665 ± 0.016 |
| NT-500m-Human-Ref | 0.690 ± 0.016 |
| NT-2.5b-1000g | 0.678 ± 0.016 |
| NT-2.5b-Multi | 0.739 ± 0.014 |
| NTv2-50m-Multi | 0.718 ± 0.015 |
| NTv2-100m-Multi | **0.756 ± 0.013** |
| NTv2-250m-Multi | 0.746 ± 0.014 |
| NTv2-500m-Multi | 0.724 ± 0.013 |
| Caduceus-Ph | 0.161 ± 0.024 |
| Caduceus-Ps | 0.702 ± 0.015 |
| Hyena-Large | 0.293 ± 0.023 |
| Hyena-Medium | 0.638 ± 0.018 |
| RNAErnie | 0.477 ± 0.021 |
| RNAFM | 0.477 ± 0.021 |
| SpliceBERT | 0.734 ± 0.015 |
| ESM-1b | 0.513 ± 0.022 |
| ESM-2 | 0.599 ± 0.019 |

**Enzyme Classification** plays a vital role in predicting the catalytic functions of enzymes, which are responsible for accelerating essential biochemical reactions in living organisms^62^. Grouping enzymes such as oxidoreductases, transferases, hydrolases, lyases, isomerases, and ligases contributes to a better understanding of metabolic pathways and supports drug development efforts. This classification task serves as an effective evaluation of a gLM’s capacity to detect subtle catalytic motifs and to distinguish fine-grained sequence variations that define different enzyme classes. As shown in Fig. 6d, for the enzyme classification task, Generator-1.2b leads with the highest F1 score (0.423), followed by Gena-LM-Base (0.380) and Gena-LM-Base-T2T (0.364). This significant performance gap suggests that the ability to capture specific enzymatic sequence features is not a general property of all gLMs and depend on specific architectural or pretraining designs.

#### Protein Property Prediction

**Beta-Lactamase Activity Prediction**. It is critical for gLMs to precisely predict the effects of single amino acid mutations^63^. In our benchmark evaluation, we focused on Beta-Lactamase activity as a regression task to explore the fitness landscape of all single codon substitutions in the TEM-1 gene, which is known to confer antibiotic resistance. The fitness of the gene was measured by dividing a library of mutants into thirteen sub-libraries, each exposed to increasing ampicillin concentrations and calculating fitness as a weighted average of allele counts normalized by wild-type fitness^64^. As shown in Fig. 6e, in the beta-lactamase activity prediction task, Generator-1.2b achieved the top Spearman score of 0.760, closely followed by NT-2.5b-Multi (0.745), NTv2-500m-Multi (0.723), and NTv2-250m-Multi (0.720). Other strong performers include NT-2.5b-1000g (0.644), NTv2-100m-Multi (0.629), and NT-500m-Human-Ref (0.604). These models demonstrate strong capabilities in modeling sequence-to-activity relationships, outperforming all other evaluated architectures by a clear margin.

**Fluorescence Prediction** assesses the gLMs’ capability to forecast the log-fluorescence values of higher-order mutant avGFP sequences. This regression task evaluates a model’s capacity to understand how complex combinations of mutations interact to determine protein function, providing critical insights for designing proteins with novel properties. As shown in Fig. 6e, for the fluorescence prediction task, NT-2.5b-1000g achieves the highest Spearman score (0.682), followed closely by NTv2-500m-Multi (0.681), Generator-1.2b (0.680), NTv2-250m-Multi (0.679) and NT-500m-Human-Ref (0.679). Several other models also demonstrate strong performance, including NT-2.5b-Multi (0.676) and NT-500m-1000g (0.676). These top models perform on par, suggesting a performance ceiling on this regression benchmark and highlighting the effectiveness of large-scale pretraining in capturing sequence-to-function fluorescence patterns.

**Protein Stability Prediction** is essential for effective protein engineering, as even minor alterations in stability can dramatically impact a protein’s functionality and overall performance. In particular, this regression task challenges models to accurately forecast the stability of variants located within a narrow, high-fitness region close to the wildtype state, a capability underscored by previous work^65^. Despite being trained on a broad range of sequences, models must exhibit the precision necessary to distinguish subtle differences among highly similar proteins. As shown in Fig. 6e, for the protein stability prediction task, ESM-1b achieves the highest Spearman score (0.762), followed by SpliceBERT (0.754) and GPN (0.710). Several other models also exhibit strong performance, including DNABERT-6mer (0.694), Gena-LM-BigBird (0.680) and DNABERT-4mer (0.678). These top-performing models show a clear advantage in capturing protein stability-related sequence patterns.

**Melting Point Prediction** is also critical for effective protein engineering, as even minor changes can significantly affect a protein’s function and overall performance. In particular, when evaluating the narrow, high-fitness regions proximal to the wild type, the task demands that gLMs accurately discern subtle differences in stability, a requirement that has been underscored in previous studies^66^. Although gLMs are trained on vast and diverse sequence datasets, they must still exhibit exceptional precision to differentiate minute stability variations among highly similar proteins. As shown in Fig. 6e, for the melting point prediction task, NTv2-100m-Multi ranks first with a Spearman score of 0.756, followed closely by NTv2-250m-Multi (0.746) and NT-2.5b-Multi (0.739). Several other models also achieve competitive performance, including SpliceBERT (0.734) and NTv2-500m-Multi (0.724). This consistent, strong performance among top-tier models suggests that large-scale pretraining is highly effective at capturing the sequence features that determine a protein’s thermal stability.

#### DNA-Pretrained gLMs Generalize Effectively to Protein Tasks

We evaluate the effectiveness of gLMs on protein-related tasks by conducting a comprehensive comparison against pLMs across ten benchmark tasks. For a controlled analysis, we focus on models of comparable size, such as NTv2-500m-Multi (500M parameters) and ESM-2 (650M parameters). NTv2 outperforms ESM-2 on seven out of ten tasks, including Transmembrane Region Prediction, Protein Domain Classification, Enzyme Classification, Fluorescence Prediction, Secondary Structure Analysis, Post-translational Modifications Prediction, and Melting Point Prediction. ESM-2 shows stronger results on three tasks: Protein Stability Prediction, Beta-Lactamase Activity Prediction, and Protein Variant Classification. These trends are consistent with observations reported in recent studies.

Beyond individual task comparisons, we observe that gLMs tend to achieve stronger overall performance across the protein benchmark. Among all evaluated models, the top five in average performance are exclusively gLMs, including Generator-1.2b and multiple models from the NTv2 family. In contrast, pLMs such as ESM-1b and ESM-2 occupy the middle tier. Traditional baselines like CNN and LSTM also rank lower on average. These results underscore the strong generalization ability of gLMs. Their effectiveness in modeling protein-level tasks stems from learning the foundational source code of biology; by training on genomic data, gLMs capture the underlying principles of the central dogma, learning the regulatory grammar and codon information that is ultimately translated into protein function but is lost to models that only see the final amino acid sequence.

### Benchmark Insights: Architecture, Scaling Laws, and Pretraining Data in Genomic Language Models

#### Dominance of Transformer Models in Genomics and the Potential of Efficient Alternatives

Transformer-based architectures continue to dominate the performance landscape of gLMs. As shown in Fig. 7c, 14 of the top 15 models by average benchmark performance are built upon Transformer designs. These models not only achieve state-of-the-art results across a wide range of tasks, but also exhibit strong robustness to hyperparameter configurations, allowing for streamlined fine-tuning across heterogeneous evaluation settings.

**Figure 7.**
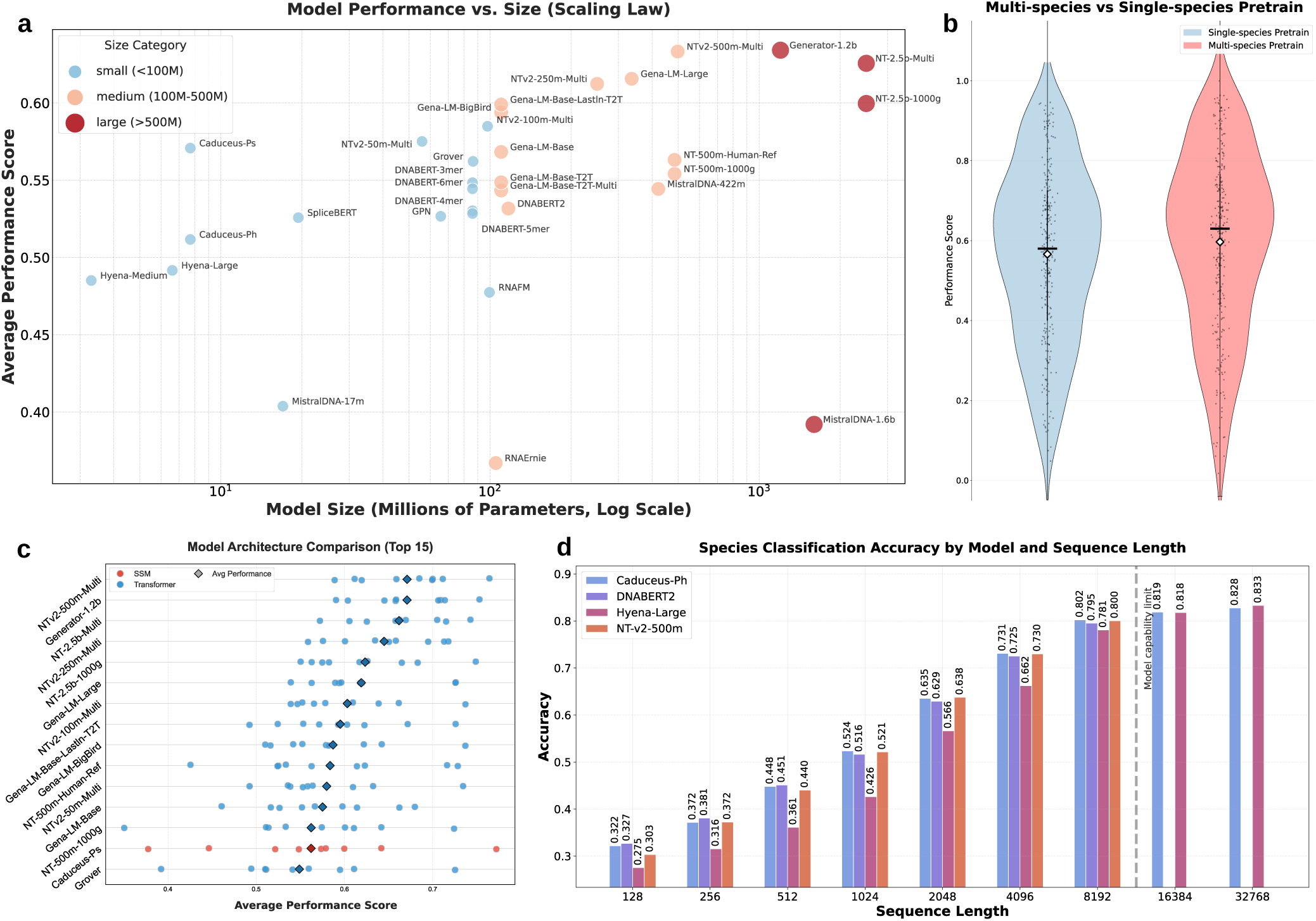
Benchmark Insights across Models and Tasks. **a**, Relationship between models’ parameter and average benchmark performance. Each point represents a pretrained model, with size on the x-axis (millions of parameters) and mean evaluation score on the y-axis. **b**, Violin plot comparing performance distributions of models pretrained on multi-species genomic data versus those pretrained only on human data. **c**, Comparison of models with different architectures. Transformer architecture remains the dominant paradigm for gLMs and SSM-based models are still promising. **d**, Bar chart of model-wise F1 scores at input lengths up to 32,768 base pairs. Models with extended context windows (e.g., Caduceus-Ph, Hyena-Large) exhibit the greatest gains as sequence length increases—particularly at lengths beyond 8,192 base pairs where other models could not be evaluated—underscoring the critical role of long-range context.

However, the attention mechanism central to Transformer architectures imposes quadratic computational complexity with respect to sequence length, resulting in practical constraints on input sizes. This limitation becomes particularly evident in tasks involving longer genomic contexts, as illustrated in Fig. 7d. While Transformer models (e.g., NTv2-500m) excel in the short-to-moderate input length regime, they fail to scale beyond 8,000 tokens due to the quadratic computational complexity of their self-attention mechanism. This highlights a fundamental architectural bottleneck for modeling ultra-long dependencies critical in certain genomics tasks.

To address this challenge, alternative architectures with linear-time complexity—particularly state space models (SSMs)—have been proposed. Notably, Caduceus^11^ adopts a Mamba-based design that leverages selective state updates to capture long-range patterns efficiently. Similarly, HyenaDNA^10^ is built on the Hyena framework, which combines convolutional operators with long-range filters to maintain scalability while preserving global context modeling. Despite their algorithmic efficiency, current SSM-based models often underperform Transformers in benchmark tasks and require greater sensitivity in hyperparameter tuning (Fig. 7c).

Nonetheless, these architectures remain highly promising. Their ability to handle extended sequence lengths with linear complexity offers a compelling path forward, especially as model scale and input demands grow. With further refinement in design and training regimes, SSM-based models may eventually rival—or even surpass—Transformer-based gLMs in both performance and generalization across diverse genomic applications. However, current results reveal a notable inconsistency. While SSM-based models perform well on certain tasks, such as those involving long-range dependencies or multi-species generalization, they often struggle on others, particularly tasks requiring precise token-level predictions. This lack of stability across benchmarks remains a significant limitation and underscores the need for further architectural and training improvements.

#### Insufficient Exploration of Scaling Laws in gLMs

A systematic investigation of scaling laws in gLMs is of critical importance. Scaling laws describe how model performance changes with respect to model size, training data scale, and computational budget.

While scaling laws have been extensively studied in natural language processing to guide model design and training strategies, their applicability to genomics remains less explored. Genomic sequences exhibit unique characteristics—such as high redundancy, strong local dependencies, and complex functional constraints—that may lead to fundamentally different scaling behavior in gLMs. Understanding how gLM performance evolves with model and data size is critical for addressing questions around model selection, generalization, and resource allocation. In particular, scaling analysis can inform trade-offs in low-data settings and guide the development of models that better align with the biological complexity of genomic tasks.

As shown in Fig. 7a, our benchmark experiments reveal that the scaling law for gLMs remains insufficiently explored. Among existing models, the NT series, including both NT and NTv2, presents the clearest scaling patterns. When pretrained on the 1000G dataset, NT models show improved classification performance score across tasks as parameter size increases from 500 million to 2.5 billion, with performance rising from 0.600 to 0.630. Furthermore, when the pretraining data for the 2.5 billion-parameter model is expanded to a multi-species dataset, its performance further increases to 0.645. A similar trend is observed in the NTv2 series, where models with 50 million, 100 million, 250 million, and 500 million parameters achieve progressively higher average performance scores of 0.599, 0.604, 0.639, and 0.660, respectively. However, when comparing NTv2 and NT directly, the 500 million-parameter NTv2 model outperforms the 2.5 billion-parameter NT model—demonstrating that superior architectural design can enable smaller models to surpass larger ones. At the same time, NTv2 has not been scaled to 2.5 billion parameters or beyond, highlighting the need for a more comprehensive exploration of scaling laws in gLMs. Meanwhile, in other model families, such consistent scaling behavior is not observed. We attribute this failure to suboptimal pretraining strategies—most notably the use of excessively short sequence lengths—which prevent models from fully leveraging their capacity and thus hinder meaningful exploration of scaling laws.

In summary, our results demonstrate that clear scaling trends emerge in the NT family—especially when model size and pretraining corpus diversity are jointly increased—but that these patterns break down both at extreme scales and in other architectures. This underscores that meaningful scaling laws for gLMs depend not only on parameter count but also on model design and pretraining protocol. Systematic, large-scale studies are therefore needed to map out these interdependencies and guide the development of truly scalable genomic language models.

#### Pretraining Data Composition and Strategy Matter in gLMs

Understanding how pretraining data influences gLM performance is essential for improving model accuracy, generalization, and biological relevance. One key observation is that training on multi-species genomic data significantly enhances performance. For example, the NT-2.5b model, which is pretrained on a phylogenetically diverse dataset spanning over 850 species, consistently outperforms its variant trained solely on human genomic data. This suggests that exposure to a broader range of evolutionary signals allows the model to better generalize across different genomic contexts and species.

Another important factor is the length of input sequences used during pretraining. Models trained on short fragments often struggle with tasks that require contextual understanding across longer genomic regions. This is evident in the case of MistralDNA, which was pretrained exclusively on short 200 base pair segments from the human genome (hg38). Despite its large parameter count, its performance lags behind that of smaller models trained on longer and more diverse sequences, indicating that insufficient sequence length during pretraining limits the model’s capacity to learn long-range dependencies.

Lastly, biologically informed pretraining strategies hold great promise but remain insufficiently explored. The Generator model demonstrates the potential of this approach by incorporating domain-specific biological priors. It is trained on 386 billion base pairs of eukaryotic DNA using a 6-mer tokenizer and employs objectives inspired by the central dogma of molecular biology. These include generating protein-coding sequences that produce valid amino acid chains and designing promoter regions that correspond to predefined gene expression levels. While such priors have led to notable performance gains, it remains unclear which types of biological knowledge consistently benefit model learning. Future work should further investigate and formalize biologically grounded pretraining strategies to enhance the interpretability and effectiveness of gLMs.

## Discussion

The central dogma of molecular biology describes the fundamental flow of genetic information from DNA to RNA and, ultimately, to protein. While gLMs are primarily trained on DNA sequences, their utility ultimately lies in enabling downstream biological insights across the full spectrum of the central dogma. It is therefore both natural and necessary to evaluate gLMs not only on DNA-centric tasks, but also on tasks involving RNA and protein sequences. To address this, we present a comprehensive benchmark—to our knowledge, the first of its kind—that systematically assesses the performance of gLMs across diverse tasks spanning the DNA, RNA, and protein modalities. Our benchmark emphasizes biological relevance, selecting or constructing tasks that reflect meaningful functional or structural aspects of molecular biology. Through this evaluation, we aim to clarify the strengths and limitations of current gLM architectures, training strategies, and generalization capabilities across the molecular continuum defined by the central dogma.

In our benchmark, we evaluate a diverse set of models, including classical baselines such as CNN and LSTM, as well as several representative families of gLMs. Across tasks involving DNA, RNA, and protein sequences, our results provide strong evidence that gLMs consistently outperform classical baselines, demonstrating robust and versatile modeling capabilities derived from their large-scale pretraining. DNA-related tasks are categorized into three groups: Genomic Function Annotation (Fig. 3), and Regulatory Mechanism Modeling (Fig. 4a) and Genetic Variant Effect Prediction (Fig. 4b). In Genomic Function Annotation tasks, gLMs achieve strong and stable performance, with even non-pretrained models such as CNN and LSTM performing competitively—reflecting the presence of consistent sequence signatures in functional regions. In contrast, Geneti Variant Effect Prediction tasks, which require the detection of subtle sequence alterations, exhibit broader performance dispersion across models. In these tasks, top-performing gLMs often show clear advantages. For example, GPN outperforms all other models with a notable F1 lead. Similarly, in the splice variant effect prediction task, Caduceus-Ps achieves the highest F1 score by a significant margin. Regulatory Mechanism Modeling, which involves long-range dependencies and complex interaction patterns, presents the greatest challenge, with no model demonstrating consistent superiority, highlighting current limitations in contextual representation learning. RNA tasks are grouped into RNA Functional Studies (Fig. 5c), Post-transcriptional Regulation (Fig. 5d), and RNA Engineering Applications (Fig. 5e). We find that DNA-pretrained gLMs transfer effectively to RNA modeling, outperforming RNA-pretrained models across all task categories. This advantage holds even when controlling for model size, as DNA-pretrained models consistently surpassed their RNA-pretrained counterparts with equivalent parameter counts (Fig. 5b). Large-scale models with diverse pretraining data, such as NT-2.5b-Multi, then build on this inherent advantage to achieve top or near-top performance. In contrast, RNA-pretrained models often underperform, likely due to character-level tokenization that fails to capture multi-nucleotide motifs and broader context. In comparison, k-mer-based gLMs trained on large genomic corpora exhibit stronger generalization across both local and global RNA features. Protein tasks span a range of Structural Analysis and Prediction (Fig. 6c), Functional Annotation (Fig. 6d), and Property Prediction (Fig. 6e). Despite being pretrained solely on nucleotide sequences, several gLMs achieve competitive or superior performance compared to protein-specific models. For example, NTv2-500m-Multi outperforms ESM-2 on seven out of ten benchmarks, particularly in tasks that depend on conserved sequence motifs such as enzyme classification and fluorescence prediction. Meanwhile, ESM-2 retains an advantage in tasks requiring codon-level precision or structural awareness, such as protein stability prediction. These results illustrate the cross-modal potential of gLMs and their capacity to capture biologically meaningful patterns along the central dogma’s axis of information flow.

Our comprehensive benchmark also highlights three key factors that influence the effectiveness of genomic language models: architectural design, model scaling, and pretraining configuration. **(1)** Transformer-based architectures remain the most consistently effective across a wide range of tasks. Under fixed input lengths, they outperform alternative designs in both accuracy and robustness, an advantage largely attributable to the powerful fine-grained modeling ability learned during large-scale pre-training and the intrinsic robustness of the self-attention mechanism. However, such fine-grained modeling ability often comes at the cost of computational efficiency. To address this limitation, recent developments in efficient sequence modeling, such as Mamba and Hyena, offer compelling advantages in scalability and computational efficiency. While these models do not yet surpass Transformers in benchmark performance, their linear-time operations and capacity for extended input contexts mark them as promising directions for future gLM development. Further refinements in their architectural design and training protocols may unlock their full potential. **(2)** The scaling behavior of gLMs remains poorly characterized. Our evaluation reveals that increases in model size and training data do not uniformly translate to better performance across all architectures. For instance, while NT and NTv2 models show some degree of improvement with larger scale, other families such as MistralDNA experience degraded performance as the parameter count increases. These inconsistencies suggest that scaling laws established in natural language processing may not apply directly in the genomic domain. More systematic studies are needed to disentangle the contributions of architecture, data composition and pretraining strategy. **(3)** Pretraining data composition and strategy critically shape gLM performance. Models trained on multi-species genomic datasets exhibit stronger generalization, particularly on tasks involving regulatory and evolutionary signals. Sequence length during pretraining also plays a decisive role, as models exposed only to short genomic segments during pretraining often underperform on tasks requiring long-range modeling. Additionally, biologically informed pretraining approaches offer a promising avenue. For example, the Generator model’s architecture directly embodies central dogma principles. It is explicitly optimized to maintain both coding and regulatory fidelity, an approach that results in superior downstream performance. These findings emphasize the importance of aligning data, biological relevance, and modeling objectives in the development of effective genomic language models.

Overall, the comprehensive benchmark we propose systematically covers key tasks across different biological molecule levels in the central dogma, including DNA, RNA, and proteins. It enables in-depth evaluation of gLMs from multiple dimensions, including modeling capacity and generalization ability. With unified data processing pipelines, task definitions, and performance metrics, this benchmark provides a reproducible and extensible evaluation framework for the gLM research community. This work contributes significantly to the development of more effective gene language models by guiding future improvements in architecture design, pretraining strategies, and cross-modal generalization. Meanwhile, we acknowledge several key limitations in the current scope of our benchmark, which highlight important directions for future work. First, our benchmark focuses on evaluating existing, publicly available gLMs. We believe our approach offers a pragmatic snapshot of the field’s current state. A key limitation, however, is that we did not conduct controlled experiments in which models were pretrained from scratch. Such experiments would enable systematic tests of foundational principles—such as architectural design, scaling laws, and pre-training strategies. Second, our evaluation focuses on discriminative tasks (e.g., classification and regression) and does not yet include generative tasks like protein or regulatory element design. This omission is primarily a reflection of the current gLM landscape, as the majority of evaluated models are not architected for sequence generation. Third, the benchmark is predominantly human-centric, with both tasks and pretraining datasets largely derived from the human genome. Consequently, the performance conclusions may not generalize across the broader tree of life, and extending the evaluation framework to include diverse eukaryotes, prokaryotes, and viruses is a critical next step. Finally, our benchmark relies on existing computational datasets and does not incorporate tasks that predict the outcomes of wet-lab experiments or involve a prospective experimental validation loop.

Building on our systematic evaluation, we also propose four possible directions to advance the future development of genomic language models. First, advancing the next generation of gLMs will require a holistic and principled approach. This begins with rethinking architectural paradigms to transcend the current trade-off between performance and efficiency, drawing inspiration from emerging models like Mamba and Hyena. Such novel architectures, in turn, necessitate a more principled framework for understanding scaling behavior, disentangling the complex interplay between model capacity and data heterogeneity. Ultimately, the full potential of these efficient and well-scaled architectures will only be unlocked through more biologically-informed pretraining strategies that incorporate evolutionary, regulatory, and structural priors, paving the way for gLMs that are not only powerful, but also interpretable and functionally grounded. Second, to address critical needs in experimental design, a key future direction is the development of conditional generative models. Such models are capable of designing novel DNA sequences tailored to specific protein-engineering objectives, such as optimizing enzymatic activity or tuning regulatory control. This capability could directly streamline essential wet-lab workflows—from mutagenesis library design to functional screening—and thereby empower biologists in their experimental planning. Third, a critical future direction is to extend the benchmark beyond the human genome to encompass a wider taxonomic range. This involves curating new datasets and tasks from diverse representative organisms, including other eukaryotes, prokaryotes, and viruses. Developing a multi-species evaluation framework would not only test the true generalization capabilities of current gLMs but also spur the development of models that can learn universal biological principles, rather than human-specific ones. Such efforts will be essential for applications in areas like microbial engineering, crop improvement, and emerging infectious disease surveillance. Fourth, to address the lack of experimental validation tasks in our benchmark and make gLMs more accessible to experimental biologists, we propose developing interactive AI agents powered by gLMs and LLMs. These agents could integrate with lab platforms to suggest protocols, design primers, interpret results, and provide real-time feedback, enabling closed-loop validation and bridging the gap between computation and experiment.

## Methods

### Benchmark Models

We aimed to conduct a comprehensive evaluation of publicly available genomic language models to fully understand their capabilities and limitations. To achieve this, we systematically surveyed the literature and compiled a representative set of models spanning various architectures, tokenization strategies, and pre-training data. Here, we present the detailed information on the models we employed, including their architecture design, pre-training data and tokenization strategies.

#### Baseline Model

**CNN**. Following^13^, we implement a three-layer 1D convolutional neural network (CNN) that first embeds input tokens into 100-dimensional vectors. Each convolutional block consists of Conv1d (kernel size 8), BatchNorm1d, ReLU, and MaxPool1d (pool size 2), with channel sizes 16→8→4. The resulting feature maps are flattened and projected through a 512-dimensional fully connected layer for the final classification and regression.

**LSTM**. We implement a two-layer bidirectional LSTM with 100-dimensional embeddings, 512 hidden units per direction, and a 0.15 dropout. The last-step hidden states are concatenated and passed through a 512-unit ReLU-activated dense layer before the final classification output.

#### Genomic Language Model

**DNABERT**^6^ is the first available gLM, a 12-layer BERT model with 86.1M parameters trained on sequences of length 512 from the human genome^67^. We not only used the checkpoint that tokenizes DNA as overlapping 6-mers, as the original DNABERT paper demonstrated superior fine-tuning performance with this configuration, but we also explored checkpoints that tokenize DNA using overlapping 3-mers, 4-mers, and 5-mers. This approach allows us to systematically investigate the impact of different k-mer sizes on model performance.

**GPN**^7^ is a transformer-inspired architecture that replaces traditional layers of self-attention with dilated CNN blocks. Specifically, the model comprises 25 stacked convolutional blocks that process fixed-length inputs of 512 base pairs, with each nucleotide encoded as a single-character token. GPN was pretrained with a masked language modeling objective on genomic sequences from Arabidopsis thaliana and seven additional Brassicales species (all sourced from NCBI Genome), enabling it to capture evolutionary patterns across these closely related plant genomes.

**Nucleotide Transformer**^8^ adopts an encoder-only architecture trained using the BERT-style masked language modelin objective. All variants of the model follow a unified architectural design, with differences arising from the diversity and scale of their pre-training data. Specifically, models were trained on three types of datasets: the human reference genome, the 1000 Genomes Project (3,202 diverse human genomes), and a cross-species dataset comprising 850 genomes. To accommodate long-range genomic contexts, DNA sequences are tokenized into non-overlapping 6-mers, enabling the model to process sequences of up to 5,994 base pairs in length.

**GENA-LM**^9^ consists of a series of medium-sized language models tailored for long-sequence modeling, built upon BERT and BigBird architectures. These models were pre-trained on both human and multi-species genomic datasets using the masked language modeling (MLM) objective. To further extend the receptive field, Byte-Pair Encoding (BPE) was employed for tokenization, allowing the model to handle input sequences of up to approximately 36,000 base pairs.

**GROVER**^68^ adopts a BERT-based architecture comprising 12 transformer encoder blocks. The model was pre-trained on the human reference genome using MLM. To encode the input sequences, GROVER applies BPE, enabling flexible token representation within sequences ranging from 20 to 510 tokens in length.

**DNABERT-2**^14^ is an enhanced version of DNABERT that expands pretraining beyond the human genome to include a large-scale multispecies genome dataset containing approximately 32.49B nucleotide bases, nearly 12 times the volume of the human genome dataset. Instead of traditional k-mer tokenization, DNABERT-2 adopts BPE to represent sequences. The model incorporates several architectural improvements to address limitations of the original DNABERT: replaces learned positional embeddings with Attention with Linear Biases (ALiBi)^69^ to remove input length constraints, integrates Flash Attention^70^ to increase computational efficiency, and revises the model structure to improve overall capacity.

**HyenaDNA**^10^ is a decoder-only language model based on the Hyena architecture, pretrained on the human reference genome with a next-nucleotide prediction objective at single-base resolution. The model consists of stacked blocks incorporating the Hyena operator, enabling it to handle input sequences up to 1 million base pairs in length, thus significantly enhancing its ability to model ultra-long DNA sequences.

**Nucleotide Transformer V2**^8^ is the second-generation model of Nucleotide Transformer, featuring several architectural improvements for enhanced efficiency. It replaces learnable positional embeddings with rotary embeddings applied at each attention layer and introduces bias-free gated linear units with Swish activation functions. NT-v2 accepts up to 2,048 input tokens, and with non-overlapping 6-mer tokenization, it can process sequences up to 12,282 base pairs in length, significantly extending the context window. The model is pretrained on a multispecies genome dataset comprising genomes from 850 species.

**Caduceus**^11^ is the first family of long-range DNA language models that simultaneously support bi-directionality and reverse complement equivariance, built on the efficient Mamba sequence modeling framework. Pretrained on the human reference genome with character-level tokenization, Caduceus can handle input sequences up to 131k base pairs in length. It integrates two key architectural components: the BiMamba module, which enables parameter-efficient bi-directional modeling, and the MambaDNA block, which enforces RC-equivariant behavior through shared computation over forward and reverse-complement sequences.

**Mistral-DNA** is a DNA-focused language model derived from the Mixtral-8×7B architecture and adapted for genomic sequence modeling. To better suit the task, the number of layers and hidden dimensions were reduced compared to the original Mixtral-8×7B. The model retains key architectural features such as Grouped-Query Attention, Sliding-Window Attention, and a Byte-fallback BPE tokenizer. Mistral-DNA was pre-trained on the human reference genome hg38 using input segments of 200 base pairs in length.

**Generator**^12^ is a generative genomic language model tailored for modeling long-context DNA sequences. Built on a transformer decoder architecture, it contains 1.2 billion parameters and supports input contexts up to 98,000 base pairs. The model is pre-trained on an extensive eukaryotic dataset comprising 386 billion bases, using the next-token prediction objective and a 6-mer tokenizer.

#### RNA Language Model

**RNAFM**^71^ is a BERT-based foundation model tailored for RNA-related studies. Trained with MLM on RNAcentral100^72^—a dataset containing 23.7 million ncRNA sequences—it processes inputs with a maximum length of 1024. RNAFM simply tokenizes 16 bases (“A”, “C”, “G”, “U”, “R”, “Y”, “K”, “M”, “S”, “W”, “B”, “D”, “H”, “V”, “N”, “-”) derived from the dataset as the model input. The model’s architecture consists of 12 transformer encoder blocks, with each encoder featuring a 640 hidden size feed-forward layer and a 20 multi-head self-attention layer.

**SpliceBERT**^73^ is based on the BERT architecture, consisting of six transformer encoder layers with a hidden size of 512 and 16 attention heads, totaling 19.4 million learnable parameters. The model encodes each nucleotide (A, G, C, T/U) as a discrete token and processes sequences up to 1024 tokens in length. Pretrained using MLM, SpliceBERT was trained on over 2 million pre-mRNA sequences (totaling 65 billion nucleotides) of 72 vertebrates obtained from the UCSC Genome Browser^74^.

**RNAErnie**^75^ is an RNA-focused pretrained model built upon the ERNIE^76^ architecture. The model consists of a 12-layer ERNIE transformer with a hidden dimension of 768, totaling approximately 105M trainable parameters. Pre-trained on 23 million ncRNA sequences from the RNAcentral^72^ database, it employs a basic tokenization scheme that encodes RNA bases and adds another functional token “[IND]” to denote RNA classes (*e.g.*, miRNA, mRNA, lnRNA) during pre-training.

#### Protein Language Model

**ESM-1b**^5^ is a 33-layer model with 650M parameters, built upon the BERT architecture with a 1280-dimensional embedding space. This model was pretrained on UR50/S^5^, a high-diversity sparse dataset comprising UniRef50^77^ representative sequences, where each 50% sequence identity cluster is represented by a single sequence.

**ESM-2**^56^ is similar to ESM-1b with a BERT architecture but introduces rotary position embeddings. Consistent with ESM-1b, we selected the 650M parameter version featuring 33 layers and 1280-dimensional embeddings. ESM-2 was pretrained on UR50/D^5^, a significantly larger high-diversity dense dataset that samples UniRef100 sequences evenly across UniRef50 clusters^77^, enabling training on over 60M protein sequences compared to ESM-1b’s 30M sequences.

### Benchmark Datasets

In this section, we detail each benchmark task and its corresponding dataset, with key statistics provided in Table 2.

#### Promoter Annotation

We downloaded human promoter data from Eukaryotic Promoter Database (EPDnew)^78^ and extracted two types of sequences centered around the transcription start site (TSS): 499 bp (from −399 to +100) for full promoter annotation, and 70 bp (from −34 to +35) for core promoter annotation. From the 499 bp sequences, we obtained 29,595 promoter regions, including 3,063 with a TATA-box, 9,785 with an Initiator, 4,803 with a CCAAT-box, and 13,365 with a GC-box. For the 70 bp sequences, only TATA-box and Initiator motifs were considered, as CCAAT-box and GC-box typically lie outside this region. Promoters containing both TATA-box and Initiator were excluded to ensure label exclusivity. Negative samples were generated by shuffling the nucleotide order of 50% of the promoter sequences, resulting in 14,797 non-promoter sequences. We then constructed two classification tasks: a six-class multi-label task based on the 499 bp sequences (four motif types, motif-free promoters, and non-promoters), and a four-class task based on the 70 bp sequences (TATA-box, Initiator, motif-free promoters, and non-promoters). All datasets were split into train, validation, and test sets in an 8:1:1 ratio.

#### Enhancer Types Classification

We followed the data processing procedure described in Nucleotide Transformer (NT)^8^, retrieving human enhancer elements from the ENCODE’s SCREEN database^79^, including both distal and proximal enhancers. Based on the vocabulary from the study^80^, the enhancers were categorized into tissue-specific and tissue-invariant enhancers. Enhancers overlapping regions classified as tissue-invariant were labeled as tissue-invariant, while all other enhancers were defined as tissue-specific. These data were created as a three-class classification task with labels including tissue-specific enhancer, tissue-invariant enhancer, and non-enhancer.

#### Splice Sites Prediction

We obtained all human annotated splice sites from GENCODE^81^ V47 gene annotation. Annotations were filtered to exclude level-3 transcripts, ensuring that all training data were manually curated, while considering only data from chromosomes 1-22 and the X and Y chromosomes. We selected 512 bp-long sequences containing a splice acceptor or donor site in the center as positive examples, with a total of 50,000 positive examples. As negative examples, we selected 50,000 sequences of 512 bp in length that do not overlap with any splice sites. Using the retrieved sequences, we constructed three supervised learning tasks to evaluate model performance on splice site prediction. The first is a multi-label classification task, termed Splice site all, where each sequence is labeled as one or more of the following: acceptor, donor, or none. In addition, we formulated two binary classification tasks: Splice acceptor, which distinguishes acceptor splice sites from non-acceptor sites, and Splice donor, which distinguishes donor splice sites from non-donor sites. These tasks allow for a comprehensive evaluation of the model’s ability to detect different types of splice sites.

#### Epigenetic Marks Prediction

Histone ChIP-seq data for ten histone modifications in the K562 human myelogenous leukemia cell line were retrieved from the ENCODE portal^82^. We downloaded the corresponding bed narrowPeak files with the following accession numbers:

- H3K4me3 (ENCFF706WUF)
- H3K27ac (ENCFF544LXB)
- H3K27me3 (ENCFF323WOT)
- H3K4me1 (ENCFF135ZLM)
- H3K36me3 (ENCFF561OUZ)
- H3K9me3 (ENCFF963GZJ)
- H3K9ac (ENCFF891CHI)
- H3K4me2 (ENCFF749KLQ)
- H4K20me1 (ENCFF909RKY)
- H2AFZ (ENCFF213OTI)

For each histone mark, we parsed all ENCODE-reported peak regions and extracted 1,024 bp genomic windows centred on the peak summits. We then unified the windows obtained from the ten marks: overlapping windows were merged, and the presence or absence of each mark within a given window was encoded as a binary indicator. The procedure yields a multi-label classification dataset, in which every 1 kb sequence may carry zero, one, or multiple histone-mark labels, providing a comprehensive benchmark for simultaneous prediction of diverse chromatin states.

#### CpG Methylation Prediction

In accordance with the data processing procedure in BEND^83^, we excluded CpG sites located on non-standard chromosomes or those with mismatched variants relative to the reference genome. Following DeepCpG^84^, CpG sites covered by fewer than 4 reads were excluded. CpG sites with at least 90% methylated reads were labeled as methylated, while those with less than 10% were labeled as unmethylated; remaining sites were discarded. The common subset of CpG sites passing these filtering criteria across all 7 experiments was retained, resulting in 959,039 sites. These sites were then extended with flanking context to form 512 bp windows centered on each CpG site. The resulting dataset was used to construct a 7-class multi-label classification task, with 74,309, 10,622, and 10,971 samples in the training, testing, and validation sets, respectively.

#### Regulation Element Prediction

Original data was sourced from Ensembl^85^, containing annotated genomic regions for “promoter”, “CTCF binding site”, “enhancer”, and “open chromatin region” in the human reference genome. To ensure a consistent input sequence length of 1,024 bp, regulatory elements exceeding this length were excluded, and the remaining elements were centered within their respective sequences. Based on this processed dataset, we designed two prediction tasks: a multi-label regulatory element prediction task, where each sequence could have multiple labels corresponding to the regulatory elements it contains, and a four-class classification task. In the latter, sequences with overlaps between different regulatory elements were excluded to maintain clarity and ensure that each sequence corresponded to a unique regulatory element category.

#### Species Classification

We selected six representative species, namely human (*homo sapien*), lemur (*lemur catta*), pig (*sus scrofa*), mouse (*mus musculus*), gorilla (*Gorilla gorilla*), and hippo (*hippopotamus amphibius*), and retrieved their complete genome assemblies together with the corresponding annotation files from the NCBI RefSeq database^86^. We constructed three data splits with minimal chromosomal overlap: the validation set comprises 5,000 sequences from chromosomes 1, 3, 12, and 13; the test set contains 5,000 sequences from chromosomes 5, 7, 9, 10, and 11; and the training set includes 40,000 sequences randomly sampled from all remaining chromosomes. Each split was subsequently stratified into sequence lengths of 512, 1,024, and 2,048 bp to systematically evaluate the effect of input length on predictive accuracy.

#### Regulatory Activity Prediction

Following DART-Eval^18^, we obtained the processed data as described in their evaluation protocol. This task predicts quantitative measurements of chromatin accessibility using DNase-Seq read counts, with data from the ENCODE consortium across five cell lines: GM12878, H1ESC, HEPG2, IMR90, and K562. Data preprocessing includes filtering peaks to remove outliers and expanding the remaining peaks to 2,114 bp, while also using matched negative genomic background sequences. Ground-truth activity is defined as the number of read endpoints that intersect the central 1,000 bp. The dataset is divided into train, validation, and test sets in an 8:1:1 ratio.

#### Enhancer Activity Prediction

For the enhancer activity prediction task, we constructed a dataset for a regression task, integrating data from two sources. First, we utilized data from DeepSTARR^87^, with the raw sequencing data available from the GEO database^88^ (GSE183939). DeepSTARR employs the UMI-STARR-seq to generate genome-wide high-resolution quantitative activity maps of enhancers in human HCT116 cells. In this dataset, 249 bp DNA sequences were mapped to their corresponding positions in the activity map, enabling the prediction of enhancer activity levels. Additionally, we incorporated data from the study^89^ on enhancers in HCT116 cells (GSE156741). This dataset provides information on the effects of eight cofactors (Brd2, Brd4, CDK7, CDK9, Med14, and p300CBP) on enhancer activity, along with control group (Parental) measurements. By integrating these two data sources, we constructed a comprehensive enhancer activity prediction dataset, offering a robust foundation for subsequent model evaluation.

#### Enhancer Promoter Interaction Prediction

Processed data was derived from TargetFinder^90^, where the authors constructed an enhancer promoter interaction dataset based on six human ENCODE cell lines (K562, GM12878, HeLa-S3, HUVEC, IMR90, and NHEK), integrating functional genomic annotations with high-resolution Hi-C chromatin interaction data. For each cell line, enhancer–promoter pairs exhibiting physical interactions identified from Hi-C experiments were labeled as positive samples, with genomic start and end coordinates provided for both enhancer and promoter regions. For each positive pair, 20 distance-matched negative pairs, i.e., pairs without supporting interaction evidence, were randomly sampled which formulates the problem as a binary classification task. This sampling strategy ensured a consistent distance distribution between positive and negative samples, allowing models to focus on regulatory characteristics rather than distance biases.

#### Gene Expression Prediction

Following the data-processing protocol in Genomics Long-range Benchmark^91^, we adopted the curated ExPecto dataset^92^, which provides comma-separated files containing representative transcription-start sites (TSS), strand information, and RNA-seq RPKM measurements for 218 GTEx tissues. In ExPecto, a CAGE peak is assigned to a GENCODE^81^ v24-annotated gene if it lies within ±1 000 bp of any annotated TSS. For each gene, the most abundant CAGE peak is chosen as the representative TSS; when no peak satisfies this criterion, the annotated gene start position is retained instead. We applied a log(1 + *x*) transformation to the raw RPKM values and subsequently standardised them (zero mean, unit variance) before model training.

Finally, sequence windows centred on each representative TSS were extracted from the GRCh37 human reference genome to serve as model inputs. To assess the impact of input length on predictive performance, we generated three window sizes per TSS, spanning 128, 256, 512, 1,024, and 2,048 base pairs, respectively.

#### Genetic Disease Classification

Original data was sourced from ClinVar^36^, which contains 13,209 disease types. Following the methodology of GV-Rep^93^, we filtered single nucleotide variants (SNVs) with a review status of at least one star, and located on the autosomes (chromosomes 1-22) and sex chromosomes (X, Y). For each variant, we extracted a 512 bp-long sequence centered on the variant by extending equally to both sides. Our primary focus was on diseases related to breast disease, cardiovascular disease, and kidney disease. For Breast Disease Type Classification, the diseases included Hereditary breast ovarian cancer syndrome, Familial cancer of breast, Breast-ovarian cancer, and others. The dataset for Cardiovascular Disorders Classification covered conditions such as Familial thoracic aortic aneurysm and aortic dissection, Dilated cardiomyopathy 1G, Hypertrophic cardiomyopathy, Long QT syndrome, Arrhythmogenic right ventricular dysplasia, Brugada syndrome, Primary dilated cardiomyopathy and Dilated cardiomyopathy 1DD. Finally, the Kidney Disease Type Classification task included diseases like Nephronophthisis, Autosomal recessive polycystic kidney disease, Renal cell carcinoma, Renal carnitine transport defect, Congenital adrenal hyperplasia due to cytochrome P450 oxidoreductase deficiency, Autosomal dominant polycystic kidney disease and Renal tubular dysgenesis.

#### Pathogenicity Classification

The data for the Pathogenicity Classification task was also sourced from ClinVar^36^, which contains annotations for various genetic variants and their associated pathogenicity classifications: likely benign, benign, pathogenic, and likely pathogenic. This constitutes a four-class classification problem. In line with the approach used in the Genetic Disease Classification task, the data was processed in the same manner. The final dataset for this task contains 47,121, 5,890, and 5,890 samples in the training, testing, and validation sets, respectively.

#### Splice Variant Effect Prediction

We utilized SpliceVarDB^94^, a database of experimentally validated splice variants in humans. To ensure data quality, we performed a series of preprocessing steps. First, we excluded variants labeled as “low-frequency” in SpliceVarDB to focus on more biologically relevant mutations. Next, we categorized the remaining variants into intronic and exonic mutations and extracted 1,024 bp-long sequence contexts centered on each variant. Finally, we split the dataset into train, validation, and test sets in an 8:1:1 ratio to ensure robust model evaluation and generalization. This task is formulated as a binary classification problem with two classes: Splice-altering and Normal.

#### BRCA1/2 SNV Functional Impact Classification

For all BRCA1 and BRCA2 SNVs with reported functional scores and classifications, we extracted sequence windows of 1,024 bp centered on the variant site from the human reference genome used in the respective original studies^40,41^. Additionally, we categorized the SNVs into coding and non-coding regions based on their genomic context. To ensure consistency in variant annotation, we adopted the classification scheme from the original studies: for BRCA1, SNVs labeled as “LOF” were classified as loss-of-function (LOF) variants, while those labeled as “FUNC” or “INT” were categorized as functional/intermediate variants; for BRCA2, SNVs labeled as “P strong”, “P moderate”, and “P supporting” were classified as loss-of-function (LOF) variants, whereas those labeled as “B strong”, “B moderate”, and “B supporting” were categorized as functional/intermediate variants. Based on these categorizations, we framed the task as a binary classification problem, with two classes: loss-of-function and functional/intermediate.

#### Mean Ribosome Loading Prediction

Mean Ribosome Loading Prediction is a task that predicts the translation efficiency of mRNA by estimating the ribosome loading on a given mRNA sequence, specifically focusing on the 5’ UTR. Ribosome loading refers to the number of ribosomes actively translating an mRNA at a given time. Based on the study^43^, a total of 91,519 random 5’ UTR sequences with lengths ranging from 25 to 100 were sampled, with 7600 human 5’ UTRs used for hold-out testing. To ensure the robustness and relevance of the data for human studies, a subset specifically tailored for human-focused research was created using Human7600^43^.

#### mRNA Stability Prediction

This task focuses on predicting the stability of mRNA. In a cell, steady-state gene expression levels can be easily affected by the mRNA stability. The mRNA stability dataset comprises 65,355 mRNA sequences with corresponding stability profiles, covering four vertebrate transcriptomes: mouse, human, frog, and fish. The coding genes in each transcriptome were collected from biomaRt^95^, and the mRNA stability for each gene was measured using iCodon^45^.

#### Abundance Prediction

Abundance Prediction aims to estimate transcript and protein abundance levels across multiple species using RNA-seq and proteomics data. These predictions are biologically significant because they help uncover fundamental relationships between gene sequences and their expression levels, which are crucial for understanding cellular processes, regulatory mechanisms, and protein production efficiency. In this task, transcript abundance prediction was performed on seven organisms: *Arabidopsis thaliana*, *Drosophila melanogaster*, *Escherichia coli*, *Homo sapiens*, *Saccharomyces cerevisiae*, *Haloferax volcanii*, and *Pichia pastoris*. Protein abundance prediction was limited to five of these species, as no protein abundance data was available for *H. volcanii* and *P. pastoris* in PaxDb. To build the dataset, RNA sequences were collected from the GEO database^88^, the EMBL-EBI Expression Atlas^96^, the primary literature^97^, the SRA database^98^, and PaxDb^99^. The sequences were clustered at 40% amino acid identity, and 7.5% were randomly sampled, stratified by RNA-seq-based protein abundance. Nucleotide BLAST was then used to remove homologous sequences (*≥*40% identity) from the training or validation set.

#### Non-coding RNA Classification

The Non-coding RNA Function Classification task focuses on classifying non-coding RNA (ncRNA) sequences into functional categories. ncRNAs are RNA molecules that do not encode proteins but play critical roles in various cellular processes such as regulation of gene expression, RNA splicing, and chromatin remodeling. The identification and classification of ncRNA types and their functions are essential for understanding their involvement in cellular processes and disease mechanisms. This task helps uncover the regulatory roles of different ncRNA classes, such as miRNAs, lncRNAs, and siRNAs, in cellular functions. Based on the study^100^, we extracted 8,920 small ncRNA sequences from the Rfam database^101^, spanning 13 classes (miRNA, 5S rRNA, 5.8S rRNA, ribozymes, CD-box, HACA-box, scaRNA, tRNA, Group I introns, Group II introns, IRES, leader, riboswitch) that represent a subset of the database. To reduce redundancy, CD-HIT tool^102^ was applied to retain 20% non-redundant sequences.

#### Expression Level and Translation Efficiency Regulation Prediction

In eukaryotic cells, where transcription and translation are uncoupled, protein expression levels depend not only on mRNA expression level (EL), which is governed by the transcription machinery, but also on translation efficiency (TE) governed by the translational machinery. To investigate this regulatory interplay, the tasks aim to predict both EL and TE from endogenous datasets using 5’ UTR sequences as inputs. This uncovers fundamental principles of translational regulation, especially in eukaryotic cells. Specifically, we utilized the endogenous human 5’ UTR data^50^, which integrates matched RNA-Seq and Ribo-Seq datasets from three distinct cell lines/tissues: Human Embryonic Kidney 293T (HEK), Human Prostate Cancer Cells (PC3), and Human Muscle Tissue (Muscle). The data includes 14,410 sequences from HEK cells, 12,579 from PC3 cells, and 1,257 from muscle tissue. For each 5’ UTR, the mRNA EL was determined using RNA-seq RPKM, while TE was calculated by dividing Ribo-seq RPKM by RNA-seq RPKM, where RPKM represents Reads Per Kilobase of transcript per Million mapped reads. Following the method described in previous work^103^, a fixed 100-bp 5’ UTR sequence upstream of the coding sequence (CDS) was chosen. Transcripts with RNA-seq RPKM <5 and Ribo-seq RPKM <0.1 were excluded. This resulted in six distinct sub-datasets, corresponding to EL and TE prediction tasks across the three cell types.

#### Modification Prediction

Modification Prediction is a task focused on predicting the types of RNA modifications that occur on a precursor mRNA (pre-mRNA) sequence. RNA modifications, such as m6A, m1A, m5C, and others, are post-transcriptional chemical changes that occur after the RNA is transcribed but before it matures. These modifications increase the structural and functional diversity of RNA molecules and regulate all stages of RNA life. Precise identification of RNA modification sites is therefore important to understanding the functions and regulatory mechanisms of various RNAs. Following the previous work^104^, we framed this as a 12-class multi-label classification task, where each site may have one or more of the 12 distinct RNA modification types (*e.g.*, m6A, m1A, m5C). The corresponding RNA sequences were collected from the GEO database^88^, RMBase^105^, and the RADAR database^106^. Negative sites were randomly sampled from unmodified bases within the same transcripts as the positive sites. After preprocessing, we compiled a dataset comprising over 300k modification sites.

#### APA Isoform Prediction

Alternative Polyadenylation (APA) Isoform Prediction is a task focused on predicting the usage ratio of proximal polyadenylation (polyA) signals (PAS) in the 3’ UTR of mRNA. In the APA process, genes can generate multiple mRNA isoforms by using different PAS, which leads to variations in the length of the 3’ UTR. These isoforms can have distinct functional roles, influencing mRNA stability, translation efficiency, and interactions with other regulatory molecules. Following the preprocessing steps in BEACON^22^, 228k sequences were filtered from more than 3 million APA reporter isoform expression data from APARENT^51^.

#### SARS-CoV-2 Vaccine Degradation Prediction

The SARS-CoV-2 Vaccine Degradation task focuses on predicting the hydrolysis rates of mRNA by using RNA sequences provided by Eterna players. We collected the dataset from OpenVaccine^52^, which contains RNA degradation profiles for the first 68 nucleotides. These profiles were measured using In-line-seq, a high-throughput method designed to characterize RNA degradation through in-line hydrolysis. The average of the deg_Mg_50C values at each nucleotide is treated as the sequence-level target, since it has the highest correlation with other degradation profiles, including deg_pH10, deg_Mg_pH10, and deg_50C. This task is crucial for addressing mRNA vaccine instability and designing stabilized RNA therapeutics. The instability of RNA molecules, particularly due to in-line hydrolysis—which is the spontaneous chemical cleavage of the phosphodiester backbone—presents significant challenges for the storage, distribution, and efficacy of mRNA vaccines.

#### Programmable RNA Switches Prediction

Programmable RNA switches, such as toehold switches, represent a pivotal class of synthetic biology tools that enable precise detection and response to molecular signals (*e.g.*, nucleic acids, proteins, or small molecules). These engineered RNA elements function as modular, sequence-specific biosensors, with applications ranging from genetic circuit design to diagnostic platforms.

To analyse and decode the complex sequence-function relationships of these RNA switches, this task aims to predict the ON/OFF states activity of programmable RNA switches using the toehold-switch library in previous study^53^, which includes 91,534 sequences spanning across the complete genomes of 23 pathogenic viruses and the entire coding regions of 906 human transcription factors. The ON and OFF signals, which characterize a given switch, were obtained by GFP signal intensity measurements and normalized from 0 to 1.

#### Tc-Riboswitches Prediction

Riboswitches are RNA-based regulatory elements that control gene expression by undergoing conformational changes upon ligand binding, typically located in untranslated mRNA regions. The tetracycline (Tc) riboswitch specifically uses a high-affinity Tc aptamer to translationally repress genes when bound to tetracycline. This task focuses on predicting the dynamic range of Tc riboswitches—the ratio of gene expression in the “on-state” (without ligand) to the “off-state” (with ligand)—based on their upstream RNA sequences. The dataset consists of 355 Tc-riboswitch dimer sequences positioned upstream of a green fluorescent protein (GFP) reporter gene, as curated from previous study^54^. Each sequence is labeled with a switching factor, reflecting the difference in GFP fluorescence with and without Tc, as measured by flow cytometry.

#### mRFP Expression Prediction

The mRFP Expression task focuses on predicting the protein production levels of monomeric red fluorescent protein (mRFP) based on variations in its coding sequence. To construct the dataset, mRFP gene libraries were generated using type IIS restriction and ligation methods, resulting in 1,459 unique gene variants that all encode the same protein but differ in their synonymous codon usage^107^. Codon usage across almost the whole gene was fully randomized to enable systematic exploration of how synonymous changes affect translation. Each of the 1,459 gene variants was expressed in *Escherichia coli*, and its corresponding protein production level was experimentally quantified following the protocol in previous work^107^. The resulting dataset provides a continuous expression value for each sequence, framing the task as a regression problem. By learning the relationship between codon usage and protein production, this task offers insights into how synonymous sequence variation influences gene expression efficiency.

#### Secondary Structure Analysis

Secondary structure elements are fundamental to protein folding and function, with *α*-helices and *β* -strands forming the basis of complex protein architectures^57^. We obtained protein sequences and secondary structure annotations from the UniProt database^108^, mapping Ensembl transcript IDs to their corresponding UniProt accessions. From these linked annotations, we extracted helix and strand data to classify each protein into three structural categories: mainly helix (helix ratio >0.6), mainly strand (strand ratio >0.6), or mixed structure.

Only valid structure elements were considered, requiring helices to be at least 3 residues long and strands to be at least 2 residues long. Duplicate protein sequences were removed to prevent data leakage. Both protein sequences and their corresponding coding DNA sequences were collected. The dataset was stratified and split into train, validation, and test sets in a 64:16:20 ratio.

#### Transmembrane Region Prediction

Transmembrane proteins are essential components of cell membranes, functioning in cellular signal transduction, substance transport, and ion channels, with approximately 60% of current drug targets being transmembrane proteins^58^. Protein sequences with transmembrane annotations were sourced from UniProt^108^. Transmembrane feature information was extracted by mapping Ensembl transcript IDs to their corresponding UniProt accessions. Only transmembrane regions with a minimum length of 15 residues were considered valid. We classified proteins into six categories based on the number of transmembrane segments: single-pass (1 region), double-pass (2 regions), triple/quad-pass (3-4 regions), multi-pass type 1 (5-6 regions), multi-pass type 2 (7-8 regions), and multi-pass type 3 (*>* 8 regions). Both protein sequences and their corresponding coding DNA sequences were collected. The dataset was stratified and split into train, validation, and test sets in a 64:16:20 ratio.

#### Protein Variant Classification

Accurate classification of protein variants has applications in understanding human genetic diseases^59^, assisting clinicians in genetic diagnosis, and providing insights for personalized medicine. Protein sequences with variant annotations were collected from UniProt^108^. Variant information was extracted through mapping between Ensembl transcript IDs and UniProt accessions. Proteins were classified into two groups based on variant type: missense-enriched (proteins with at least 2 missense variants and more than twice as many missense as silent variants) and mixed variants (all other variant patterns). We also classified proteins by variant density: low (*≤* 3 variants), medium (4-8 variants), and high (*>* 8 variants). Both protein sequences and their corresponding coding DNA sequences were collected. The dataset was stratified and split into train, validation, and test sets in a 64:16:20 ratio.

#### Post-translational Modifications Prediction

post-translational modifications (PTMs) are crucial regulators of protein function^109^, affecting protein-protein interactions, stability, and cellular localization. Protein sequences and PTM annotations were sourced from UniProt^108^. PTM information was obtained by linking Ensembl transcript IDs to UniProt accessions. We classified PTMs into three main categories: proteins with predominantly phosphorylation sites (*>* 50% of total modifications), proteins with predominantly glycosylation sites (*>* 50% of total modifications), and proteins with other specific PTMs or mixed modification patterns. PTM types were identified by keyword analysis of the annotation descriptions. Both protein sequences and their corresponding coding DNA sequences were collected. The dataset was stratified and split into train, validation, and test sets in a 64:16:20 ratio.

#### Protein Domain Classification

This classification task helps understand protein interaction networks, as proteins with complementary rather than identical domains are more likely to interact functionally, even when their sequences are highly similar^61^. Protein sequences with domain annotations were derived from UniProt^108^. Ensembl transcript IDs were linked to UniProt accessions to access protein domain annotations. For domain selection, we implemented a minimum frequency threshold of 30 occurrences and focused on proteins containing single domains to reduce classification ambiguity. The most frequent domains (KRAB, Protein kinase, BTB, and others) were selected as classification targets, with numerical labels assigned to each category (0-3). Duplicated protein sequences were removed to prevent data leakage. Both protein sequences and their corresponding coding DNA sequences (CDS) were collected. The dataset was stratified and split into train, validation, and test sets in a 64:16:20 ratio. Protein domains represent conserved structural and functional units that serve as basic interaction modules.

#### Enzyme Classification

We obtained protein sequences and their enzyme annotations from the UniProt database^108^. We mapped Ensembl transcript IDs to UniProt accessions and linked these to enzyme classification annotations. We classified enzymes based on the EC number system, focusing on the six main enzyme classes: oxidoreductases (EC 1), transferases (EC 2), hydrolases (EC 3), lyases (EC 4), isomerases (EC 5), and ligases (EC 6). For proteins with multiple EC numbers, we assigned the predominant enzyme class as the label. To balance the dataset, we grouped less frequent classes (lyases, isomerases, and ligases) into a single category, resulting in four classification targets (classes 1-3 and others). Duplicate protein sequences were removed to prevent data leakage. Both protein sequences and their corresponding coding DNA sequences were collected. The dataset was stratified and split into train, validation, and test sets in a 64:16:20 ratio. Enzymes are biological catalysts that accelerate chemical reactions in living organisms, and accurate classification supports understanding protein functional mechanisms, drug development, and disease treatment.

#### Beta-Lactamase Activity Prediction

The Beta-Lactamase Activity task explores the fitness landscape of single codon substitutions in the TEM-1 gene. Following^23^, we adopted the dataset derived from^64^. The data measures a mutant gene’s ability to confer ampicillin resistance, with fitness values calculated as a weighted average of allele counts across multiple ampicillin concentrations. The dataset includes both nucleotide sequences from the original study and their corresponding amino acid translations. The dataset is provided in two versions: a “Complete” set containing all CDS samples except those degenerate with respect to test set sequences, and a “Unique” set with a random maximal subset of non-degenerate coding sequences.

#### Fluorescence Prediction

The Fluorescence Prediction task evaluates the ability to predict log-fluorescence of higher-order mutant green fluorescent protein (avGFP) sequences. We collected data from^110^, which maps the avGFP fitness landscape through random mutagenesis in E. coli. Following^111^, the dataset has the training set restricted to amino acid sequences with three or fewer mutations from parent GFP sequences, while the test set contains sequences with four or more mutations. Unlike the Beta-lactamase dataset, only a small fraction of sequences were degenerate due to the less systematic mutagenesis process. From the original 58,417 total sequences, 54,025 uniquely translating coding DNA sequences were identified in the training and validation sets. The dataset includes both nucleotide sequences and their corresponding amino acid translations.

#### Melting Point Prediction

The Melting Point Prediction task evaluates a model’s ability to predict protein melting temperatures in a sequence-level regression framework. We followed^112^, whose dataset was compiled using a mass spectrometry-based proteomic approach. Following the methodology in^113^, we use the “mixed” splitting strategy where sequences are clustered at 20% identity with 80% of clusters allocated to the training dataset and the remaining 20% assigned to the test dataset. This approach helps avoid over-representation of large sequence clusters.

#### Protein Stability Prediction

The Protein Stability Prediction task evaluates the ability to predict stability around a small region of high-fitness sequences. We derived dataset from^65^. The training and validation sets contain a broad selection of de novo computationally designed proteins composing a small number of topologies, while the test set consists of the neighborhoods of single codon mutations around a few of the most stable candidates. Stability is measured as a function of resistance to increasing levels of protease, defined as the difference between the EC50 value of the protein and that of the predicted EC50 in the unfolded state, calculated on a log 10 scale.

### Experimental Settings

#### Finetuning Details

We initialize each gLM with its pretrained checkpoint and append a task-specific head—either a single-layer regression head for continuous targets or a softmax classification head for categorical outcomes. During finetuning, we allow all parameters of the base model and the added head to be updated so as to fully adapt to each task. Input representations from the base model are passed through this head to produce predictions for each dataset. In addition to the gLMs, we include two baseline architectures: a CNN and a LSTM. Both baselines were trained from scratch on each dataset using the same finetuning protocol as the gLMs. We employ mean squared error loss for regression tasks and cross-entropy loss for classification tasks, ensuring that the objective aligns with the nature of each target. For optimization, we utilized the AdamW optimizer with a cosine learning rate scheduler. To ensure a fair comparison, we conducted an independent learning rate search for each model to ensure that its true performance potential was realized. For each model and dataset, we choose the epoch with the minimum validation loss to evaluate performance on the test set. All reported evaluation metrics were computed on the test set. All reported evaluation metrics were computed on the test set.

#### Evaluation Metrics

We comprehensively evaluate the performance of various models across the aforementioned tasks using a diverse set of assessment metrics. These metrics are carefully adapted to accommodate the specific characteristics of each task.

**Accuracy.** The Accuracy metric measures the proportion of correctly classified instances out of all predictions made, as given by the following formula:

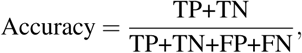

where TP,TN,FP,FN represent True Positives, True Negatives, False Positives, False Negatives, respectively.

**F1 Score.** However, in cases where class imbalance is prevalent, Accuracy alone may not provide a complete picture of model performance, as a model might achieve high accuracy by favoring the majority class. To address such issues, the F1 score is used to provide a more nuanced evaluation. The F1 score is the harmonic mean of Precision and Recall, balancing the trade-off between the two. It is computed as:

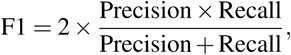

where Precision 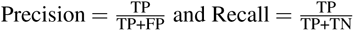.

**Macro-F1 Score.** Macro-F1 is computed by averaging the F1 scores of each class independently, thereby assigning equal weight to all classes regardless of their frequency:

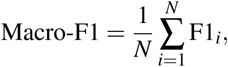

where *N* is the number of classes, and F1*_i_* is the F1 score of class *i*. Macro-F1 score is essential for a comprehensive assessment of model performance, particularly in scenarios where class distribution is skewed.

**Coefficient of Determination** *R*^2^. *R*^2^ is commonly used to assess the goodness-of-fit in regression models. It indicates the proportion of the variance in the dependent variable that is predictable from the independent variables:

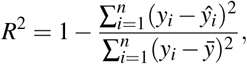

where *y_i_*are the true values, *y*^*_i_* are the predicted values, and *ŷ* is the mean of the true values.

**Spearman’s Rank Correlation Coefficient** *ρ*. is a non-parametric metric that captures the strength and direction of monotonic relationships between predicted and actual values. This metric is particularly robust to outliers and non-linear associations, making it suitable for measuring ranking consistency in various prediction tasks:

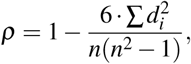

where *d_i_*is the difference between the ranks of the predicted and true values for each observation, and *n* is the total number of observations.

#### Computing Software and Hardware

We conducted all experiments and analyses in this study using Python (v3.8.19) and PyTorch (v2.2.0, CUDA 12.1) (https://pytorch.org) unless stated otherwise. For each gLM, we employed the official pretrained checkpoint, with the corresponding download links provided in Table 1. Models with fewer than 500 million parameters were finetuned on a single NVIDIA RTX 3090 GPU. Models with more than 500 million parameters were finetuned on a single NVIDIA H800 GPU.

## Data availability

A complete list of the source databases used is provided in Table 39.

**Table 39.**
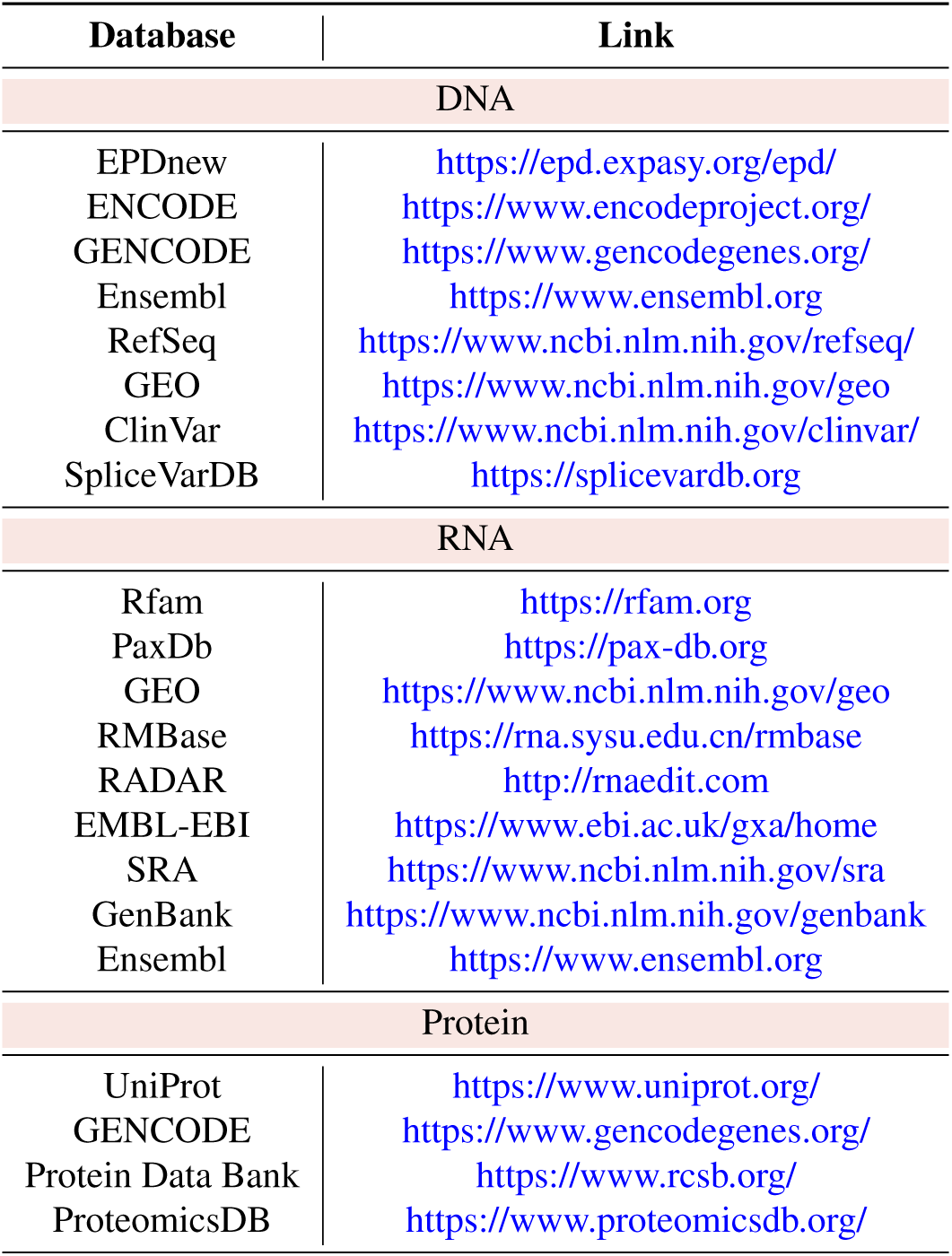
List of databases used in the Genomic Touchstone.

## Code availability

A comprehensive list of the models and pre-trained weights evaluated in this study is provided in Table 1.

## Ethics declarations

This study was approved by the Institutional Review Board under protocol number HREP-2024-0269. All data collection and analyses were conducted in accordance with the approved ethical guidelines and regulations.

## Author contributions

Y.W., Z.C., and H.C. conceived and designed the work. Y.W. contributed to the technical implementation and conducted experiments. Y.W., Z.C., Q.Z., Y.G., and J.O. participated in discussions regarding the design of the experimental evaluations, model selection, and data usage. Y.W., Q.Z., Y.G., and J.O. collected and preprocessed the data used in the benchmark. Y.X, S.Y., S.H., Y.N., Y.C., F.Z. and C.J. offered insightful suggestions for the experimental design. Z.X., D.Z., T.X., C.Y., X.F., J.W., K.Z., J.Y. and R.R. provided critical biological insights and guidance. Y.W., Z.C., Q.Z., Y.G., and J.O. contributed to the drafting of the manuscript. X.W., Z.X, K.T.C., J.W., X.F. and R.R. contributed to revising of the manuscript. H.C. supervised the research.

## Declarations

J.Y. was employed by Tencent AI Lab when this work was done. The company did not influence the research. The other authors declare no competing interests.

## Acknowledgements

This work was supported by the Hong Kong Innovation and Technology Commission (Project No. GHP/006/22GD and ITCPD/17-9), and the Research Grants Council of the Hong Kong Special Administrative Region, China (Project No. R6003-22 and T45-401/22-N).

